# Modeling the START transition in the budding yeast cell cycle

**DOI:** 10.1101/2023.11.05.564806

**Authors:** Janani Ravi, Kewalin Samart, Jason Zwolak

## Abstract

Budding yeast, *Saccharomyces cerevisiae*, is widely used as a model organism to study the genetics underlying eukaryotic cellular processes and growth critical to cancer development, such as cell division and cell cycle progression. The budding yeast cell cycle is also one of the best-studied dynamical systems owing to its thoroughly resolved genetics. However, the dynamics underlying the crucial cell cycle decision point called the START transition, at which the cell commits to a new round of DNA replication and cell division, are under-studied. The START machinery involves a central cyclin-dependent kinase; cyclins responsible for starting the transition, bud formation, and initiating DNA synthesis; and their transcriptional regulators. However, evidence has shown that the mechanism is more complicated than a simple irreversible transition switch. Activating a key transcription regulator SBF requires the phosphorylation of its inhibitor, Whi5, or an SBF/MBF monomeric component, Swi6, but not necessarily both. Also, the timing and mechanism of the inhibitor Whi5’s nuclear export, while important, are not critical for the timing and execution of START. Therefore, there is a need for a consolidated model for the budding yeast START transition, reconciling all known regulatory and spatial dynamics. We built a detailed mathematical model (START-BYCC) for the START transition in the budding yeast cell cycle based on all established molecular interactions and experimental phenotypes. START-BYCC recapitulates the underlying dynamics and correctly emulates key phenotypic traits of ∼150 known START mutants, including regulation of size control, localization of inhibitor/transcription factor complexes, and the nutritional effects on size control. Such a detailed mechanistic understanding of the underlying dynamics gets us closer towards deconvoluting the aberrant cellular development in cancer. All wildtype and mutant simulations of our START-BYCC model are available at sbmlsimulator.org/simulator/by-start, and the supporting data is available on GitHub: github.com/jravilab/start-bycc.

## Introduction

### Background and Significance

The cell cycle underlies all biological growth, reproduction, development, and repair, and its misregulation results in complex human diseases, such as cancer. Genetic and molecular research over the past several decades has provided a rigorous understanding of the molecular mechanism coordinating the cell cycle^1–4^. The cell cycle growth-and-division process is characterized by alternating DNA replication (S phase) and mitosis (M phase), interspersed by temporal gap phases G1 and G2 to ensure balanced growth and division. Several checkpoints occur throughout the cell cycle progression and control the commitment to DNA replication, proper alignment, and segregation of chromosomes and mitosis.

The molecular machinery of the cell cycle is highly conserved across eukaryotes^5^, and much of the regulatory system has been systematically worked out in the budding yeast (*Saccharomyces cerevisiae*). Budding yeast is an especially attractive model organism for study of the cell cycle, largely because it is genetically tractable and can exist as a haploid. In eukaryotes, cyclin-dependent kinases (CDKs) play a key role in initiating crucial cell cycle events by phosphorylating specific protein targets. CDK levels remain constant throughout the cell cycle, but their activity and substrate specificity are governed by their binding partners, cyclins, whose concentrations fluctuate throughout the cell cycle. The cyclin/CDK complexes are tightly regulated by synthesis, degradation, and/or sequestration by the cyclin-dependent kinase inhibitor (CKI) complexes. The budding yeast presents a less complex cell cycle circuitry than mammalian counterparts, with only one CDK (Cdc28)^1^ that binds to one of nine cyclins of type Cln or Clb. The most important cyclins involved in the different phases of a budding yeast cell cycle are Cln3 (G1), Cln1,2 (G1/S), Clb5,6 (S), Clb3,4 (early M), and Clb1,2 (late M) (**Table S1**). The names of the cyclin-binding partners during different phases of the cell cycle are used to refer to the relevant heterodimer.

The yeast cell division process begins with the cell budding and then dividing asymmetrically at the neck to produce a large mother and a small daughter cell, each with a set of sister chromatids. Soon, the mother cell repeats the process; in contrast, the daughter cell has a long G1 phase before producing the first bud and entering the S phase. This characteristic commitment step in the budding yeast cell cycle, including bud initiation, the onset of DNA synthesis, and spindle pole body duplication, is referred to as ‘START’^6^.

Classic studies on ‘size control’ in budding yeast have shown that small newborn daughter cells in G1 have longer lag periods than larger newborn cells before the START transition (first appearance of bud), even if they are genetically identical and placed under identical physiological conditions^7^. This delay allows the small cells to grow large enough to attain a minimum size threshold before their START transition and commitment to the S phase. Notably, this size threshold depends on the nutrient medium, with the threshold increasing in proportion to the richness of the medium^8–11^. Cells passing through the START transition also abruptly lose their response to pheromones (mating factors), which are inhibitors of the cell cycle^12–14^ and to which cells are sensitive in the G1 phase. Contingent on favorable external (nutrients, pheromones) and internal (DNA damage) cues, START is the point of commitment towards coordinated DNA replication and cell division^10,15^.

### Key dynamical processes modeled

The most important aspects underlying the molecular mechanisms for the budding yeast cell cycle have been studied in great detail^16–18^. We previously built mathematical models of the budding yeast cell cycle^19,20^ based on these extensive molecular studies. Here, we build upon a detailed mechanistic cell cycle model by supplementing the most relevant findings in START.

#### START transition

In early G1, only Cln3 is available, and its level increases with cell size^2^. When the cell attains a critical size, Cln3 activates two transcription factors: SBF, a heterodimer of Swi4 and Swi6^21^, and MBF, a heterodimer of Mbp1 and Swi6^22^. In the absence of Cln3, Bck2 plays a role in the activation of SBF and MBF in response to cell size^21,23^. SBF and MBF drive the irreversible START transition (except under starvation^24^) by activating the transcription of G1 cyclins Cln1,2, and S phase cyclins Clb5,6^21,23,25^. SBF and MBF have a large functional overlap^26^. In the absence of SBF, MBF can activate Cln1,2. Likewise, SBF can activate Clb5,6 in the absence of MBF.

Clb5,6 are not active initially in late G1 due to the presence of their stoichiometric CDK inhibitor, CKI (Sic1 and Cdc6)^27^. Active Cln1,2 phosphorylate Sic1 (and Cdc6), signaling it for rapid degradation^28^, resulting in Clb5,6 activation soon after Cln1,2 activation. At START transition, as SBF and MBF are activated, Cln1,2 accumulate first, leading to bud emergence, followed closely by Clb5,6 activation, leading to the initiation of DNA synthesis (**Fig. 1A**). In wildtype cells, these two events occur almost simultaneously – a characteristic of START transition^6^.

**Figure 1.**
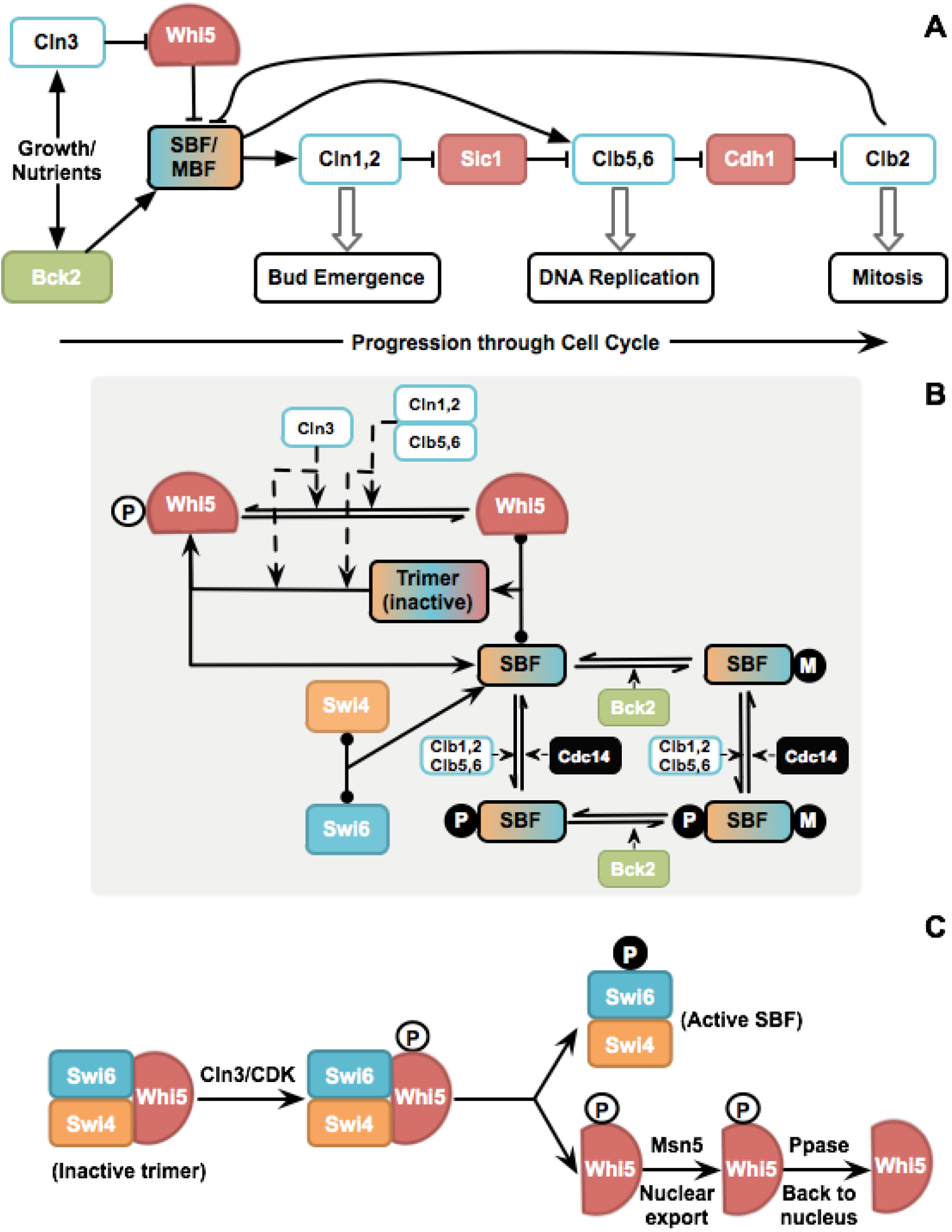
Simplified mechanism of cell cycle progression and START transition. (A) Progression through the cell cycle in budding yeast. Cln3 and Bck2 are activators of START (turn on SBF, MBF needed for Cln1,2; Clb5,6). Cln1,2 phosphorylates and inhibits Sic1, a stoichiometric inhibitor of Clb5,6, thus allowing DNA replication to occur. S-phase cyclins, Clb5,6, inhibit Cdh1, an antagonist of the mitotic cyclins, Clb1,2, thus allowing progression through the mitotic events, and finally exit from mitosis leading back to G1. (B) The earlier hypothesis that Whi5 phosphorylation is crucial for SBF activation. This model for START includes (i) activation of SBF by Clns (Cln3, Cln1,2, Clb5,6) by inactivation of Whi5 (by phosphorylating free and SBF-bound Whi5) in late G1, (ii) activation of SBF by Bck2 to an alternate form independent of Whi5 and CDK, and (iii) inactivation of SBF by Clb1,2 in late S phase. (C) Earlier hypothesis on association, dissociation, and translocation events underlying SBF activation. Inactive SBF-Whi5 trimer gets phosphorylated by Cln/CDK on both Swi6 and Whi5, followed by the trimer dissociation to give active SBF (with Swi6 phosphorylated) and phosphorylated Whi5 that can get exported to the cytoplasm (by transport protein, Msn5).

#### G2/M transition

Clb5,6 activated at START inhibits Cdh1, a protein that degrades mitotic cyclins Clb1,2^29^. Thus, Clb1,2 accumulate when their synthesis is turned on later in the G2 phase, followed by entry into mitosis (**Fig. 1A**). Thus, the START transition also facilitates the progression of the cell cycle to the G2/M phase^30^. At the end of the S phase, SBF is turned off by Clb1,2^31,32^, and MBF by Clb1,2 and Nrm1^33^.

#### FINISH transition

At the end of mitosis, when all sister chromosome pairs are attached to opposite poles of the spindle, Cdc20 is activated^34^. Cdc20, along with the anaphase-promoting complex (APC), causes the dissolution of the cohesin that holds the sister chromatids together, allowing them to move apart to the opposite poles of the spindle. The dissolution of chromatids triggers the activation of a phosphatase, Cdc14, known for three key roles: (i) activation of CKI synthesis^35^, (ii) dephosphorylation and stabilization of CKI^36^, and (iii) activation of Cdh1 with the help of Cdc20, leading to the degradation of Clb1,2^37–39^. Degradation of mitotic cyclins enables the cell to exit from mitosis, resetting the cell for the next cycle^40^. The details of the metaphase-to-anaphase transition involving other critical proteins such as Esp1, Pds1, MET, and other proteins are beyond the scope of this model, but are incorporated in detail in Hancioglu and Tyson 2012^41^. In summary, at FINISH, the cell exits mitosis and returns to G1 if DNA is fully replicated and undamaged, with chromosomes aligned perfectly at the metaphase plate, ready to be separated into daughter cells^42^. The mechanistic details underlying the FINISH transition have been studied and addressed by various mathematical models^41,77–79^ including the latest developed model combining the deletion of other genetic elements of the FINISH transition with Clb2 overexpression to predict the presence or absence of Cdc14 endocycles in mutant strains^46^.

#### START vs. R-point

Activation of SBF by Cln3 is central to START, and prior studies suggest that a double-repression mechanism underlies this activation (**Fig. 1, A–B**). SBF is bound and inactivated by the inhibitor, Whi5^47,48^. Cln3 to (Cln3/Cdc28) phosphorylates Whi5, which causes its dissociation from SBF, rendering SBF active (**Fig. 1B**). Bck2, another known activator of START, acts by a mechanism independent of CDK and Whi5^49^. In the late S phase, SBF is phosphorylated and turned off by Clb2^31,32^ and Clb6^50^ (**Fig. 1B**). It is noteworthy that the Cln3-Whi5-SBF (cyclin-inhibitor–transcription factor) circuit that operates at the START transition in budding yeast is thus reminiscent of the CycD-pRb-E2F system operating at the restriction point (R-point) in mammals, albeit with different dynamics: [Cln3 –| Whi5 –| SBF → Cln1,2 → START] vs. [CycD –| pRb –| E2F → CycE → R-point]^51^. The inhibitor pRB is a known tumor suppressor, and genes resulting in cyclin activation are known oncogenes^52–55^, with pRB phosphorylation playing a key role in G1/S transition^56^.

#### Whi5 phosphorylation

Due to the parallels between the mammalian R-point and yeast START systems, the prevailing notion was that the timely activation of START solely depends on the phosphorylation of Whi5 and resulting SBF activation (**Fig. 1C**). As cells progress through START, Whi5 gets progressively phosphorylated and inactivated^57^. Consequently, it becomes cytoplasmic in late G1, leaving SBF in the active state for Cln1,2 transcription. Whi5 moves back to the nucleus to inhibit SBF only at mitotic exit. Phosphorylation was also thought to play a role in the nuclear export of Swi6 in the late S phase concurrent with SBF inactivation^50,58^. These observations together made up the simple story of the activation/inactivation of SBF and the relocalization of the inhibitor (**Fig. 1C**). Accordingly, if the phosphorylation of Whi5 were key to its export and SBF activation, then mutating all the phosphorylation sites (to alanine) should retain Whi5 in the nucleus, delaying SBF activation, and hence resulting in larger cells. Likewise, mutation of the phosphorylation sites on Swi6 should cause Swi6 retention in the nucleus, thereby advancing START and resulting in smaller cells.

The study by Wagner et al.^59^, however, challenged both of these expectations. *WHI5-12A*, a non-phosphorylable mutant with all known CDK (and non-CDK sites) phosphorylation sites in Whi5 mutated to alanine, showed no difference in size compared to wildtype^59^. *SWI6-SA4*, a mutant with non-phosphorylable Swi6, was also observed to have a wildtype size^58,60^. Only the double non-phosphorylable mutant *WHI5-12A SWI6-SA4* showed a 40% increase in size^59^. Together, these experiments indicate that either Whi5 or Swi6 needs to be phosphorylated (if not both) for timely activation of SBF (so that there is no difference in size). These observations present a significant departure in the START transition process from that of the mammalian restriction point. While Whi5 phosphorylation is not necessary for the activation of SBF in budding yeast, pRb phosphorylation has been shown to be absolutely crucial for the activation of E2F in mammalian cells^47,61^.

#### Modeling the yeast cell cycle

Mathematical modeling has proven to be an invaluable tool for understanding the workings of cellular regulatory systems such as the cell cycle^42,62,63^. The budding yeast cell cycle has been studied and is understood in great detail using the deterministic mathematical models published by Chen et al.,^19,20^ (using ordinary differential equations). The 2004 model^20^ (henceforth referred to as the BYCC model), in particular, incorporates several aspects of the cell cycle mechanism from START transition through mitotic exit and can explain the phenotypes of ∼120 cell cycle mutants. The model, however, contains a very simplistic depiction of START: SBF/MBF activation by Cln3 and Bck2, and inactivation by Clb2 in a condensed phenomenological abstraction (using an ultra-sensitive switch^64^). It does not include several of the more recently discovered details pertaining to START discussed above.

Subsequently, we extended the BYCC model to incorporate the role of Whi5, the relevant localization between nucleus and cytoplasm, enabling deeper insights into the critical cell size required for the G1/S transition and budding^65^. A few recent models from our group and others have included more mechanistic details for cell size control and viability^66–69^, including the separation of Clb5,6 into Clb5, Clb6 and emphasizing the positive feedback loops in the START transition^66^. Genome-scale models have applied reaction-contingency language^70^ while we also included aspects of START and FINISH modules^42,46^. Concurrently, we also built alternate and abstracted START models with a standard component model (SCM) and boolean approaches leveraging continuous, discrete, and stochastic methods^71,72^. Other studies have focused on the cell fate decision at G1 involving the mating factor Far1^73^ and the feedforward regulation-mediated loop^74^. Additionally, a recent study discovered that the START transition could be reversed under starvation by interrupting the positive feedback loop that activates the G1/S transition, resulting in re-importing Whi5 to the nucleus to inhibit SBF^24^. In summary, while we and others have developed and derived insights from several kinds of models representing various stages of the cell cycle, we still lack a detailed understanding of the molecular mechanisms underlying START dynamics.

In light of all the findings of the role of Whi5 phosphorylation and their discrepancies with the previously held consensus mechanisms, there is a need to reconcile a coherent model for the START transition in budding yeast that recapitulates the START dynamics while staying consistent with the observed genetics and mutants. Such a model would offer broad insights into the study of cell cycle dynamics and molecular mechanisms, as well as address specific questions about the dynamics underlying the BYCC START transition. In this paper, we have developed a comprehensive nonlinear ordinary differential equation mathematical model for START, with the following features: i) a detailed mechanism for the activation and inactivation of SBF, along with the inhibitor Whi5, that is compliant with the experimentally determined phenotypes^47,65^, ii) a mechanism for activation and inactivation of MBF^21,33,48^, iii) highlighting the role of Bck2 in the START transition, iv) critical aspects of the localization of the monomers in the transcription-factor/inhibitor complex (Whi5, Swi6, and Swi4)^57,58,75^, and, v) size control operating at the START transition^8,65^, and we have also taken into account the specific contribution of Cln3 in setting the cell size threshold^76^.

Our new model for START has been integrated with the full cell cycle model^20^ (BYCC), resulting in START-BYCC. Our current model (START-BYCC) explains wildtype cell-cycle dynamics, timely localization of monomers, and size control, as well as experimentally observed phenotypes of over ∼120 START mutants, few of which have been experimentally corroborated^46^.

## Results and Discussion

We used the detailed BYCC mathematical model^20^ as a starting point and built a detailed model for START transition upon it by reconciling several recent studies (**Table S2; Fig. 2, 3**). The main focus of START-BYCC is the description of the activation and inactivation of G1/S transcription factors, SBF and MBF. At its core, the model contains the SBF and MBF monomers Swi4, Swi6, and Mbp1, and the inhibitor protein, Whi5. Additionally, we consider two distinct promoters for the two sets of genes involved in budding and DNA synthesis that are turned on by SBF and MBF, respectively. We take into account all pools of monomers and protein complexes that are either bound or unbound to the promoter. We also incorporate all the pertinent phosphorylation states for each of these components based on whether they are Cln/CDK or Clb/CDK targets. Further, we include any available information on the cellular localization of these molecules during different phases of the cell cycle.

**Figure 2.**
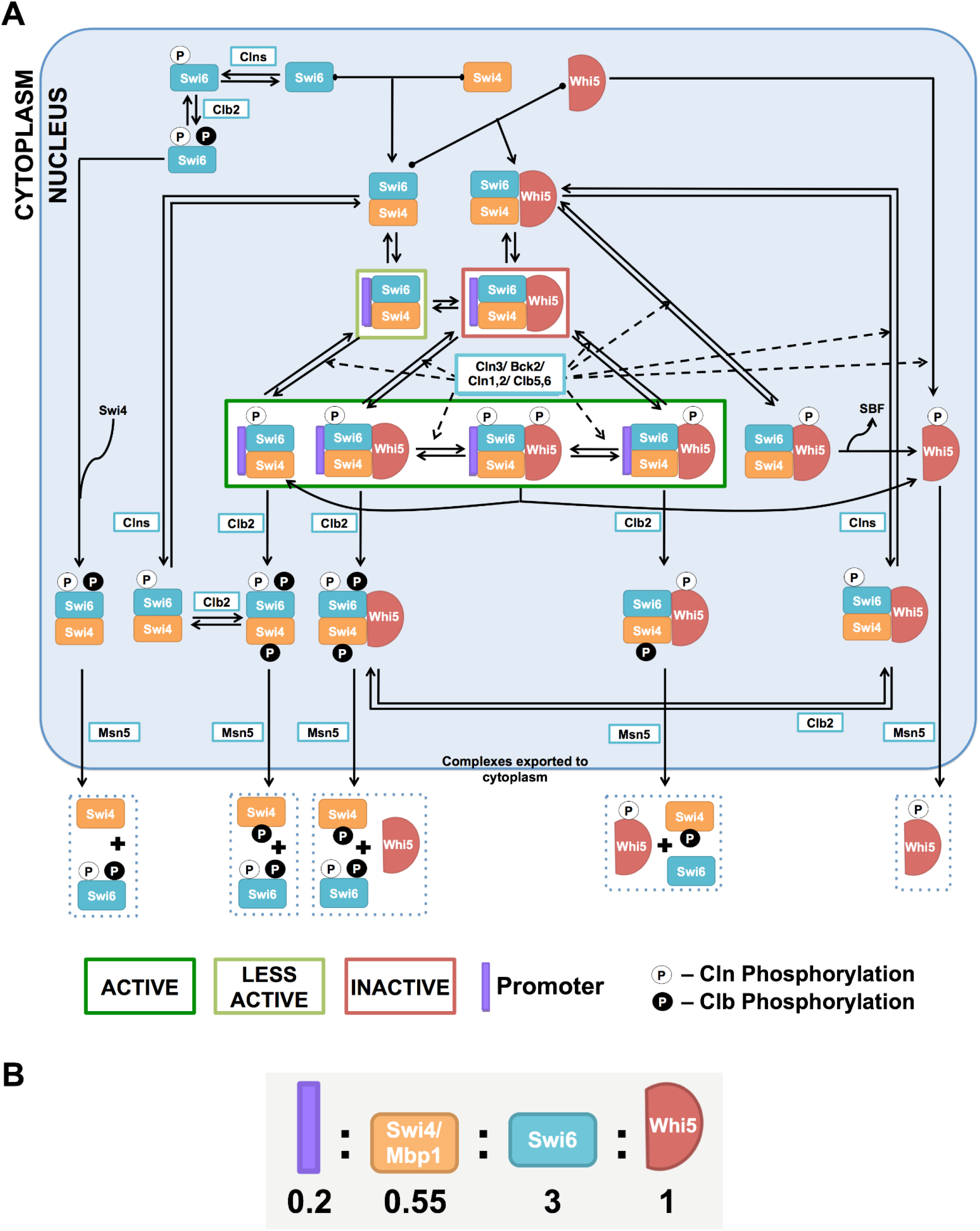
SBF regulation. (A) The core model of SBF activation and inactivation by Clns and Clbs. Presented in the figure are the most important interactions considered in START-BYCC for SBF activation and inactivation. The core components are Swi4 (orange icon), Swi6 (turquoise icon), Whi5 (red icon), the promoter (purple bar), kinases (Cln3, Cln1,2, Clb5,6, Clb1,2) and export protein (Msn5) (white box). The nucleus is represented with a blue-gray background, while the white space corresponds to the cytoplasm. Promoter-bound complexes enclosed in boxes with borders in dark green, light green, and red represent complexes with maximal, residual, and no activity, respectively. White-filled circles represent activatory phosphorylations done by Cln3, Cln1,2, and Clb5,6; whereas the black-filled circles represent the inactivating phosphorylations by Clbs (Clb5,6 & Clb1,2 for Swi6 phosphorylation and Clb1,2 for Swi4 phosphorylation). The Cln (white) phosphorylation on Whi5 and Clb (black) phosphorylation on Swi6 (S160) are needed for export to the cytoplasm. To avoid overcrowding, remaining complexes corresponding to modifications on other free forms are not included in the figure. All the concerned equations are listed under Supplementary Materials. Key facts about the abundance, regulation, and localization of all the components are described in **Table S1**. (B) Ratios of the promoter to Swi4/Mbp1, Swi6, and Whi5 (Roughly based on Ghaemmaghami et al.^77^; e.g., when rescaled, we would have four promoters, 11 each of Swi4/Mbp1, 20 Whi5, and 60 Swi6 molecules).

**Figure 3.**
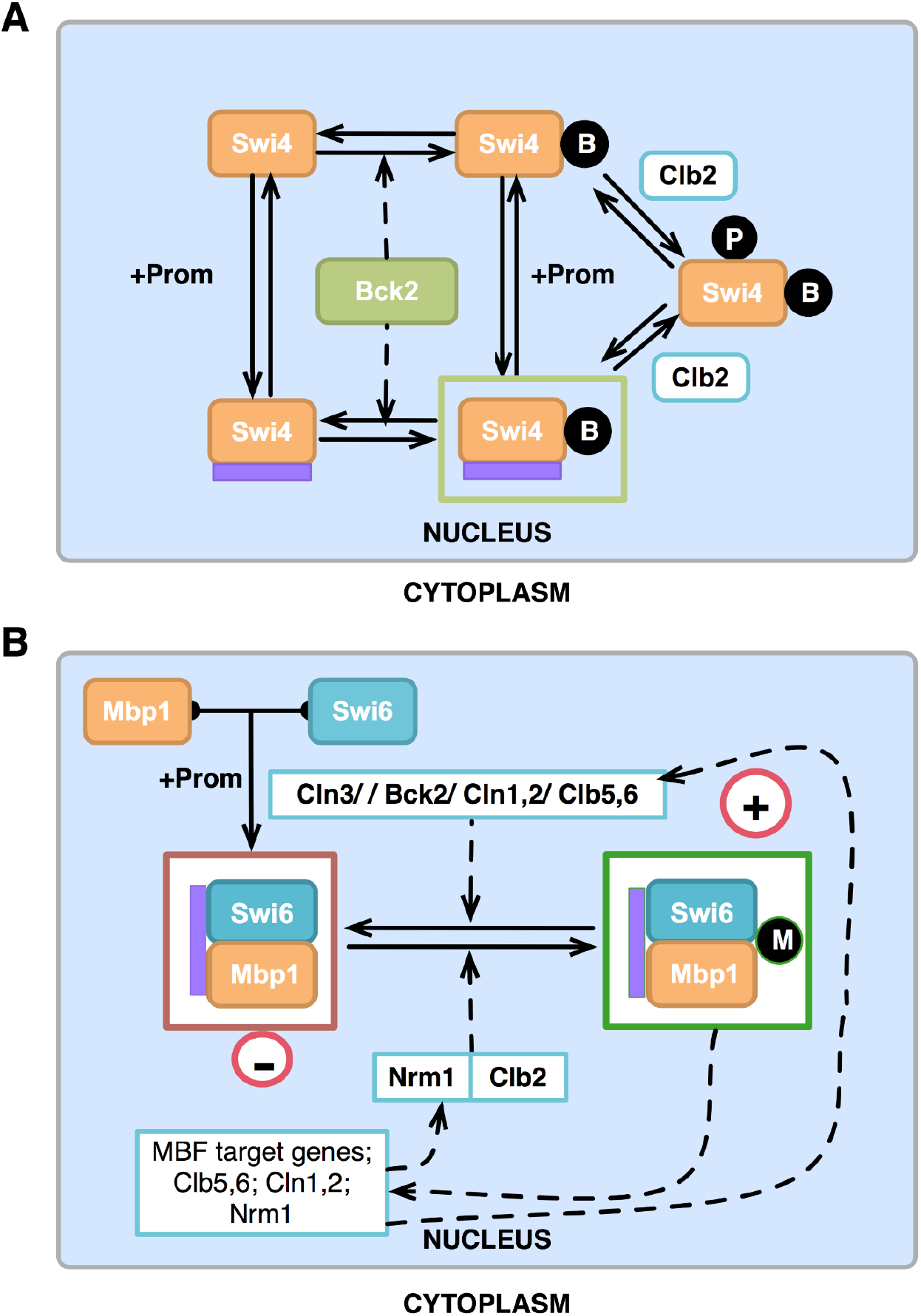
Other mechanisms involved in the START transition. (A) Role of Bck2 in the activation of SBF. Bck2 acts on Swi4 and modifies it to a less active form (indicated by the enclosing light green box). The cartoon is a concise representation of regulation by Bck2 (intermediate steps not included). Black-filled circles with ‘B’ represent the Bck2-induced modification, and active forms are enclosed in a light green box (since these complexes are not as active as Cln-activated forms). Note that only promoter-bound forms are active. Clb2 causes complex inactivation (see equations in Supplementary Materials for details). (B) MBF regulation. MBF alone is inactive in a repressed state (indicated by the enclosing dark red box). It is activated either by cyclins (Cln3, Cln1,2, Clb5,6) or by Bck2 (more active, as indicated by the dark green box). MBF is primarily inactivated by its transcriptional target, Nrm1, resulting in the negative feedback highlighted by the ‘–’ sign. Clb2 is a secondary and minor inhibitor of MBF.

Our current model of the cell cycle with a detailed START transition (START-BYCC; github.com/JRaviLab/start-bycc) is very complex, highlighting the key molecular mechanisms. The START transition subsystem of the model (SBF/MBF regulation) includes 51 species and 56 parameters (**Table S3**), compared to 1 species and 8 parameters in BYCC, allowing us to simulate the START dynamics and yeast genetic system (with knockouts and overexpression) in such detail for the first time. To systematically detail the critical changes we have made, we describe and discuss each new aspect of our current model of START transition in the subsequent sections:

a. a detailed mechanism for the activation and inactivation of SBF (involving Whi5), MBF, and the role of Bck2 in wildtype cells,
b. the timing and localization of different monomers corresponding to activation and inactivation of SBF (Swi4, Swi6, Whi5),
c. the role of phosphorylation in START transition, and
d. the mechanism for cellular size control involving Ydj1 and Ssa1.

We present the cell cycle dynamics and phenotypes of all relevant cell cycle mutants (200+, including >100 START mutants and several critical FINISH mutants from BYCC) as time-course simulations (See *Methods* and *Supp Tables* for details). We successfully fit 95% of the mutant phenotypes (**Table S4**) and describe in some detail key underlying mechanistic aspects of the model and challenges (See *sections below*). We also make a few predictions based on our current detailed model for the budding yeast START transition (**Table 1**). Mathematical models rely on underlying assumptions for their usefulness and accuracy (**Table S2**). To this end, we have highlighted and justified the key molecular players, mechanisms, and their interactions, along with the detailed wiring, simulated here: http://sbmlsimulator.org/simulator/by-start.

**Table 1.**
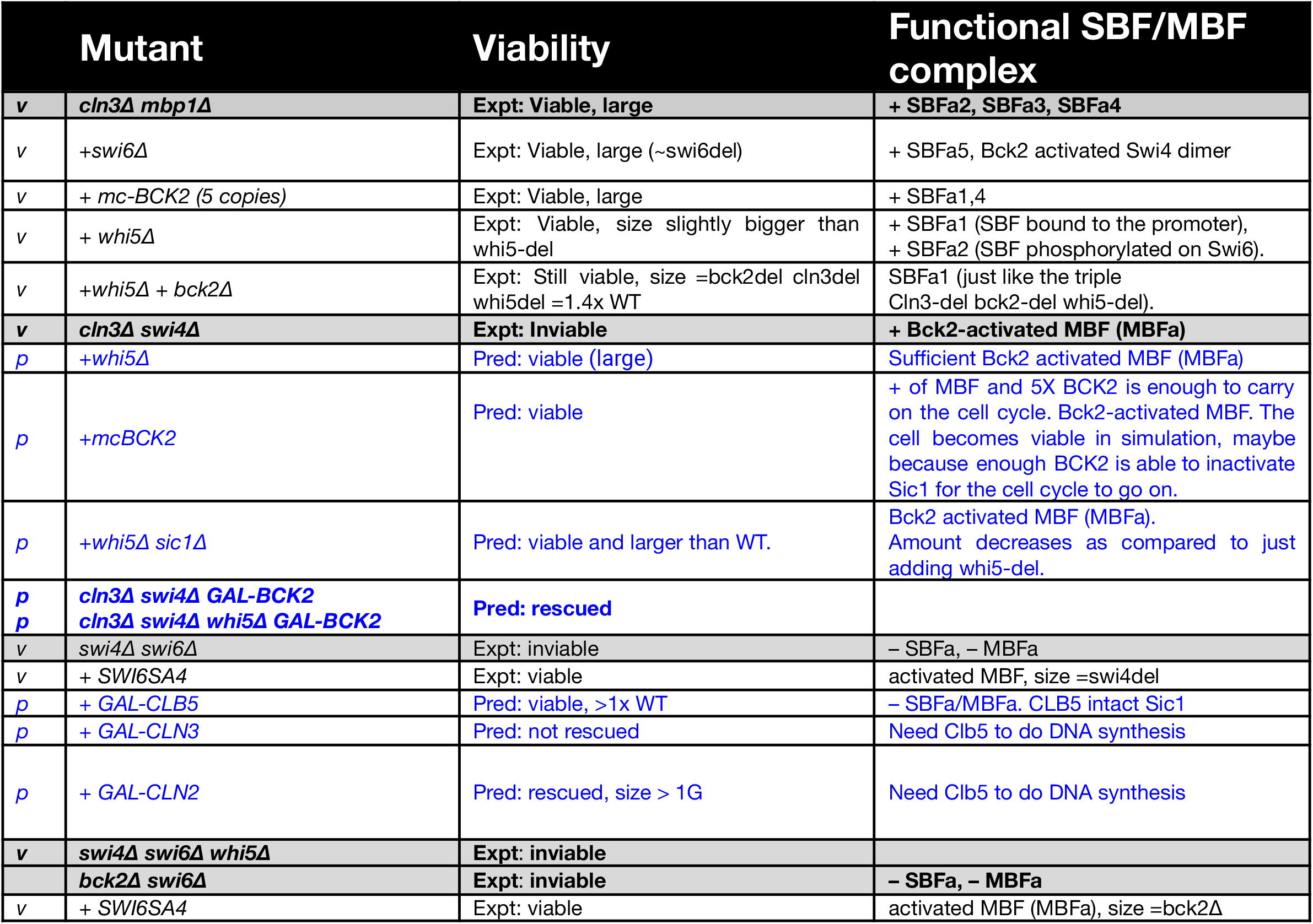
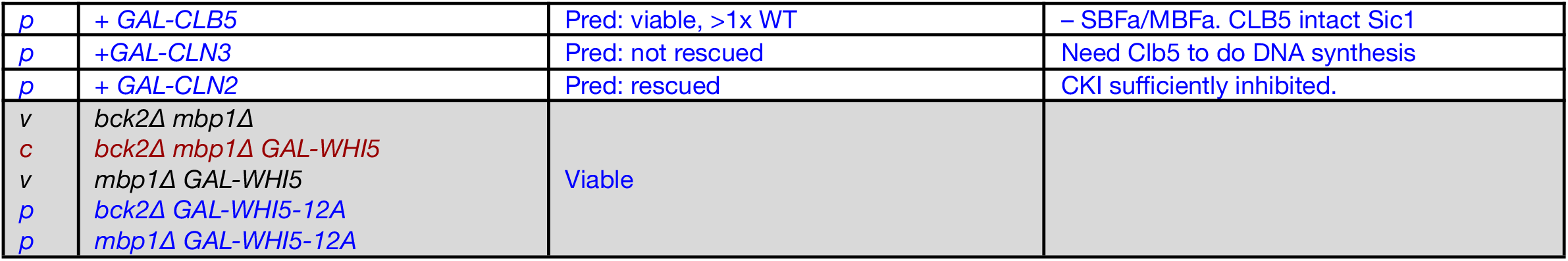
Model Predictions and Validations. (A more exhaustive list of experimentally determined phenotypes is shown in **Table S4**). Single letter notations in column one denote: v (black): validated, p (blue): predicted, and c (red): contradictory. Grey rows indicate parent groups of mutants, with variations that follow.

### Regulation of START in WT budding yeast cells

#### Model description

In order to understand the role of SBF and MBF in the START transition, we first describe molecular events surrounding the monomers and complexes of these transcription factors, the inhibitor, Whi5, and their target promoters (**Fig. S1**). Based on known relative protein levels^77^ we infer that the monomers Swi6, Whi5, Swi4/Mbp1, and promoter exist in the ratio 3:1:0.55:0.2 (**Fig. 2B; Table S2**). These monomers bind together rapidly to form complexes based on their stoichiometry and starting concentrations. Due to their high relative levels, Swi6 and Whi5 are available as free molecules in the cell. Most of Swi4 is bound to Swi6 and Whi5 (*i.e.* most SBF is in the Swi4/Swi6/Whi5 form), and most of Mbp1 is bound to Swi6 forming MBF complexes.

Since the promoters are present in limiting levels, most are occupied by their respective transcription factors, Swi4/Swi6/Whi5 (inactive SBF) or Mbp1/Swi6 (inactive MBF). The remaining SBF and MBF complexes are free and unbound to promoters. This sets the stage to discuss the various regulatory events that ensue during the START transition.

#### SBF activation by Cln kinases in the late G1 phase

Since a transcription factor complex can be transcriptionally ‘active’ only when bound to a promoter, for the following discussion, we focus on promoter-bound SBF complexes and describe their activation by Cln kinases and inactivation by Clb kinases. We assume that a similar kind of regulation occurs on the promoter-free complexes.

When the yeast cell is in G1, most of the promoter-bound SBF complexes are in an inactive, Whi5-bound state. When Cln3 accumulates in late G1, Cln3/CDK activates SBF by phosphorylating Whi5 and Swi6 at several residues^47,48^, resulting in the doubly phosphorylated form. We assume that this form is unstable and dissociates into phosphorylated SBF and phosphorylated Whi5 (**Fig. 2, S2**). Aided by the karyopherin export protein Msn5, phosphorylated Whi5 subsequently moves to the cytoplasm (and stays there until mitotic exit/FINISH transition)^57,59^, leaving SBF transcriptionally active.

#### SBF inactivation in the late S phase

In late S phase, SBF is inactivated by a second round of phosphorylation by mitotic cyclins (Clb1,2; Clb6)^50^ (**Fig. 2, S2**). Presumably, these phosphorylations occur on Swi4^31^ and Swi6 (on residue S160, different from those targeted by Cln3)^58^, leading to dissociation of the complex from the promoter and cytoplasmic localization of Swi6 (facilitated by Msn5^78^).

The START-BYCC model explicitly includes the discrete steps by which important active forms of SBF (enclosed in dark green boxes) are inactivated and exported to the cytoplasm (**Fig. 2, S3**). This series of molecular events comprises a detailed consideration of intermediate active complexes that we do not expect to see in significant portions in wildtype cells. However (as described in later sections), these considerations become useful in understanding the regulatory logic underlying non-phosphorylable mutant phenotypes. Thus, following Clb phosphorylation on the Swi4 moiety of SBF, each of the intermediate SBF complexes dissociates from the promoter, turning off SBF-regulated genes. Then, the free SBF forms (promoter-unbound) move to the cytoplasm, aided by Msn5. We assume that these complexes dissociate once in the cytoplasm and remain there until their corresponding phosphatases reverse their modifications.

#### Activation of SBF by Bck2

In addition to activation by cyclins, SBF is also activated by Bck2 in late G1^49^. The viability of *cln3Δ* and the inviability of the double mutant *cln3Δ bck2Δ*^49^ emphasize the importance of Bck2 in SBF activation. Bck2’s mechanism of action is not very clear, but it is thought to be independent of CDK phosphorylation and Whi5^47,49^. In START-BYCC, we consider a mechanism by which Bck2 modifies SBF to alternate active forms (**Fig. 3A**). We determine the relative contributions of Cln3 and Bck2 to SBF/MBF activation from the relative sizes of cln3Δ and bck2Δ mutants (*cln3Δ* >> *bck2Δ* >> WT; see the section on the role of Bck2^49,79^). We assume that the complexes activated by Bck2 are less active than the Cln-activated forms and indicate this with light green outlined boxes (**Fig. 3A**).

In addition to the Swi4/Swi6 heterodimer of SBF, we assume that (in the absence of Swi6) Swi4 too has some residual activity and that Bck2 activates homodimeric Swi4. This assumption becomes important in the context of mutants *swi6Δ*, *swi4Δ swi6Δ*, and *swi6Δ* in the genetic background of *cln3Δ* or *bck2Δ* (described below).

#### Regulation of MBF

Alongside SBF, the other important transcription factor operating at START is MBF, a heterodimer of Mbp1 and Swi6. The primary cell cycle targets of MBF considered in START-BYCC are Clb5,6. Due to the functional overlap between SBF and MBF^26^, we assume, in START-BYCC, that MBF can also activate Cln1,2 (targets of SBF) and that SBF can activate Clb5,6 (targets of MBF).

MBF is activated by both Cln3 and Bck2 by independent mechanisms and inactivated by Clb2 and by Nrm1, one of MBF’s targets^80^ + Cln2 and Clb5 activate MBF, albeit to a lesser extent. In START-BYCC, we have incorporated the positive and negative feedback loops for the activation and inactivation of MBF, as depicted in **Fig. 3B**.

### Wildtype simulations of the budding yeast cell cycle

The various mechanisms described thus far (**Fig. 2, 3**) correspond to the typical set of biomolecular interactions that occur in a wildtype cell through the cell cycle. In START-BYCC, we have incorporated several intermediate reactions that could potentially occur only in specific overexpression or knockout mutants. For example, **Fig. S3** considers intermediates that we expect to observe only in non-phosphorylable mutants. The model also considers different pools of complexes (free and promoter-bound) and monomers, and mechanisms for their modification and localization (**Table S2**). The levels (and significance) of each of these complexes/intermediates depend on the amounts of starting monomers, the nature of the mutant, and the growth rate (reflected in the mass doubling time).

We convert this entire picture^20^ (highlighted in **Fig. 2, 3, S3**) into ordinary differential equations, each reflecting the dynamic fate of one biomolecular entity (variable). These equations are then used to run time-course simulations of the system (solved numerically with the equations, parameters, and initial conditions; details provided in **Supplementary Text;** http://sbmlsimulator.org/simulator/by-start). Results from a typical time-course simulation of a wildtype cell are shown in **Fig. 4**. Each graph tracks the concentration of different sets of variables in normalized units spanning 300 min (∼3 cell cycles). By default, cells grow in glucose with a mass doubling time of 90 min (in reasonable agreement with Brewster 1994^81^). The total cycle time for the daughter corresponds to 107 min and the G1 length is 62 min.

**Figure 4.**
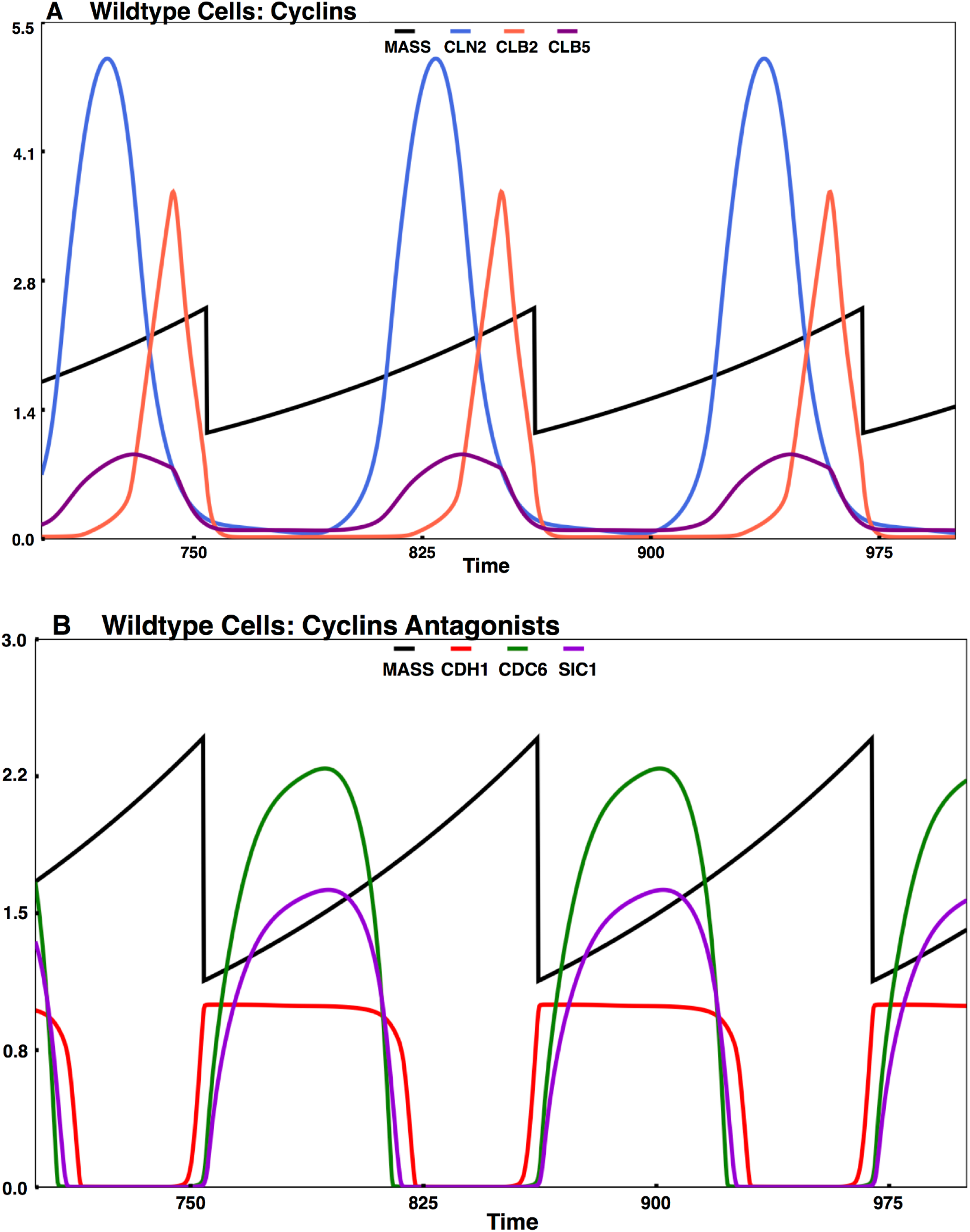
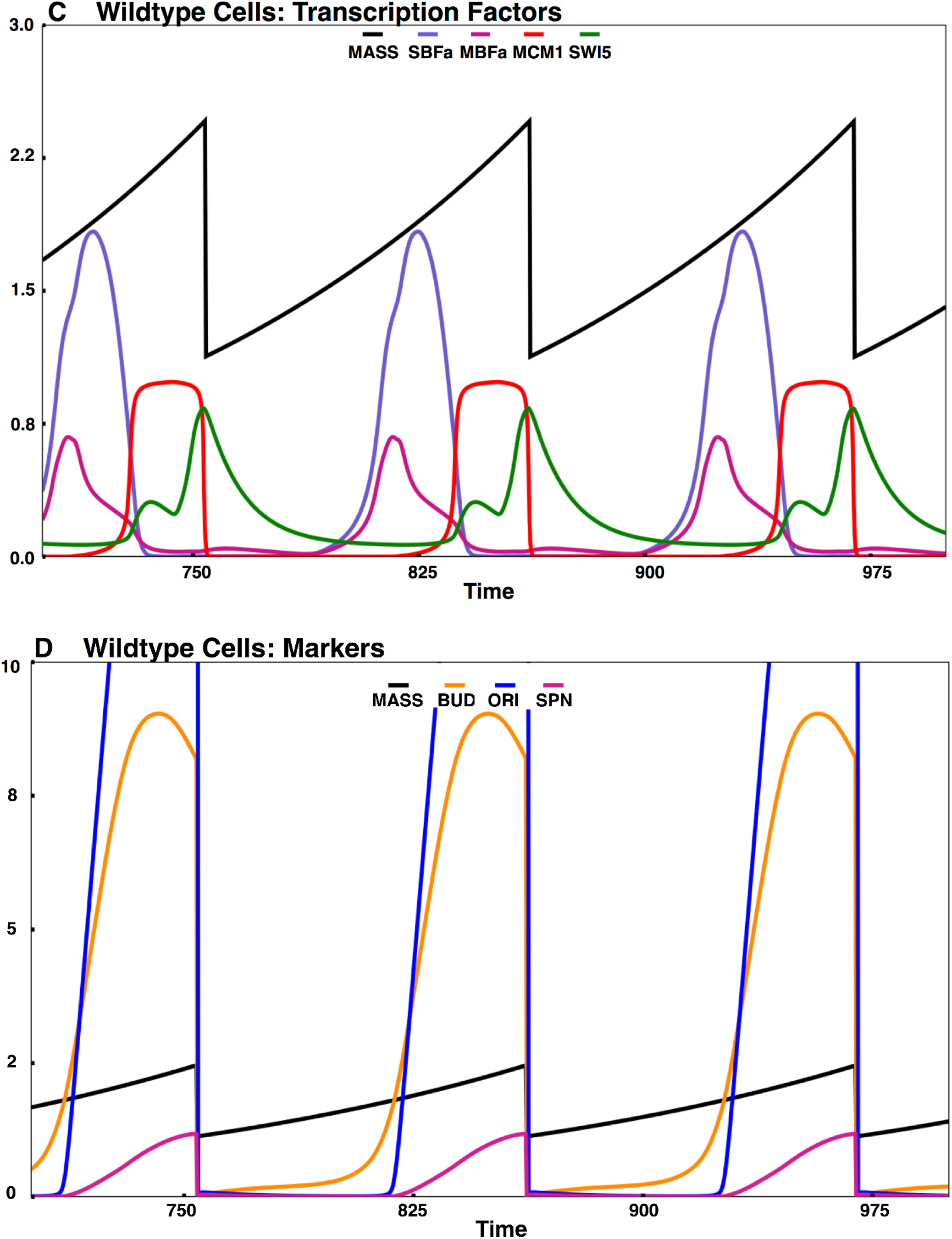
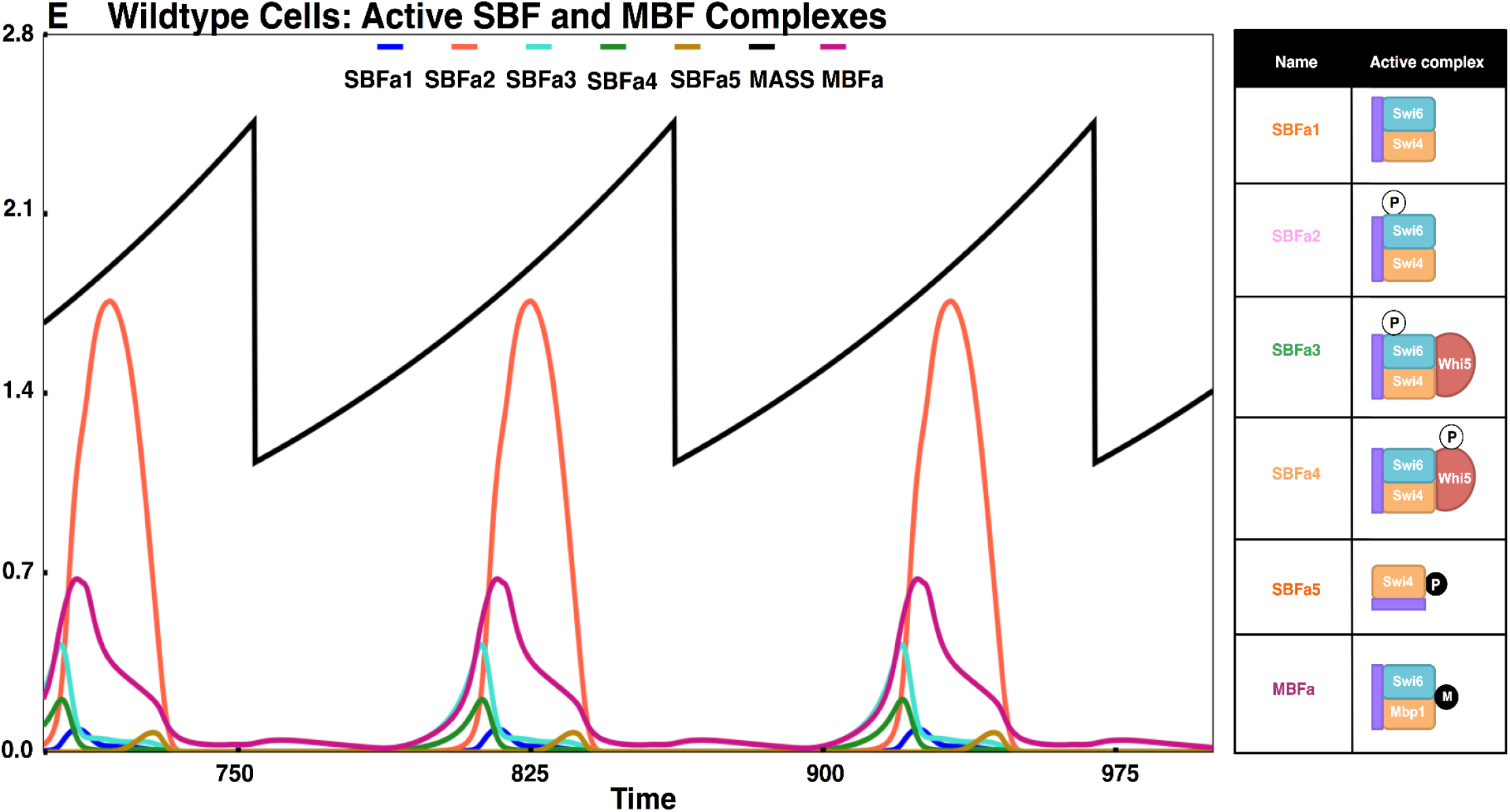
Simulation of wildtype cells in glucose. Each panel tracks the following proteins/components for ∼3 cell cycles in glucose with mass doubling time 90 min and daughter cycle time ∼107 min. (A) Cyclins (Cln2, Clb5 (active), Clb2 (active)), (B) Cyclin antagonists (Cdh1, Sic1 (active), Cdc6 (active)), (C) Transcription factors (SBF, MBF, MCM1, Swi5), (D) Markers (CDK targets (BUD, ORI, SPN) used to indicate the occurrence of physiological events via concentration threshold (BUD=1 (bud emergence), ORI=1 (DNA synthesis initiation), SPN=1 (spindle alignment in metaphase)), (E) Active SBF and MBF complexes. Each form shown (SBFa1-a5, MBFacln, MBFabck) is the activity of SBF or MBF contributed by a particular form. The complexes contributing to the different forms (all bound to promoter) are as follows: SBFa1=unmodified SBF (SBFB in the model); SBFa2=SBF phosphorylated on Swi6 (SBFB6P+SBFB6PQ); SBFa3=SBF-Whi5 complex phosphorylated on Swi6 (WSB6P+WSB6PQ); SBFa4=SBF-Whi5 complex phosphorylated on Whi5 (WSB5P); SBFa5=Swi4dimers activated by Bck2 (Swi4B); MBFa = MBF activated by Clns or Bck2. Check and cross marks denote the presence or absence (due to mutation) of specific complexes. The absence of any sign denotes that the specified complex is absent in the simulation of that strain. The black curve in all panels denotes mass (corresponding to exponential cell growth and division). As shown in (E), for wildtype cells, the dominant form of SBF is SBFa2, the form of SBF phosphorylated on Swi6, whereas the dominant form of MBF is the Cln-activated MBF.

Our current expanded model captures the cellular dynamics of cyclins, cyclin antagonists, transcription factor complexes, and checkpoint proteins (**Fig. 4, A–D**), similar to the BYCC model^20^. Wildtype cells start in G1, with G1 stabilizers active (high Sic1, Cdc6, Cdh1). In late G1, as cells reach a size threshold to trigger the START transition, a nuclear build-up of G1/S activators Cln3 and Bck2 results in the activation of SBF, MBF, and subsequently, Cln1,2 and Clb5,6. The activation of the feedback between the cyclins and SBF/MBF ensures an irreversible START transition^82,83^ (when not under starvation). The nuclear export of phosphorylated Whi5 is another hallmark of SBF activation (in accordance with Di Talia 2007^57^). In turn, the S phase cyclins (Cln1,2; Clb5,6) lead to concurrent initiation of DNA synthesis (origin of replication, ORI) and bud formation (BUD), as observed experimentally. We set the firing of the origin, ORI=1, as the marker for START. Cln3 remains nuclear until late G1 and moves to the cytoplasm in late G2 (as observed by Verges et al., 2007^76^).

Transcriptional activation of S phase cyclins by SBF and MBF causes CKI degradation (by Cln1,2) and Cdh1 inactivation (by Clb5,6). This results in the accumulation of mitotic cyclins, Clb1,2 (further aided by the Clb1,2, Mcm1 feedback), and entry into the M phase. Following this transition, sister chromatids align at the metaphase plate, with ensuing Cdc20 and Cdc14 activation resulting in exit from mitosis. Since the current model focuses on START, we use the simplified, older version of BYCC to emulate the mitotic exit and FINISH transition. Incorporation of our detailed START model with our more recent detailed FINISH models^41^, and the relevant detailed descriptions of FEAR and MEN pathways and the spindle position checkpoint, are beyond the scope of this work^84,85^.

In line with the expansion of the molecular mechanism underlying START, we track not only the total complexes, but the distribution between various active forms of SBF and MBF (**Fig. 4E**). In later sections, we present simulations of mutant cell types similar to the wildtype plot of SBF/MBF complexes in **Fig. 4E**. For viable cells, we show cycles after the cell has reached a steady state. The cell size at division in our simulations roughly corresponds to the mean cell volume determined in experiments. In addition to quantitatively measuring cell size from simulations, we can also deduce a qualitative size for different mutant cell types by comparing the amounts of their active SBF and MBF complexes (icon table beside **Fig. 4E** and other mutant simulations). In all cases (wildtype and other mutant phenotypes), we report the observed phenotypic size of the mutants as fold-change relative to wildtype cells in glucose.

### Size Control

Nutrient modulation of size control represents a key aspect of the budding yeast cell cycle^7^, wherein budding yeast exhibits a thresholding effect – i.e., cells must grow to a critical size to initiate budding and S phase. This control is achieved through tight coupling between cell growth and division. We, therefore, revise the molecular details of the size control mechanism in START-BYCC for a more accurate depiction of size control and its coupling to cell cycle.

#### Cln3 activation of SBF, MBF

Cln3 is implicated in size control, since mutants with Cln3 deletion or increased Cln3 cytoplasmic export are larger than WT^86,87^ and mutants overexpressing Cln3 or with increased Cln3 nuclear import are smaller than WT. Cln3 was thus thought to be a nuclear sensor of cell size, triggering START only when the cell reached the threshold size. Although Cln3 has been identified as an important regulator of START, Cln3 knockout mutants are still viable. This is due to the activity of Bck2, which is activated by glucose and known to promote START^49^.

The earlier models of the cell cycle, including BYCC^19,20^, incorporate the role of Cln3 in sensing size in a direct mass-dependent manner: as cell volume grows, the total Cln3 protein level in the cell increases in parallel, with its concentration remaining constant. As its level rises, Cln3 migrates to the nucleus and concentrates there. Assuming the nuclear volume does not change significantly, the nuclear concentration of Cln3 grows proportionally to cell size. When the cell reaches a threshold size, this mass-dependent nuclear accumulation of Cln3 (& Bck2) triggers START by activating SBF abruptly (modeled with a classic ultrasensitive Goldbeter-Koshland GK switch)^64^. Thus, the START transition behaves like a switch in the BYCC model^19,20^. Recent experiments, however, suggest that the nucleus grows proportionally to the cell during the cell cycle, challenging this hypothesis^88^.

#### Size control in START-BYCC

We treat cell size control as the outcome of actively regulated nuclear import of Cln3 (and Bck2) prior to START. Recent experiments show that, in early G1, the Cln3/Cdc28 complex is sequestered to the endoplasmic reticulum (ER) membrane by Ssa1/2, Whi3, and other negative regulators of START. In late G1, the J-chaperone protein Ydj1 accumulates in response to cell size, relieving this inhibition on Cln3 and enabling the nuclear accumulation of the Cln3/Cdc28 complex^76^. Thus, Ydj1 is believed to act as the cell size sensor via positive regulation upstream of Cln3.

The START-BYCC model assumes that Ydj1, in response to increasing mass, moves Cln3 from ER into the nucleus abruptly in late G1. Nuclear Cln3 phosphorylates and inactivates Whi5, resulting in nuclear export of Whi5 and activation of SBF^47,48^. Since Ssa1 is the protein that retains (and thus inhibits) Cln3 in the ER, we assume that Ssa1 inactivates Cln3^76^. In order to explain the observation that Cln3 is nuclear from late G1 to late S/G2, we propose that Ssa1 is activated by Clb2 and Swi5 (**Fig. S4**). Since little is known about the regulation of Bck2, we assume that its regulation is similar to that of Cln3. We have modified the size control mechanism from the BYCC model to reflect these hypotheses and assumptions, but retain Cln3 and Bck2 as the ultimate size sensors (albeit through Ydj1/Ssa1^10,11,57^).

Results from simulations of START-BYCC compare well with observations from a classic experiment in size control measuring cell size distribution for different nutrient media and growth rates^89^ (**Fig. 5**). Slower growth rates result in smaller cell volumes (taken to be mass at division in START-BYCC simulations) and higher growth rates (richer nutrient media) result in higher median cell volumes. Furthermore, we show that START-BYCC reproduces the observation that daughter cells display a progressive G1 delay (activation of ORI/BUD in the model) in response to slower growth rates (poorer nutrient media) that is characteristic of strong size control^89^ (**Fig. S5A**). Also, in line with our expectations, our simulation shows that the longer cycle times in slower growth media occur mostly due to a delay in G1 (wherein smaller, slower-growing cells wait to reach the size threshold), leaving the budded phase almost constant (**Fig. S5B**).

**Figure 5.**
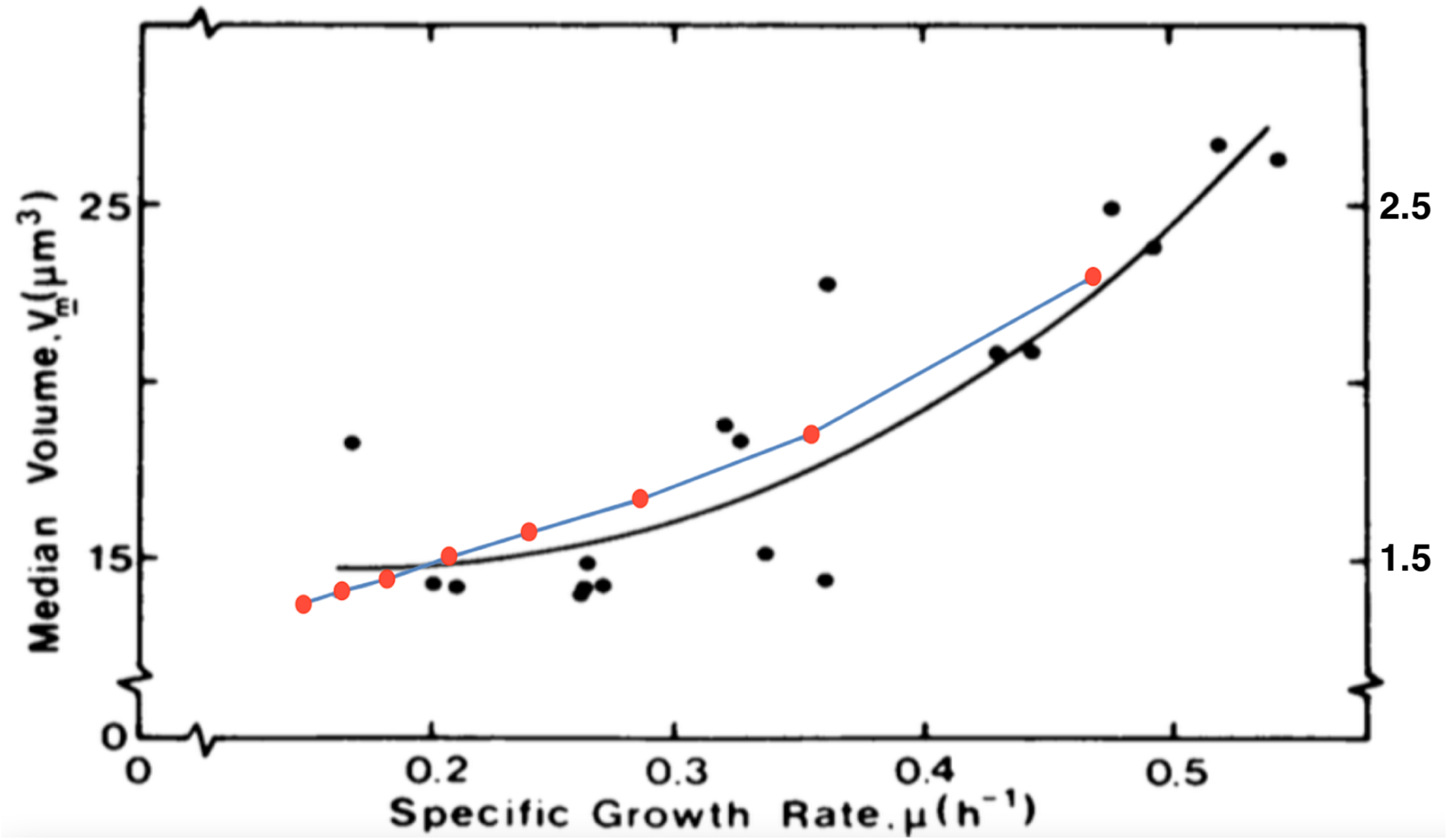
Cell size as a function of growth rate in budding yeast (daughter cells). Comparison with the experiment by Lord and Wheals (1980)^89^ (Fig. 6 used with permission, see License in *Supplementary Material*). The blue curve connecting red circles shows cell size at division at specific mass doubling times (MDT) from 90-300min (30 min interval) and corresponding specific growth rates (ln2/MDT) from START-BYCC simulations. This curve is overlaid upon the experimental graph where median cell volumes for populations at different growth rates are plotted. The experimental curve is from the original paper^89^.

### Export and Localization of START proteins

Several studies on START transition have emphasized the crucial role of localization of the core proteins (in their various modified/complexed forms)^57–59,75^. We consider the export and localization of the START monomers by building a compartmentalized model (nucleus, cytoplasm inside a single cell). The ratio of the sizes of the nucleus and cytoplasm is predefined (0.2:0.8 in our case) and remains constant throughout the cell cycle^88^. We detail the specifics of the localization of Whi5, Swi6, and Swi4 below, and summarize them in **Fig. S6**.

#### Whi5 localization

Single-cell experiments reveal Whi5 is in the nucleus only in G1, moving to the cytoplasm with the onset of the START transition and remaining there until the next cycle begins^57^. We have encoded this behavior in the model by allowing phosphorylation of all forms of Whi5 in the nucleus by G1 and G1/S cyclins, Cln3, Cln1,2, and Clb5,6. These phosphorylation events signal both free and SBF-bound Whi5P to be exported to the cytoplasm in late G1 by the transport protein Msn5 (**Fig. S6**). Additionally, any Whi5 bound to SBF with Swi6 phosphorylated by Clb kinase (Q-form; see below) is also exported to the cytoplasm. Cytoplasmic Whi5P is then dephosphorylated by Cdc14, which is activated at mitotic exit (**Fig. S6, Step 4**).

#### Swi6 localization

Swi6 localization is known to depend on the serine residue S160, which, when phosphorylated in late S-phase by Clb6, allows export to the cytoplasm^58^. In the Swi6 S160A mutant (serine mutated to alanine), Swi6 remains in the nucleus throughout the cell cycle. START-BYCC designates this S160 phosphorylated form as the ‘Q-form’ (black phosphorylation site in **Fig. S6**) to distinguish it from the other activatory phosphorylations on Swi6 by Cln kinases (designated ‘P-forms’; white phosphorylation site in **Fig. S6**). In addition to Clb5,6^50^, we assume that Clb1,2 can also bring about the S160 phosphorylation (Q-form). Swi6 in its Q-form is exported to the cytoplasm (in the presence of Swi4, aided by Msn5) and stays there from the late S phase through the end of the cycle^78^. In START-BYCC, the phosphatase, PP2A dephosphorylates the P-form (**Fig. S6, step 3**). Similar to Whi5, we assume that the final dephosphorylation (on Q-form) of Swi6 happens at the mitotic exit by Cdc14, followed by quick re-import^50,90^ (**Fig. S6, step 4**).

#### Swi4 localization

Unlike Whi5 and Swi6, Swi4 shows nuclear localization throughout the cell cycle^75^. For this reason, we assume that phosphorylated Swi4 (irrespective of the compartment) is dephosphorylated by an unspecified active phosphatase (Ppase in our case) without allowing it to wait for Cdc14 to accumulate at mitotic exit (**Fig. S6, step 3**).

#### Simulation of localization and export of monomers

Through numerical simulations based on the afore-mentioned formalization, we have captured nearly all localization events of monomers during different phases of the cell cycle (**Fig. 6**). Firstly, we examine the profiles of phosphorylated and cytoplasmic monomeric components and the timing of their nuclear export. We observe that phosphorylated Whi5 is exported to the cytoplasm in late G1, while phosphorylated Swi6 is exported after START (late S phase). Both monomers stay in the cytoplasm until mitotic exit (**Fig. 6**). There is a small level (<10%) of phosphorylated Swi4 in the cytoplasm in S/G2 phases of the cell cycle. These simulation results correspond well with previously discussed experimental findings on the localization of Whi5, Swi6, and Swi4. We observe that most of Swi6 goes to the cytoplasm in late S phase, following Whi5. This is because Swi6 constituting MBF, and certain phosphorylated forms of SBF are incapable of moving to the cytoplasm in START-BYCC. Only the doubly phosphorylated Swi6 forms are capable of cytoplasmic export (**Fig. 2, S6**).

**Figure 6.**
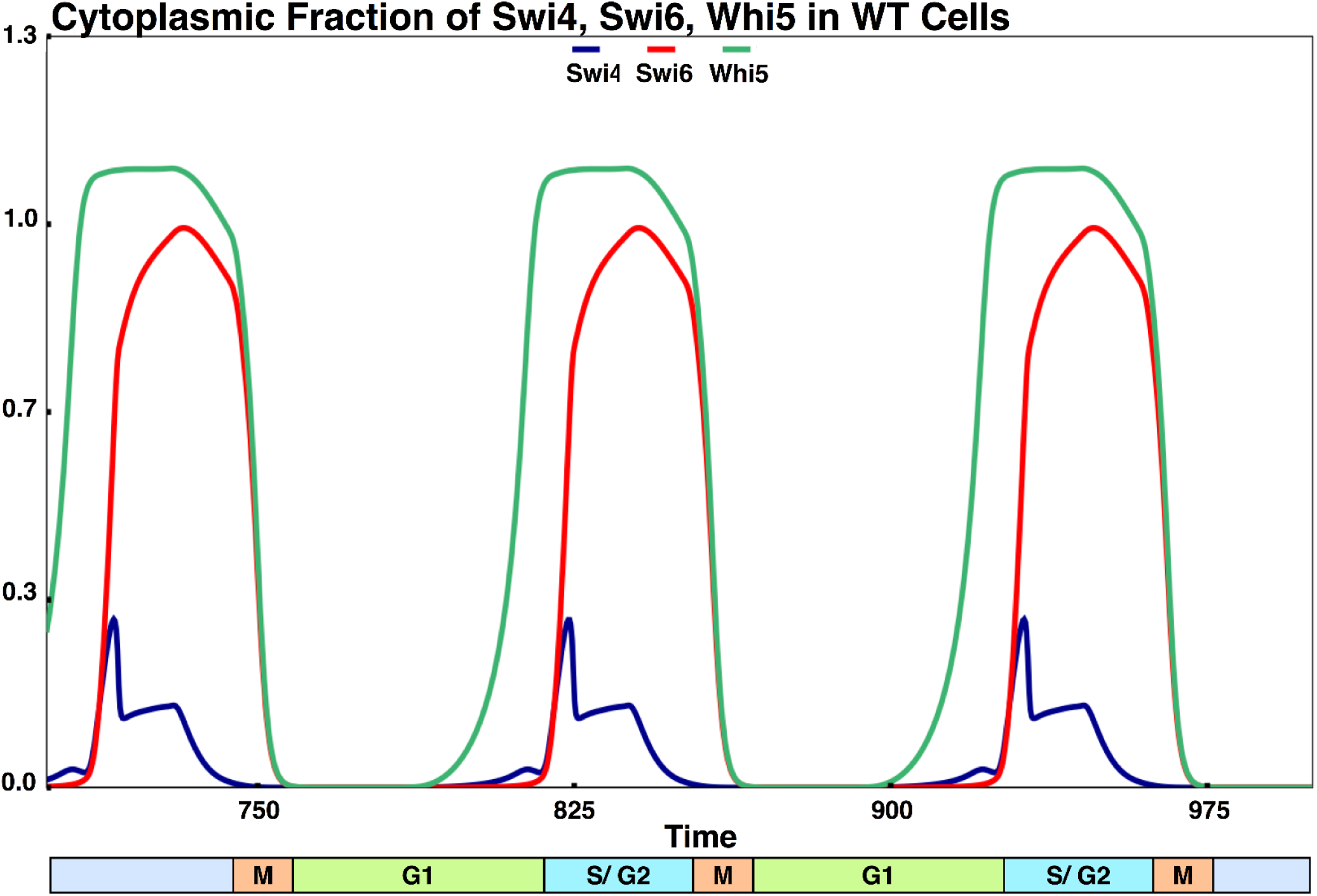
Simulation of the timing of localization (export) of different monomers. The cytoplasmic fraction of monomers Swi4, Swi6 and Whi5 are plotted against time. The bar at the bottom of the graph shows timing w.r.t. phases of the cell cycle. In the model, the onset of S and M phases correspond to DNA synthesis (ORI=1) and spindle assembly checkpoint (SPN=1), respectively. In compliance with experiments, Whi5 enters the cytoplasm (exits the nucleus) in late G1, followed by Swi6 in S-phase. Both Swi6 and Whi5 are cytoplasmic until mitotic exit. Swi4 is mostly nuclear at all times. Details are discussed in the main text.

Secondly, we are also able to reproduce the results pertaining to the importance of the export protein Msn5 in the cell. *msn5Δ* mutants are known to be larger than WT cells^78^, and our simulations show that these cells are indeed considerably larger, since SBF localization is upset and there is only active MBF (**Fig. S7**). In *msn5Δ*, we observe that both Whi5 and Swi6 are nuclear at all times, in accordance with experiments.

In summary, for wildtype budding yeast cells, our current START-BYCC model can explain the dynamics and underlying mechanisms by recapitulating known experimental phenotypes, from START and the rest of the cell cycle. Next, we discuss extending our model to explain mutant phenotypes.

### Case of the non-phosphorylable mutants

We use the START-BYCC model to describe several mutant phenotypes, most of which are implicated in the START transition. Since a critical gap in our previous understanding of START is the complex role of phosphorylation in regulating the transition, we start with the detailed characterization of the non-phosphorylable START mutants: single mutants *WHI5-12A*, *SWI6-SA4*, and the double mutant *WHI5-12A SWI6-SA4*. Recent studies have shown that there is a delay in the START transition only when neither Whi5 nor Swi6 can be phosphorylated^58,59^. This is contrary to the previous belief that Whi5 phosphorylation is essential for relieving the inhibition of SBF^20,47,48^.

The START-BYCC model accommodates these new findings by considering all possible phosphorylation states and intermediate complexes of Whi5, SBF, and the target promoter (**Fig. S8**). In the mutant *WHI5-12A*, Whi5 cannot be phosphorylated, but the phosphorylation sites on Swi6 are intact (**Fig. S8**). Therefore, only Swi6 gets phosphorylated in both SBF and SBF-Whi5 complexes to yield the active transcription factor (**Fig. S8A**). On the other hand, in the *SWI6-SA4* mutant, Swi6 cannot be phosphorylated, but the phosphorylation sites on Whi5 are intact. Therefore, in this mutant, the transcription factor complex can be phosphorylated only in its Whi5-bound form (**Fig. S8B**). In both single mutants, we assume that the phosphorylated transcription factor complexes are fully active (akin to the complexes present in wildtype cells) so that there is no difference in size (**Fig. 7A–B**).

**Figure 7.**
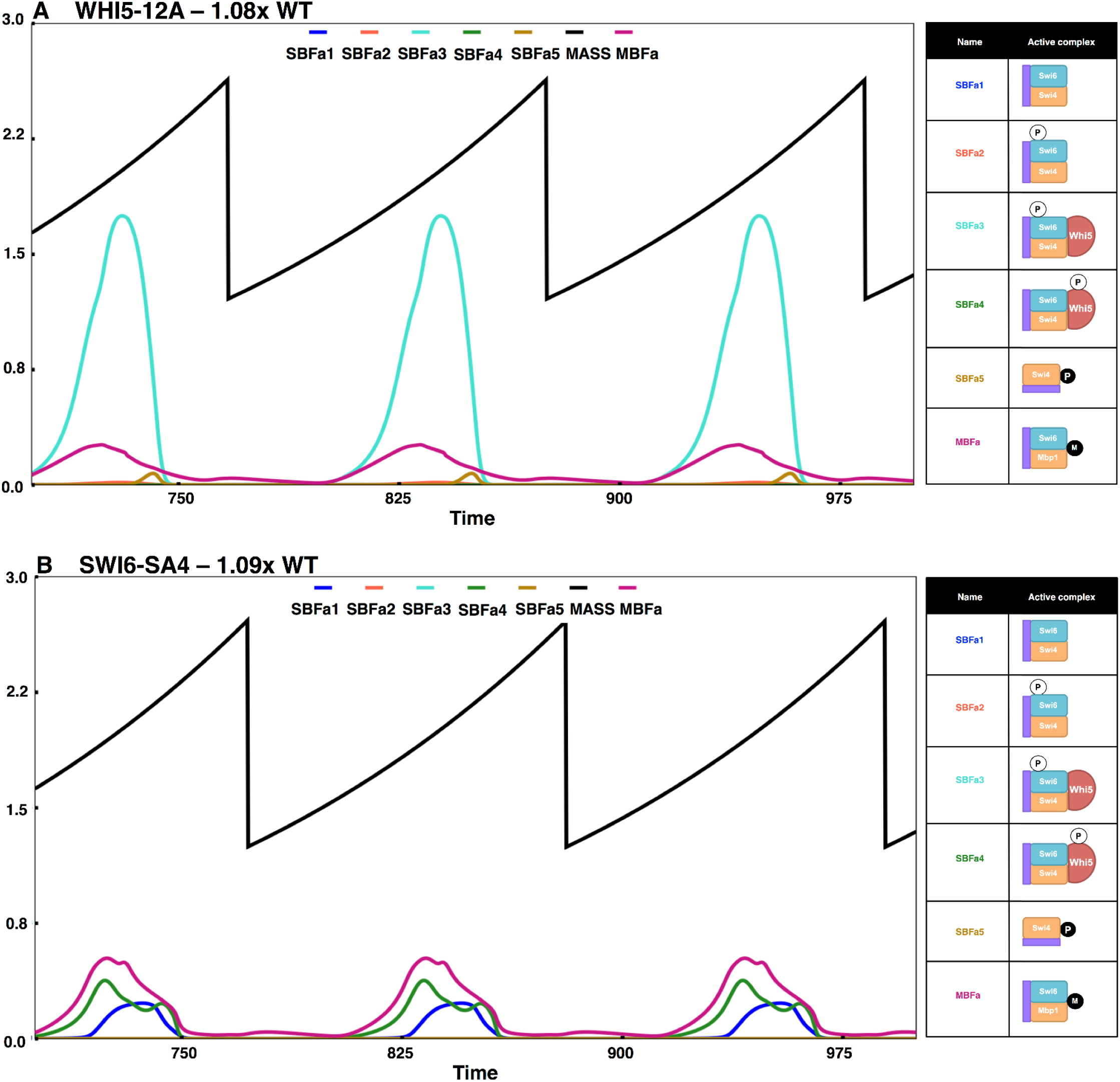
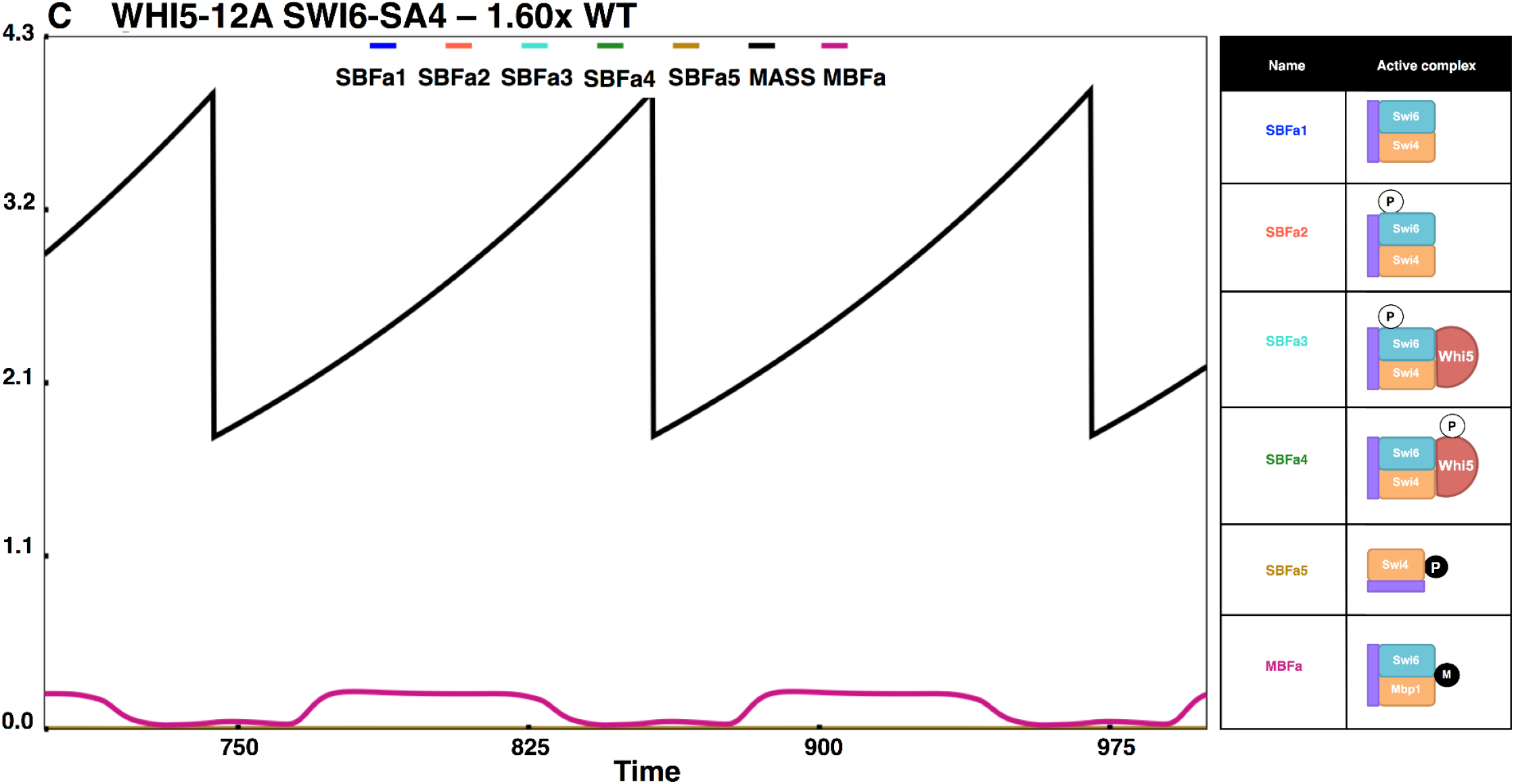
Simulation of non-phosphorylable mutants. The simulations shown here track the cell size and key active SBF/MBF fractions, in specific non-phosphorylable START mutants, and the descriptions are as in Figure 4. (A) *WHI5-12A* (SBF-Whi5 complex phosphorylated on Swi6 and MBF are the primary active forms – SBFa3, MBFa; cells are ∼WT size), (B) *SWI6-SA4* (SBF activated by Bck2, SBF-Whi5 complex phosphorylated on Whi5, and MBF are the primary active forms – SBFa4, MBFa; cells are ∼WT size), (C) *WHI5-12A SWI6-SA4* (only very little of Bck2 activated MBF forms are present –MBFa; cells are larger than WT).

In the double mutant *WHI5-12A SWI6-SA4*, neither Swi6 nor Whi5 can be phosphorylated (**Fig. S8C**). Consequently, the SBF-Whi5 complex remains unphosphorylated and inactive. Even though promoter-bound unmodified SBF has some residual activity and Bck2 is still present, excess Whi5 renders SBF inactive (**Fig. S8C**). So, in the double mutant, SBF is completely off. Despite the absence of active SBF, these cells are still viable, albeit large (since nuclear Whi5-12A can inhibit MBF, and Swi4 can compete with Mbp1 for Swi6). The viability of *WHI5-12A SWI6-SA4* can be explained by the presence of the almost intact MBF that can transcribe Clb5,6 and Cln1,2 (due to functional overlap with SBF; **Fig. 7C**). The double mutant *GAL-WHI5-12A SWI6-SA4* is inviable because the excess Whi5 can inhibit the activity of MBF (**Fig. 8J**), in line with experimental observations^47,48^. In START-BYCC, Whi5-12A is a stronger inhibitor of SBF than wildtype Whi5 due to its nuclear localization. Similarly, Swi6-SA4 makes SBF weaker than wildtype Swi6 (weaker form dominates), as observed in the difference in size between *bck2Δ* and *bck2Δ swi6Δ SWI6-SA4* (slightly larger) (**Fig. 8B, O**).

**Figure 8.**
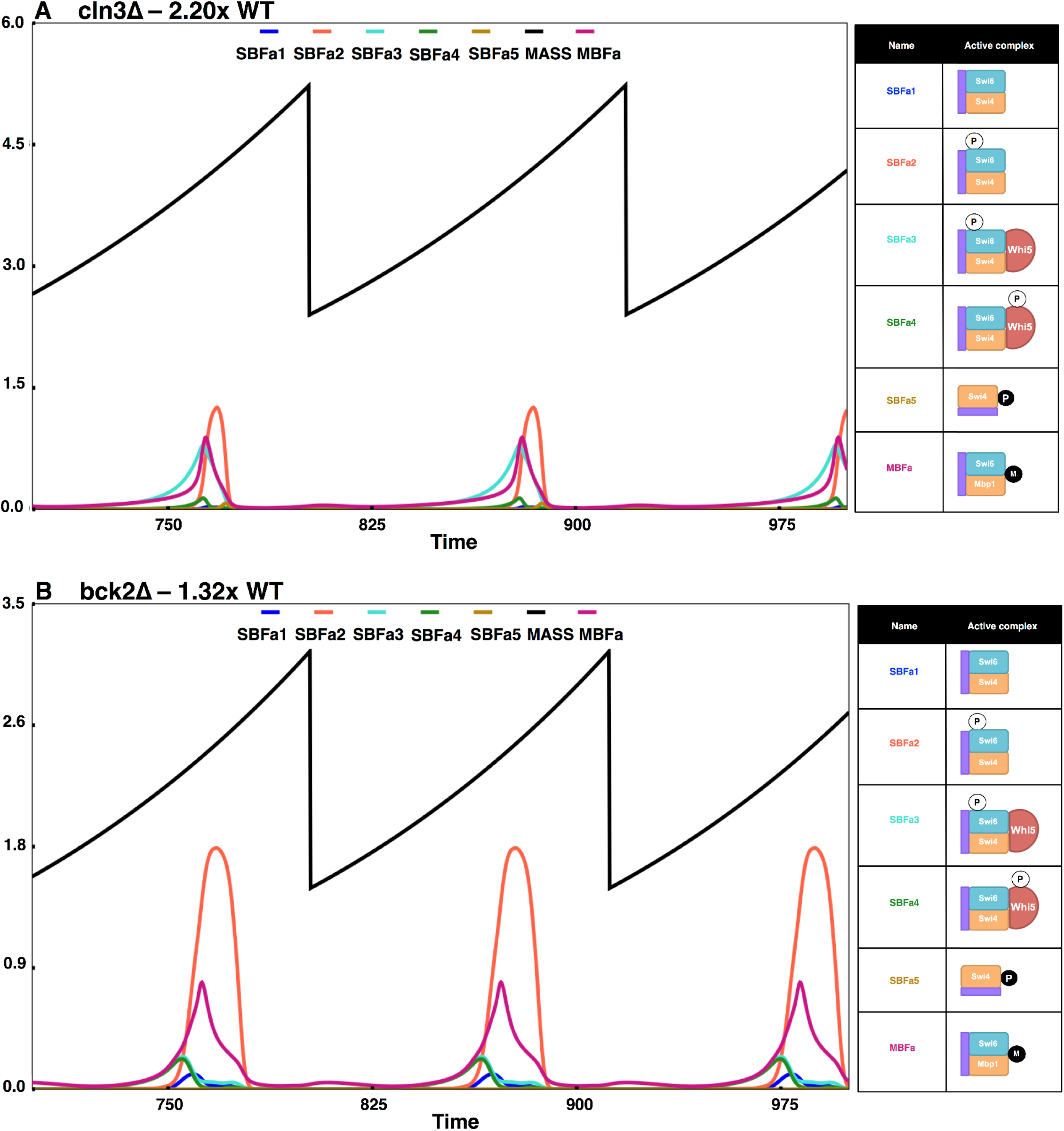
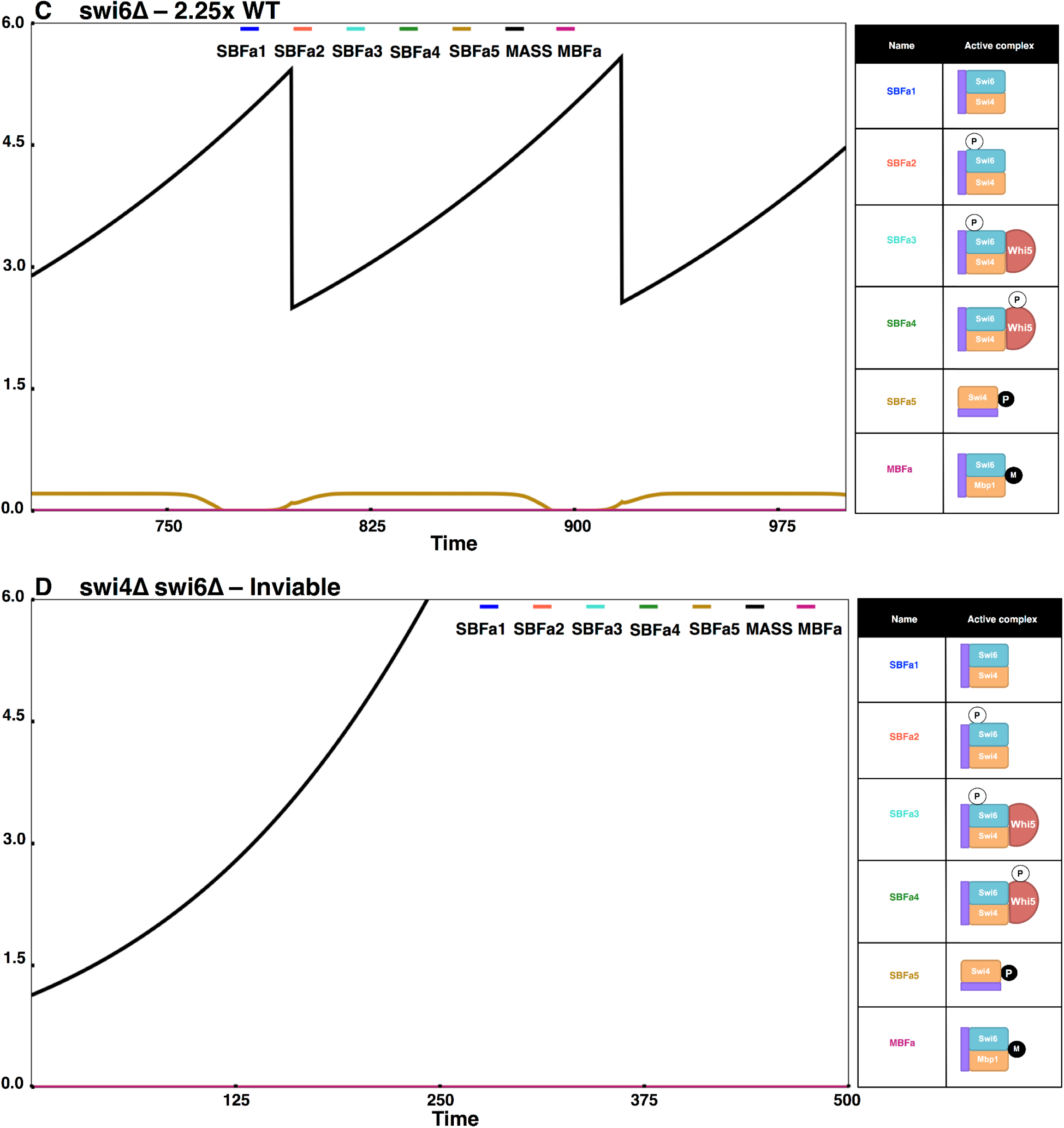
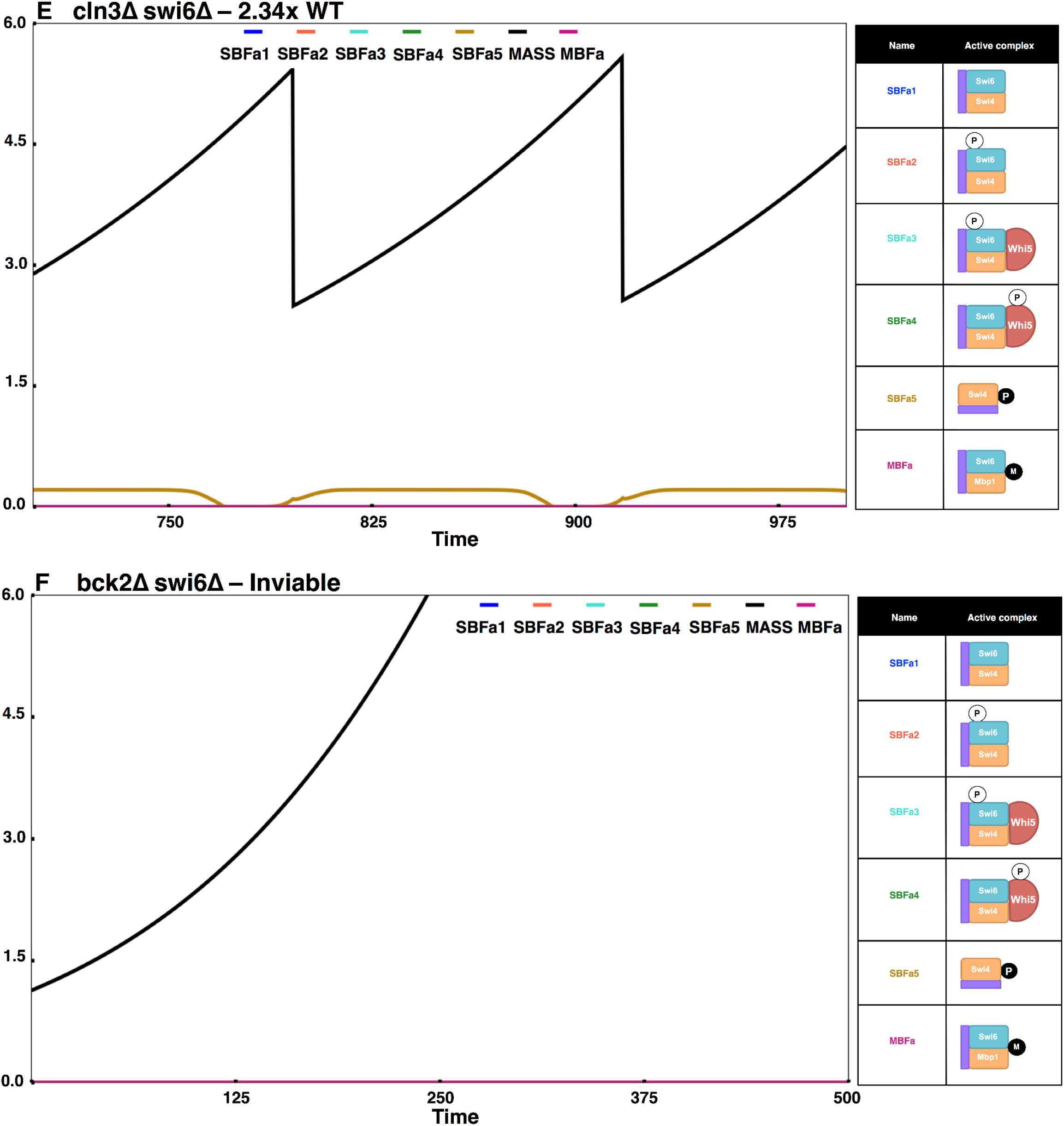
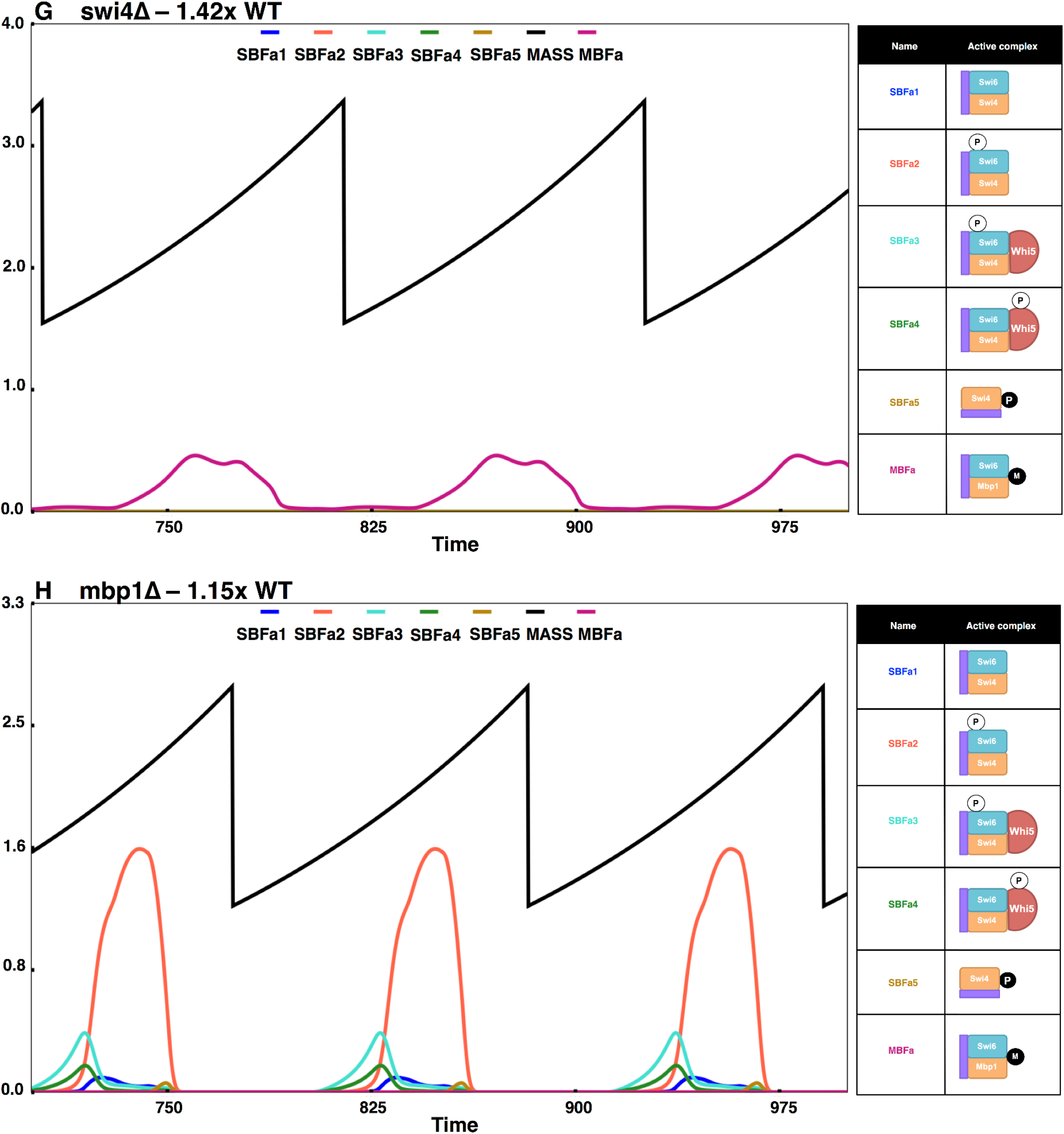
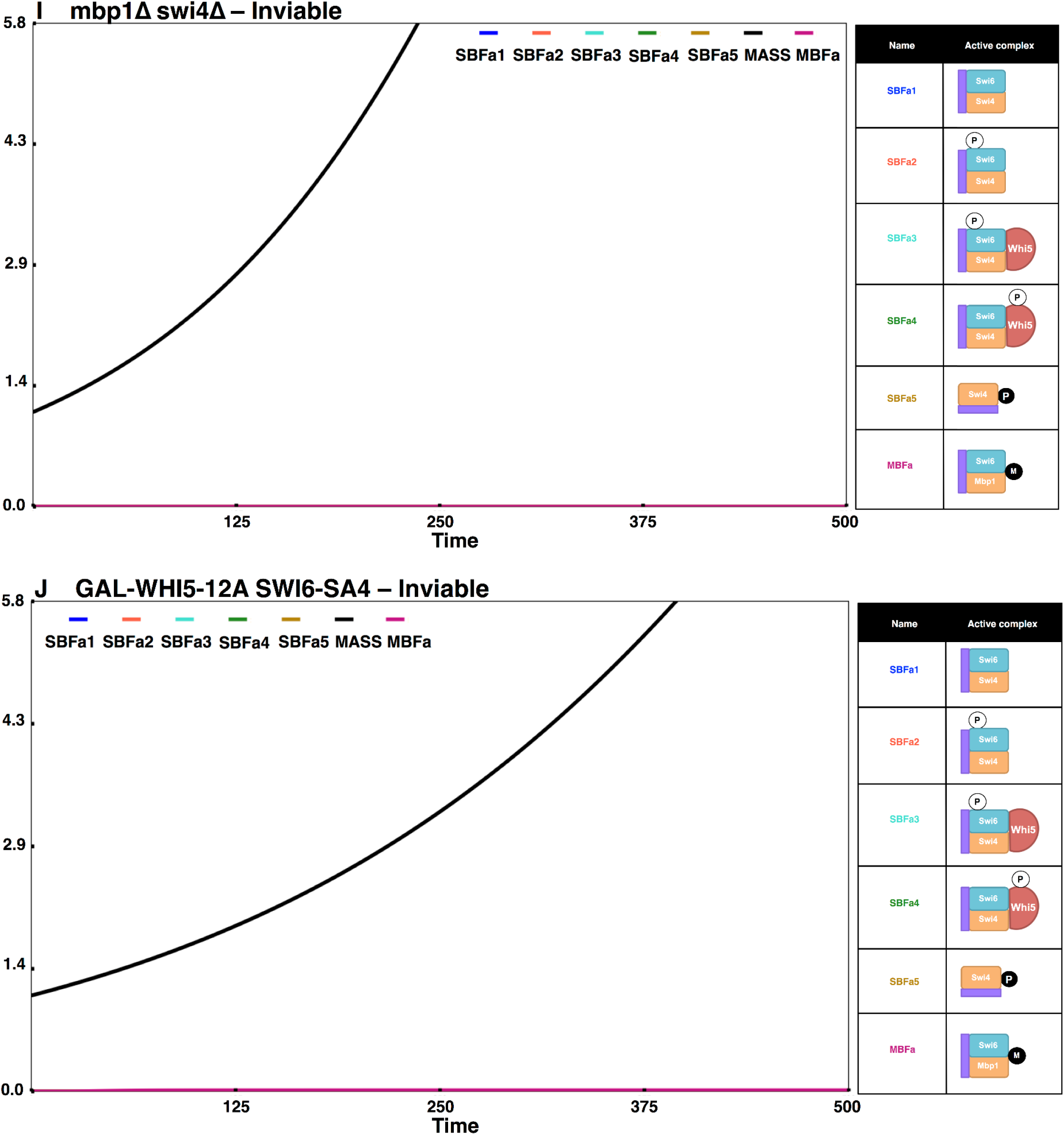
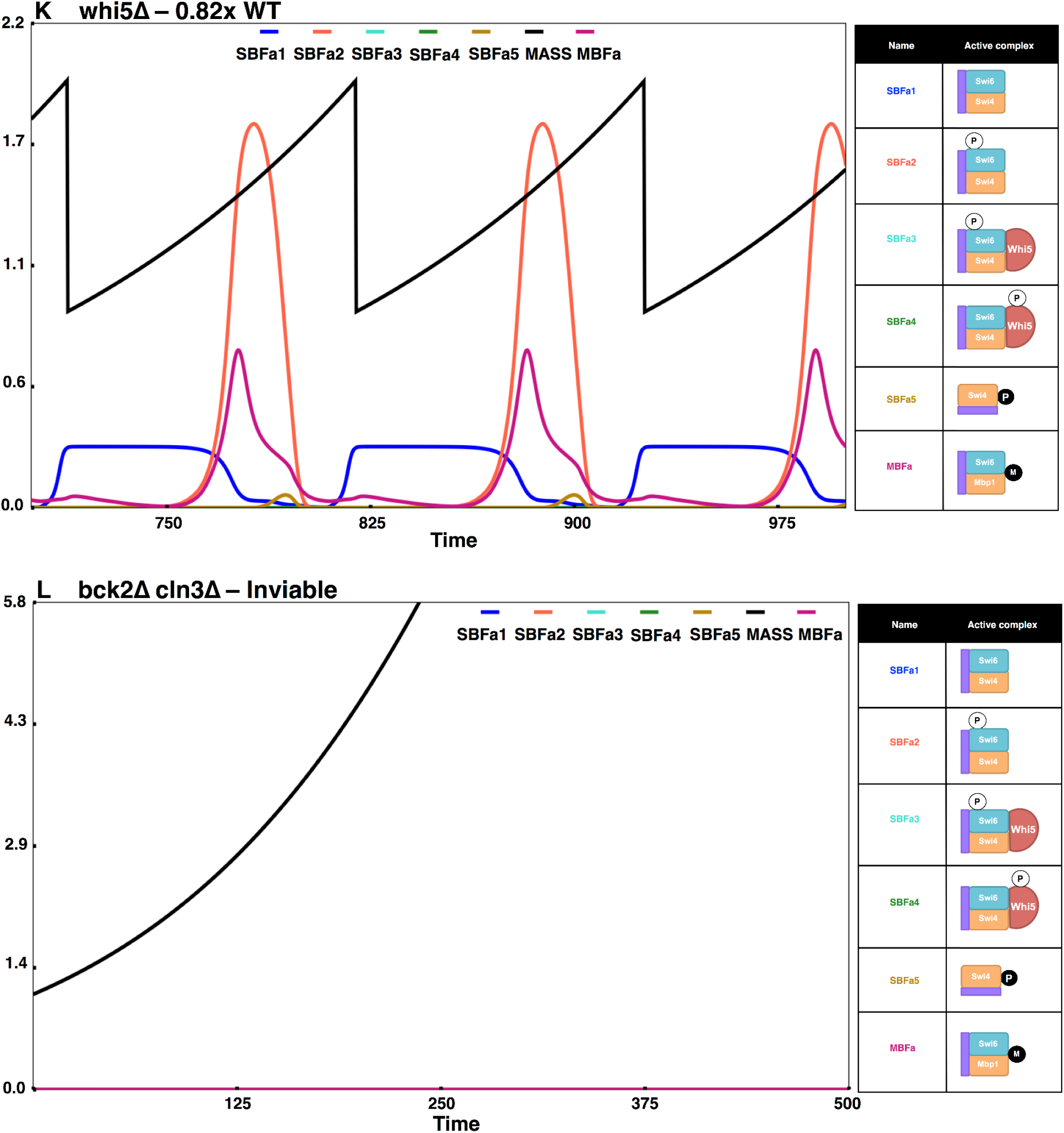
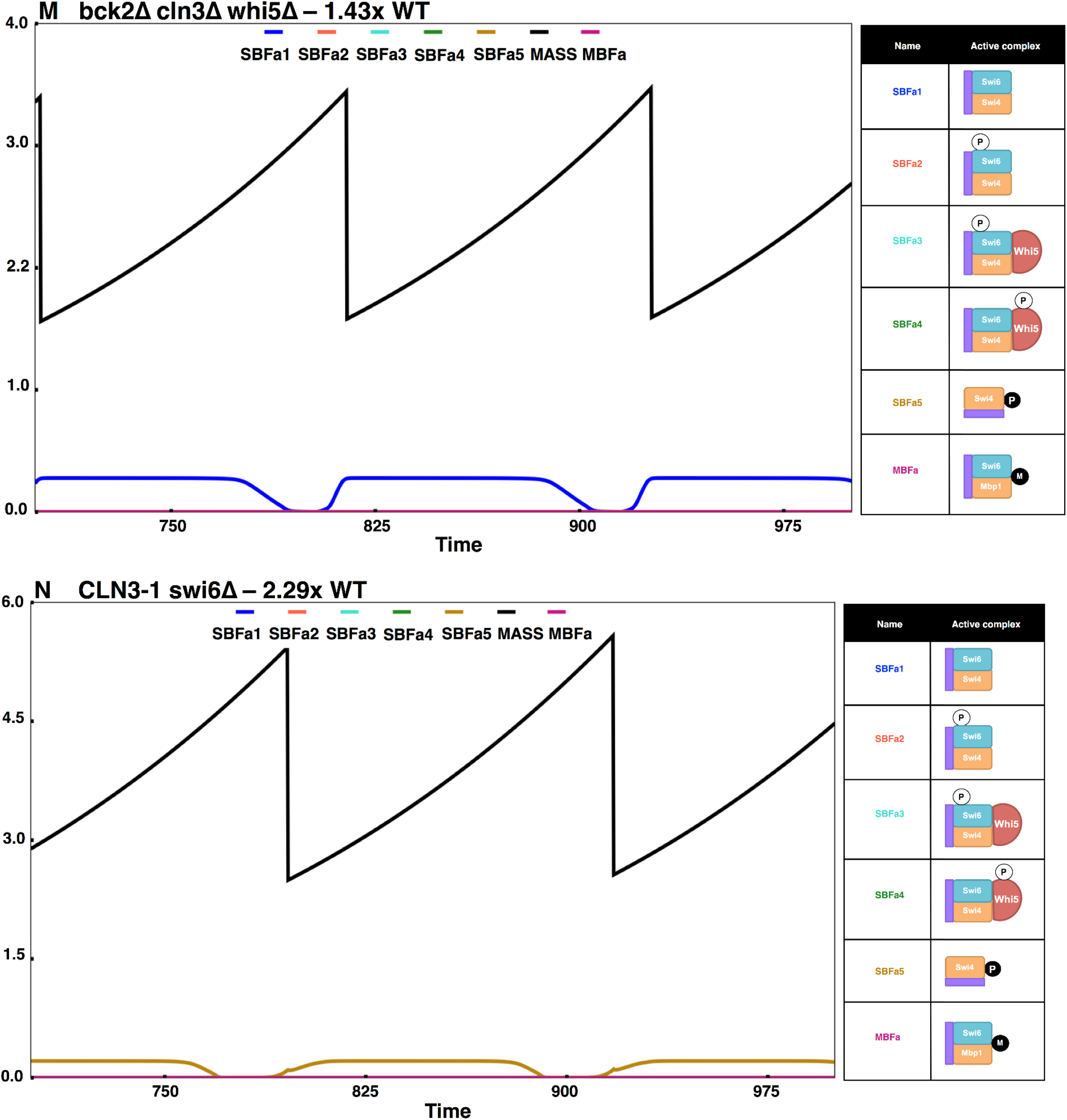
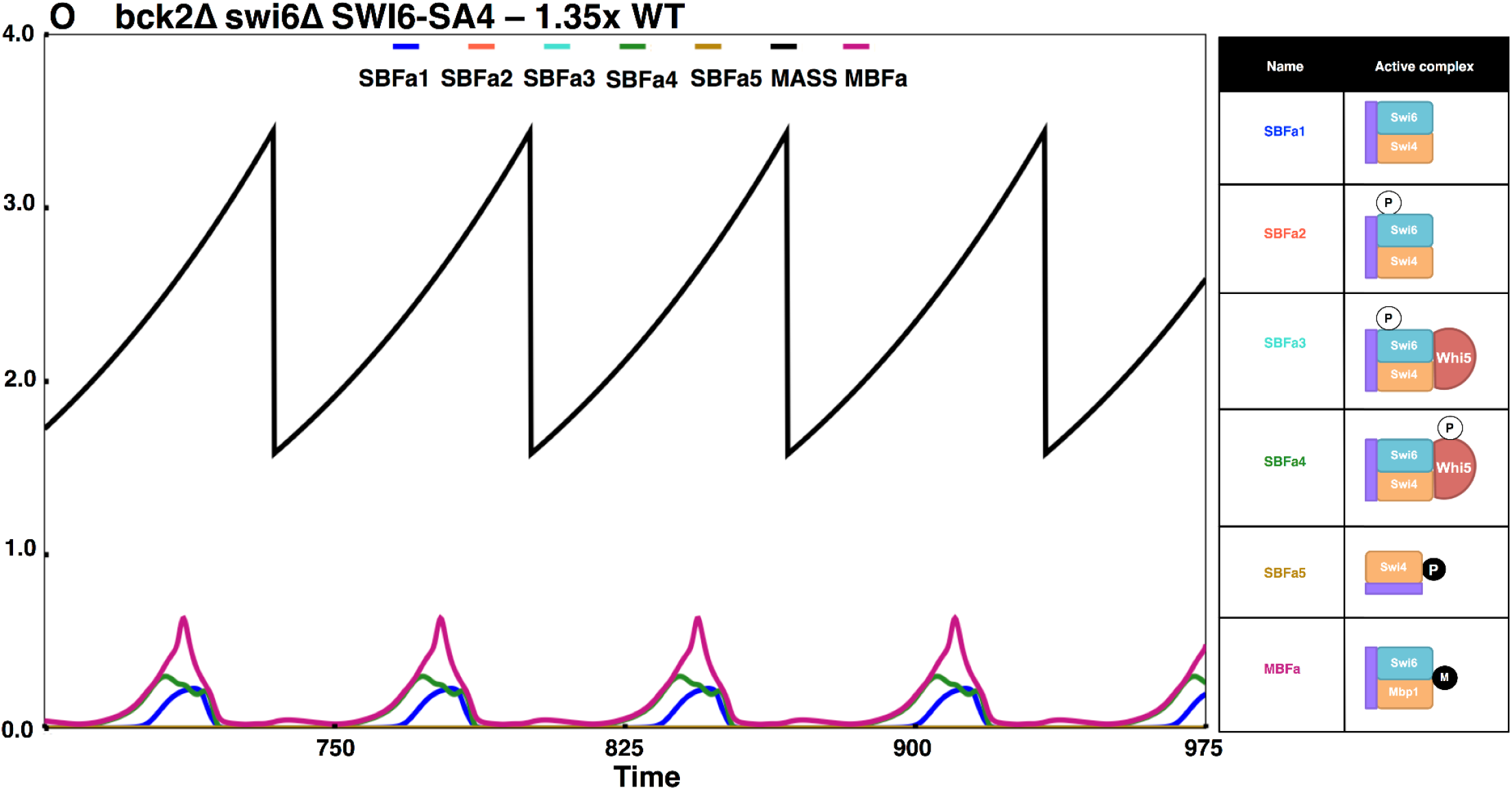
Simulation results of a few important START mutants. Simulations of the following mutants and their steady state sizes (viability/inviability) and the active SBF/MBF complexes present are shown: (A) *cln3Δ* (only Bck2 activated forms are present; cells are very large), (B) *bck2Δ* (only Cln-activated forms are present; cells are slightly larger than WT), (C) *swi6Δ* (only Swi4dimers (SBFa5) present; cells are viable yet large), (D) *swi4Δ swi6Δ* (no SBF/MBF; cells are inviable), and (E) *cln3Δ swi6Δ* (only Swi4dimers (SBFa5) present; cells are viable yet large), (F) *bck2Δ swi6Δ* (no active SBF/MBF; cells are inviable), (G) *swi4Δ* (only MBF present; cells are very large), (H) *mbp1Δ* (only SBF present; cells are slightly larger than WT), (I) *swi4Δ mbp1Δ* (no SBF or MBF; cells are inviable), (J) *GAL-WHI5-12A SWI6-SA4* (excess non-phosphorylable Whi5 inhibits MBF; cells are inviable), (K) *whi5Δ* (cells begin the cycle with less active SBF (SBFa1) instead of Whi5-bound inactive SBF, and get converted into more active forms by Clns (SBFa3). Active MBF is also present. Hence, cells are smaller than WT.), (L) *bck2Δ cln3Δ* (no active SBF/MBF; cells are inviable), (M) *bck2Δ cln3Δ whi5Δ* (deletion of Whi5 relieves SBF and the unmodified form of SBF (SBFa1) is consistently present and cells become viable), (N) *CLN3-1 swi6Δ* (only Swi4dimers (SBFa5) present; cells are viable yet large), (O) bck2Δ swi6Δ-SA4 (SBFa1, SBFa4, and MBFa present; cells are slightly larger than WT).

In summary, START-BYCC and simulations explain in detail the role and working of these non-phosphorylable mutants by replicating the experimental findings with the sizes of *WHI5-12A* (**Fig. 7A**) and *SWI6-SA4* (**Fig. 7B**) being comparable to WT, but the double mutant significantly larger (**Fig. 7C**).

### Simulation results of mutant phenotypes

Using our current model of the cell cycle, we have simulated several mutants pertaining to START and other phases of the cell cycle (as per **Table S4, Fig. 8**). We have summarized our results in **Table S4** where we list the mutants, their phenotypes as observed in experiments, and the results from the simulations of our current model. Our model predictions agree with all the known expected phenotypes, and some have been experimentally validated^91^.

In the following subsections, we present simulations of a few important START mutants that highlight the roles Bck2, Cln3, and the transcription factor monomers Mbp1, Swi6, and Swi4. In each simulation plot corresponding to the mutants, we have included iconic representations (cartoon depictions) of the major active SBF and MBF complexes to follow the relative abundances of these complexes and the resulting mutant phenotypes. We also indicate the names of these complexes below.

#### Mutants pertaining to the role of Bck2

The role of Bck2 in START is not well understood. We, therefore, use START-BYCC to simulate mutants of Bck2 and Cln3. In the *cln3Δ* mutant, most SBF complexes are bound to Whi5 and are inactive, with very little SBF left for activation by Bck2 (there is very little SBFa1 and SBFa5). As for MBF, only the Bck2-activated form is present, and the remaining MBF units are inactive. Therefore, *cln3Δ* mutants attain a very large size to accumulate enough Bck2 to sequester SBF to an active form (from Whi5) to trigger START^79^ (**Fig. 8A**).

In contrast to the *cln3Δ* mutant, in the *bck2Δ* mutant, most of the active SBF complexes are intact, and only the Swi4B (SBFa5) forms remain (**Fig. 8B**). These cells are therefore only slightly larger than wildtype cells (1.32x WT; **Fig. 8B**). Note that the active SBF/MBF complexes responsible for START in *cln3Δ* and *bck2Δ* mutants complement each other. Hence, for *bck2Δ* to show a significant increase in size (as observed in experiments; 1.3x WT^37^), the contribution from Bck2-activated forms should be significant in wildtype cells. This would automatically result in a smaller size for *cln3Δ* mutants. Therefore, ensuring a large size for *cln3Δ* in our simulations (to match 1.8–2.7x WT^35,51^) would make *bck2Δ* larger than wildtype.

Besides SBF and MBF, in START-BYCC, we assume that homodimeric Swi4 has residual transcriptional activity. This assumption is consistent with the following mutant phenotypes: *swi6Δ* is viable and large, whereas *swi6Δ swi4Δ* is inviable^23^) (**Fig. 8, C–D**). The activation of Swi4 dimers is then brought into question. Here, while the deletion of Cln3 in *swi6Δ* cells has no deleterious effect on their viability^23^ (**Fig. 8E**), the mutant *swi6Δ bck2Δ* is inviable^60^ (**Fig. 8F**). These results suggest that the Swi4 dimers depend on Bck2 for their activation (resulting in Swi4B and SBFa5 in the model; **Fig. 3A**).

#### Mutants pertaining to regulation of MBF

The transcription factor MBF has a large functional overlap with SBF^26^. In START-BYCC, we calibrate the relative importance of SBF and MBF, and their functional overlap based on the relative sizes of several known single and double deletion mutants of Mbp1, Swi4, and Swi6 (**Table S4**). For instance, *swi4Δ* cells (absence of SBF, presence of MBF) are about 1.3–1.5x WT size (**Fig. 8G**), whereas *mbp1Δ* cells (absence of MBF, presence of SBF) are approximately 1.2–1.3x WT size^21^ (**Fig. 8H**). The double mutant is inviable, signifying that either SBF or MBF should be present for the activation of START and the viability of cells (**Fig. 8I**).

To explain the experimentally determined phenotypes of *GAL-WHI5* mutants in START-BYCC, we consider that Whi5 inhibits MBF through non-specific binding in the presence of excess Whi5 (**Fig. 3B**). For example, the lethality of the mutant *GAL-WHI5-12A SWI6-SA4* relies on consideration of this inhibition^47,59^. The double mutant *WHI5-12A SWI6-SA4* is viable and large solely due to the presence of MBF (MBFa in **Fig. 7**) and to a lesser extent due to SBF activated by Bck2 (SBFa2) (**Fig. 7C**). However, the mutant *GAL-WHI5-12A SWI6-SA4* is inviable^59^. While Whi5 binding only to SBF does not explain this scenario, assuming that excess Whi5 can bind to and inhibit MBF accounts for the inviability of *GAL-WHI5-12A SWI6-SA4* mutant (**Fig. 8J**). In vitro studies support this hypothesis of Whi5 inhibition of MBF^47^.

#### Mutants pertaining to the interplay between Whi5, Cln3, and Bck2

Next, we survey the interplay between the activators, Cln3 and Bck2, and inhibitor, Whi5, through a set of relevant mutants. The *whi5Δ* mutant is small in size^47,48^ because, in the absence of Whi5, the SBF complex starts G1 in its uninhibited form, which still carries some residual activity. Thus, in *whi5Δ*, the cells need not wait for the accumulation of Cln3 or Bck2 to activate SBF fully, and therefore, START is advanced sooner than in wildtype cells (**Fig. 8K**).

The large size of *cln3Δ* can be explained by the absence of Cln3-activated SBF and MBF^79^, and dependence on the less active Bck2-activated forms (SBFa5, and MBFa) (**Fig. 8A**). Although SBF (unmodified; SBFa1) and Swi4 dimers (SBFa5) could contribute to the transcription factor pool, with Whi5 present, most of SBF is in a Whi5-bound inhibited complex. In agreement with this notion, additionally deleting Whi5 in *cln3Δ* cells makes them much smaller (closer to WT size), since SBF and Swi4 dimers (to a lesser extent) would now be present.

As discussed previously, Cln3 and Bck2 play mutually complementary roles in promoting the START transition (**Fig. 8, A–B**). This is evident from the lethality of the double mutant *cln3Δ bck2Δ*^49^. In this mutant, both Cln3- and Bck2-activated forms of SBF and MBF are absent, while the inhibitor inhibits any available SBF (**Fig. 8L**). The double mutant can nevertheless be rescued by additionally deleting Whi5 (*cln3Δ bck2Δ whi5Δ*)^47,48^, owing to the availability of unmodified SBF (SBFa1) that is free of inhibition and ready to transcribe (**Fig. 8M**). These mutants are, however, larger than WT cells because they only depend on a less active form of SBF for transcription (due to lack of activators and resultant modifications; light green boxes in **Fig. 2, 3**).

Another set of mutants, *swi6Δ*, *cln3Δ*, and *swi6Δ cln3Δ*, can be understood with the help of the iconic representations shown in **Fig. 8A, C, E**, and the simulations showing the redistribution of SBF and MBF forms. We note from the simulations that the mechanistic reasons explaining the viability of *swi6Δ* (2.25x) and *cln3Δ* (2.20x) are very different, even though they are both viable and large START mutants (**Fig. 8A, C**). Although the *swi6Δ* mutant is viable due to the presence of Swi4 that can be activated by Bck2, the cells are extremely large (2.25x WT) due to the low activity of Swi4. In contrast, the viability of *cln3Δ* is due to MBF activated by Bck2 (**Fig. 8A**). Furthermore, this explanation accounts for the similar size observed in *swi6Δ* and *swi6Δ cln3Δ* (∼2.30x) cells^60^ (**Fig. 8C, E**). Both mutants depend on Bck2 for survival. Supporting evidence for our observation comes from the fact that both of these single mutants are lethal in a *bck2Δ* background^37,42^ (**Fig. 8F, L**). However, the observation that *swi6Δ CLN3-1* is also similar in size to *swi6Δ* and *swi6Δ cln3Δ*, indicates epistasis of Swi6 to Cln3^60^ (**Fig. 8N**). Since Cln3 has secondary effects downstream of SBF (in START-BYCC), the sizes of these three mutants were not quite comparable to other experiments.

The results above suggest that deletion or over-expression of Cln3 would have no additional effect on *swi6Δ*, since the viability of *swi6Δ* only depends on the Swi4 dimer form (SBFa5) activated solely by Bck2 (**Fig. 8, E, N**). However, to pass the START transition, Cln1,2 (resulting from SBF transcription) and Cln3 are needed to i) inactivate Sic1, so that active Clb5,6 (transcribed by MBF genes) can accumulate to trigger DNA synthesis, and ii) inactivate Cdh1 to allow Clb2 to accumulate in preparation for mitosis. Thus, we reparameterized START-BYCC to take into account the lower effect of Cln3 on the cyclin antagonists (while ensuring that all other mutant phenotype emulations continue to hold). This also required changes in other parameters to ensure the viability of the double mutant *cln1Δ cln2Δ* (which depends on Cln3 and Bck2 to inactivate Sic1 for Clb5 to start DNA synthesis). Our current parameter set explains these three mutants’ phenotypes, as well as the large size of *cln1Δ cln2Δ* (**Table S4**). Here, again, we benefit from looking at the complexes present in each of these single, double, or triple mutants using the corresponding representative iconic tables and time-course simulations for different complexes.

Thus, we have parameterized our current model and adjusted the activity/contribution of different complexes such that the time-course simulations match the known experimental phenotypes as much as possible. Of the ∼125+ mutant phenotypes that we simulate, we are able to reproduce most mutants to a high degree of accuracy (**Table S4**). From among the remaining mutants, START-BYCC fails to explain only a few qualitatively, and one quantitatively. We will describe these contradictions in some detail below.

### Model Inconsistencies

While the emulation of a predominant fraction of mutant phenotypes gives much credence to START-BYCC, the few contradictions point out gaps in our current understanding of the budding yeast cell cycle.

#### swi6Δ GAL-WHI5

Whi5 only binds to intact SBF complexes and not to either Swi4 or Swi6 in isolation^48^. Accordingly, in START-BYCC, we do not allow Whi5 to bind to and inhibit the only available active form in *swi6Δ*, the Swi4 complex activated by Bck2. Therefore, contrary to experimental findings^47^, we expect *swi6Δ GAL-WHI5* cells to be similar in size to *swi6Δ* cells (**Fig. S9A**). Such inhibition leads to the presence of free nuclear Whi5 molecules in *swi6Δ* cells, and their inhibition on Swi4 would reduce the propensity for swi6Δ to trigger START, making the single mutant too big to fit the experimental data. It is also likely that Whi5 has non-specific binding to START components, yet unexplored in this model.

#### msn5Δ swi4Δ and msn5Δ swi6Δ

We expect the double mutants involving the export protein, Msn5, and SBF components, *msn5Δ swi4Δ* and *msn5Δ swi6Δ*, to be the same size as *swi4Δ* and *swi6Δ* cells, respectively (**Fig. S9, B–C**). This is because the only direct effect of Msn5 in START-BYCC is through the export of Whi5 and Swi6, which predominantly affects the localization of SBF, and, to a much lesser extent, MBF. In both these double mutants, there is no SBF to begin with. Therefore, additional deletion of Msn5 has no significant effect on either of these single mutants *swi4Δ* and *swi6Δ*. There is, however, a minor difference in size between *swi4Δ* and *msn5Δ swi4Δ*. The single deletion mutant depends only on MBF for its viability, and further deletion of Msn5 results in retaining phosphorylated Whi5 and Swi6 in the nucleus. Although dephosphorylation of these forms occurs in the nucleus, there is a substantial amount of phosphorylated Swi6 that cannot form MBF. The decrease in the amount of net MBF could explain the slightly larger size of *msn5Δ swi4Δ* as compared to the single deletion mutant of *swi4Δ* (**Fig. S9B**). On the other hand, *swi6Δ* only depends on Swi4 for its viability. Therefore, the double mutant *msn5Δ swi6Δ* has the same size as *swi6Δ* cells (**Fig. S9C**). This contradicts the experimental observation that these double mutants are inviable^78^. Clearly, START-BYCC cannot explain this scenario because we do not consider the other diverse effects of Msn5 on the cell, such as the transport of other phosphoproteins like Cdh1^92^. Adding these interactions to the model in a subsequent update could help in explaining these mutants.

#### cln1Δ cln2Δ cdh1Δ and cln1Δ cln2Δ cdh1Δ GAL-CLN2

Two other mutants that START-BYCC currently fails to explain are *cln1Δ cln2Δ cdh1Δ* and *cln1Δ cln2Δ cdh1Δ GAL-CLN2*. Contrary to experimental findings^86^ and simulations from BYCC, the former triple mutant is inviable, while the latter quadruple mutant is viable yet very small in our simulations (**Fig. S9D1, D2**). We suspect that this is due to abnormally high inhibition of CKI by Clb2 in the model, which is an artifact of re-parameterizing the model to fit several other START mutants. We found that lowering the efficiency of Clb2 inhibition on Sic1 indeed rescues the phenotypes, but results in other problems such as the viability of the lethal phenotype *swi4Δ swi6Δ*. The latter mutant does not have functional SBF or MBF, and therefore, no Cln1,2 or Clb5,6 and relies on Clb2 to keep CKIs’ levels low. Therefore, the alteration of these parameters results in the cycling of *swi4Δ swi6Δ* cells. Similarly, lowering the efficiency of Clb2 on the other CKI, Cdc6, results in toggling the inviable mutant *sic1Δ cdh1Δ* to viable. This is because *sic1Δ cdh1Δ* depends on Clb2 to inhibit Cdc6. Lowering the efficiency would result in Cdc6 accumulation and reentry into G1. Therefore, even in this case, it is difficult to keep *cln1Δ cln2Δ cdh1Δ* viable and *sic1Δ cdh1Δ* inviable simultaneously. These scenarios let us posit that the balance and interplay between the mitotic cyclins and CKIs is more complicated than this that additional careful experiments might help resolve.

In *cln1Δ cln2Δ cdh1Δ* mutant, cells are very large at the time of mitotic exit due to *cln1Δ cln2Δ*, and high Clb2 (due to *cdh1Δ* and high mass). High Clb2 inhibits the accumulation of CKI needed for *cdh1Δ* cells to exit from mitosis. Therefore, the model was modified to include additional experimental details on Cdh1’s effect on Cdc20 degradation. Effectively, this results in a reduction of Clb2 levels in the absence of Cdh1. Despite several modifications made in the parameters and wiring of the model, the current model still does not include the effect of Cdh1 on Polo kinase (Cdc5). If we consider these effects, then in the *cdh1Δ* mutant, Polo would be stabilized, causing Cdc14 release and, in turn, resulting in CKI synthesis. Thus, with higher CKI and a lower amount of Clb2, the mutant *cln1Δ cln2Δ cdh1Δ* might be able to exit mitosis in line with observed experiments.

Currently, in START-BYCC, *cln1Δ cln2Δ cdh1Δ GAL-CLN2* is viable but extremely small (**Fig. S9D2**). *GAL-CLN2* inhibits CKI, which increases at the mitotic exit. If the efficiency of CLN2 inhibition on CKI is lowered, then the mutant *cln3Δ* grows too big and dies. This is because MBF active in *cln3Δ* cells makes a small amount of Cln2, and the viability of the mutant relies heavily on Cln2 to inhibit Sic1. This inhibition in *cln3Δ* would result in Clb5 accumulation and DNA synthesis. Therefore, it is quite difficult to achieve simultaneous viability of both *cln1Δ cln2Δ cdh1Δ GAL-CLN2* and cln3Δ. Here again, incorporating the effect of Cdh1 on Polo and Cdc14 in START-BYCC might make the mutant viable due to higher levels of CKI (the viability of *cln1Δ cln2Δ cdh1Δ GAL-CLN2 GAL-SIC1* (**Fig. S9D3**) supports our hypothesis).

We predict that a future version of START-BYCC with a more detailed version of mitotic exit would successfully explain these mutants, as well as several others.

### Model Predictions and Validations

The advantages of our detailed mathematical model extend beyond understanding the molecular mechanisms underlying known mutant phenotypes. It also summarizes, organizes, and reconciles current knowledge about cell cycle regulation, from which we can make testable predictions (see tested predictions here^91^). In this section, we enumerate some of the predictions from START-BYCC (**Fig. S10**) and provide the entire list as part of **Table S4** (highlighted in blue). More of these predictions can be explored in our online simulator: sbmlsimulator.org/simulator/by-start.

#### bck2Δ mbp1Δ

Consider the following cases: deletion of Bck2 and Cln3 in *mbp1Δ* background. Based on the sizes of *cln3Δ* and *bck2Δ* (*cln3Δ* >> *bck2Δ*), we infer that Cln3 is a more efficient activator of SBF and MBF than Bck2. Similarly, based on the sizes of *swi4Δ* and *mbp1Δ*, SBF is more active than MBF. We, therefore, expect the mutant ***bck2Δ mbp1Δ*** to be viable since the major activator of SBF, Cln3, is still present (**Fig. S10A**). In the simulation, we observe that the Cln-activated SBF forms are present (SBFa2>SBFa3, SBFa4, SBFa1) and that they contribute to the viability of the cell in the absence of MBFa and Bck2, in agreement with the validated phenotype in Adames et. al., 2015^91^.

#### bck2Δ mbp1Δ GAL-WHI5

Even though Whi5 is an inhibitor of SBF, *GAL-WHI5* need not always be highly detrimental to the cell. This is because we expect the G1 cyclin (Cln3) and the downstream cyclins that engage in a positive feedback loop (Cln1,2 and Clb5,6) to efficiently phosphorylate the additional Whi5-bound SBF added to the system. We therefore expect that in the mutant ***bck2Δ mbp1Δ GAL-WHI5***, the Cln-activated forms (SBFa2>>SBFa3, SBFa4) would contribute to cell viability (**Fig. S10B**), but the experimental phenotype shows that this mutant is G1 arrested^91^. Since we assign full activity to Whi5-bound SBF that is phosphorylated on Swi6 by Cln kinases, SBFa3 and SBFa4 will contribute to overall SBF activity, and the triple mutant will be viable. Similar to this triple mutant, the double mutants ***mbp1Δ GAL-WHI5*** and ***mbp1Δ GAL-WHI5-12A*** are also viable despite excessive Whi5 in the system because we expect that they are converted to active Whi5-bound SBF forms (SBFa2, SBFa3, SBFa4 in the former case and SBFa3 in the latter non-phosphorylable case) (**Fig. S10, H-I**). These forms are present at sufficiently high levels to ensure viability. Very high doses of Whi5 might saturate the kinases in the system, resulting in cell inviability. For other moderate increases in Whi5 or Whi5-12A levels, we expect the aforementioned response in the background of *mbp1Δ* cells.

#### cln3Δ mbp1Δ, cln3Δ mbp1Δ swi6Δ, and cln3Δ mbp1Δ whi5Δ

In START-BYCC, we would expect the double mutant ***cln3Δ mbp1Δ*** to be viable and large (with a delayed START for cycle 1). This is because, with no MBF, only SBF is available as the activator of Cln1,2 and Clb5,6, which delays the cycle and results in a larger cell size. The triple deletion mutant cln3Δ mbp1Δ swi6Δ is predicted to be viable and large with neither SBF nor MBF. Only swi4 dimers exist (only SBFa5 is present). Since swi4 dimers can only be activated by Bck2, this delays the cell cycle, enhancing cell growth (**Fig. S10D**; almost the same size as cln3Δ mbp1Δ). We predict another triple deletion mutant, cln3Δ mbp1Δ whi5Δ, to reduce the cell size (from cln3Δ mbp1Δ). As seen in **Fig. S10E**, the size of cells is smaller because whi5 is not present, so there is no inactivation, and SBF activations happen much earlier. All of our simulations here agree with the experimentally validated phenotypes^91^.

#### Rescue of cln3Δ swi4Δ by whi5Δ, GAL-BCK2, and whi5Δ sic1Δ

The double mutant ***cln3Δ swi4Δ*** is inviable because the only active form present is MBF activated by Clns or Bck2^93^. Our simulations show that an additional deletion of Whi5 (***cln3Δ swi4Δ whi5Δ***) can rescue the phenotype, as MBF is relieved in the absence of Whi5 (**Fig. S10F**), in line with experimental validation^91^. The mutant ***cln3Δ swi4Δ whi5Δ*** is predicted to be further rescued by the additional deletion of Sic1 due to early relieving of inhibition on active MBF (this quadruple mutant with *sic1Δ* is smaller than the parent triple deletion). Similarly, overexpression of Bck2 also rescues the double mutant that lacks SBF, because there is sufficient Bck2-activated MBF (***cln3Δ swi4Δ GAL-BCK2***, MBFa forms in **Fig. S10G**) to drive forward the cell cycle.

#### Rescue of swi4Δ swi6Δ by GAL-CLB5 and GAL-CLN2 (not by GAL-CLN3)

Both the double mutants *swi4Δ swi6Δ and swi4Δ mbp1Δ* are inviable in our simulations due to the absence of SBF/MBF, in strong agreement with experimental observations^21,23^ (**Fig. 8**). The logical prediction from the model follows that an additional deletion of Whi5 would not rescue the double deletion of Swi4 and Swi6; the triple deletion *swi4Δ swi6Δ whi5Δ* is inviable (**Table 1**). These cells do not have any SBF or MBF, and hence Whi5 does not have any stoichiometric partner to bind with and inhibit. However, ***swi4Δ swi6Δ*** can be rescued by ***GAL-CLB5*** and ***GAL-CLN2*** (but not by ***GAL-CLN3***), since CKIs are still high enough to prevent triggering DNA synthesis (**Table S4**). Similarly, *GAL-CLB5* and *GAL-CLN2* can also rescue *bck2Δ swi6Δ* (but not ***GAL-CLN3***) (**Table S4**).

#### Rescue of swi4Δ swi6Δ and bck2Δ swi6Δ by SWI6-SA4

Similar to rescue by multi-copy Swi6^49^, we also expect the lethal mutants ***swi4Δ swi6Δ* and *bck2Δ swi6Δ* to be rescued by *SWI6-SA4***, since MBF would be available again in the former case, and both SBF (although non-phosphorylable at Swi6) and MBF available in the latter case, to rescue the lethal phenotype (**Table S4**).

## Conclusion

The cell cycle circuit is the central machinery of the cell, driving its controlled growth and division. The START transition in the budding yeast cell cycle serves as an important checkpoint to ensure the absence of inhibitory signals and the growth of the cell to a critical size before commitment to cell division. In this study, we developed a detailed mathematical model of the START transition (START-BYCC), incorporating molecular details of this process, including signaling/regulatory interactions, phosphorylation states, and subcellular localization of key START proteins. The model recapitulates the experimentally observed phenotypes of over 120 mutants pertaining to START and other important cell cycle phases (several representative mutants can be accessed here: sbmlsimulator.org/simulator/by-start). START-BYCC also captures the subcellular localization and translocation of the transcription factor, SBF, and provides a mechanism for size control.

Most importantly, the analysis we present here establishes a foundation for novel hypotheses and precise predictions on the mechanisms of the START transition that can be verified experimentally (including some successful validations highlighted recently^91^). Secondly, the nutritional conditions of a cell greatly influence its size control mechanism, with several recently discovered molecular players mediating this effect. These interactions, and any newer findings, can be dovetailed into START-BYCC to generalize the existing mechanism for size control. Future work will entail appending detailed current models of other phases and events of the budding yeast cell cycle, including DNA damage response, spindle assembly checkpoint, and mitotic exit. START-BYCC provides an opportune setting to incorporate these modules (the model is available here for any future extensions and expansions: github.com/jravilab/start-bycc). Finally, this detailed model for budding yeast offers a path towards dealing with the enormous complexity of the cell cycle mechanism in high eukaryotes, including humans allowing us to uncover the fundamental principles and regulatory bottlenecks that are essential to delineate the underlying system dynamics. The evolutionary conservation of the molecular circuitry underlying the START transition in the yeast cell cycle and R-point in mammalian counterparts supports this view, and our detailed model would help these parallel eukaryotic studies in bridging our gaps in understanding these complex systems. Modeling the R-point not only offers an understanding of the workings of a healthy cell cycle, but also tenders insights into its characteristic deregulation in diseases like cancer.

## Materials and Methods

### Current START-BYCC model

A large number of experimental findings were reconciled to establish a detailed wiring diagram for the START transition (**Fig. 2, 3**) in conjunction with the BYCC model for the budding yeast cell cycle. Several new aspects of the regulation of START have been revealed since 2004, including the role of the protein Whi5, which is a stoichiometric inhibitor of the transcription factor SBF. The START module of the model has been described in detail under the ‘Results’ section (**Fig. 2, 3**). The remaining cell cycle model (from START to mitotic exit) has been used almost as-is from the BYCC model (except for changes in equations of START proteins like Cln3, Bck2, Cln2, and Clb5). The wiring diagram was converted to chemical reactions (mass-action) and algebraic relations, which were then translated to ordinary differential equations and algebraic equations. The codes, wiring diagrams, equations, and parameters will be available via GitHub (https://github.com/jravilab/start-bycc) and our easy-to-use online simulator (http://sbmlsimulator.org/simulator/by-start).

### Modifications from the BYCC model

We started with the well-established budding yeast cell cycle (BYCC) mathematical model from Chen et al., 2004. The major modifications are associated with the START transition of the cell cycle. The simplified Goldbeter-Koshland switch, previously used to describe the activation of SBF and MBF in the BYCC model, has been replaced with the substantially detailed model presented in **Fig. 2, 3**. This new model delineates several important phosphorylation and transportation events occurring at START. Cellular compartments — nucleus and cytoplasm — are explicitly modeled to track localization. They are assumed to have a constant volume ratio of 1:4 throughout the cell cycle^88,94^. The species that move across compartments are scaled with the corresponding volumes to make the concentrations in the equations consistent.

Additionally, in the current version of the model, we have removed mass dependence from cyclins such as Cln1,2 represented as a single variable in the model, Cln2, since they are not concentrated in the nucleus and are distributed roughly throughout the cell. We have retained mass dependence only on Cln3 (through Ydj1 as the size sensor and on Cln3 level in the nucleus), Bck2, Clb5, and Clb2 in order to explain the complicated contributions of these species to size control and/or their nuclear localization. We also use a Hill function to model Cln activation at START (equation of Vpcln in Supplementary text) due to the existence of cooperativity^83^, and to avoid the complexity and additional intermediates that would stem from considering multi-site phosphorylation of Whi5, Swi6, and Swi4. We tested a simpler version of the START module modeled with multi-site phosphorylation and confirmed that it showed a behavior similar to the Hill function-based model. Thus, we concluded that the Hill function is a reasonable approximation and phenomenological abstraction of multi-site phosphorylation in our case, and used it in START-BYCC.

### Equations and Parameters

The equations and parameters used for our simulations are listed under the Supplementary text, and the files have been provided in the following formats: ODE, SBML, and PET, including the parameter set used. We have also listed all changes made to the initial conditions and parameters in simulating specific mutants, along with key assumptions (**Table S2**). The codes, equations, and parameters will be available via GitHub (github.com/jravilab/start-bycc) and our online simulator (sbmlsimulator.org/simulator/by-start).

### Simulations

We used JigCell^95^ to build START-BYCC by defining biochemical reactions (or interactions), algebraic rules, conservation relations, discrete events at the end of the cell cycle, and multiple compartments. All reactions, equations, and conservation relations were verified manually. Parameter Estimation Toolkit (PET) was used to run all our simulations (using LSODAR)^96^ for wildtype and mutants by defining the specified conditions in **Table S3**. The software offers the option of simultaneously running simulations for different conditions (termed ‘simulation runs’) and for several sets of rate constants (termed ‘basal sets’). XPPAUT^97^ (sites.pitt.edu/~phase/bard/bardware/xpp/xpp.html) was also used to check our simulations numerically (for wildtype cells).

## Supplementary Material

Available as a separate PDF.

## Acknowledgments

First and foremost, we would like to thank John Tyson (now emeritus) for invaluable advice, discussion, and support during the early stages of the project. We would like to thank Kartik Subramanian, Debashis Barik, Sandip Kar, Tongli Zhang, Teeraphan Laomettachit, Vandana Sreedharan, and Arjun Krishnan for providing the authors with several iterations of constructive feedback. We are also extremely grateful to Krishnan Raghunathan, Arjun Krishnan, and Emily Meyer for their detailed comments on the manuscript. We have benefited from several conversations and diverse mutant phenotypes and challenges brought to us by researchers in the budding yeast cell cycle field that helped us fine-tune the model, including Jan Skotheim, Stefano Di Talia, and Fred Cross. We are also grateful to Jean Peccoud and their team for the first set of experimental validations.

## Author Contributions

JR conceived and designed the study; JR and KS acquired the data, performed all the analyses, and made the figures and tables; JR chalked the wiring diagram and mathematical model, parameterized the model, and ran all the initial simulations; JZ built the Parameter Estimation Toolkit (that was mostly used to run and parameterize the model and simulations), and build the online simulator used to visualize the dynamics in this manuscript; JR wrote the first draft of the manuscript; JR and KS revised the manuscript.

## Data Availability and Reuse

All the simulation data and visualizations (for wildtype and 100s of mutants) are available in our interactive online simulator: sbmlsimulator.org/simulator/by-start. The web simulator is licensed under the MIT license. Our code, differential equation model, and parameters are available via GitHub: github.com/jravilab/start-bycc.

## References

1. Hartwell, L. H., Culotti, J. & Reid, B. Genetic Control of the Cell-Division Cycle in Yeast, I. Detection of Mutants. Proc. Natl. Acad. Sci. 66, 352–359 (1970).

2. Nurse, P., Thuriaux, P. & Nasmyth, K. Genetic control of the cell division cycle in the fission yeast Schizosaccharomyces pombe. Mol. Gen. Genet. MGG 146, 167–178 (1976).

3. Evans, T., Rosenthal, E. T., Youngblom, J., Distel, D. & Hunt, T. Cyclin: A protein specified by maternal mRNA in sea urchin eggs that is destroyed at each cleavage division. Cell 33, 389–396 (1983).

4. Murray, A.W. & Hunt, T. The Cell Cycle: An Introduction. (1993).

5. Nurse, P. Universal control mechanism regulating onset of M-phase. Nature 344, 503–508 (1990).

6. Hartwell, L. H., Culotti, J., Pringle, J. R. & Reid, B. J. Genetic Control of the Cell Division Cycle in Yeast: A model to account for the order of cell cycle events is deduced from the phenotypes of yeast mutants. Science 183, 46–51 (1974).

7. Johnston, G., Pringle, J. & Hartwell, L. Coordination of growth with cell division in the yeast. Exp. Cell Res. 105, 79–98 (1977).

8. Johnston, G. C., Ehrhardt, C. W., Lorincz, A. & Carter, B. L. Regulation of cell size in the yeast Saccharomyces cerevisiae. J. Bacteriol. 137, 1–5 (1979).

9. Jagadish, M. N. & Carter, B. L. A. Genetic control of cell division in yeast cultured at different growth rates. Nature 269, 145–147 (1977).

10. Jorgensen, P. A dynamic transcriptional network communicates growth potential to ribosome synthesis and critical cell size. Genes Dev. 18, 2491–2505 (2004).

11. Lorincz, A. & Carter, B. L. A. Control of Cell Size at Bud Initiation in Saccharomyces cerevisiae. J. Gen. Microbiol. 113, 287–295 (1979).

12. Chang, F. & Herskowitz, I. Identification of a gene necessary for cell cycle arrest by a negative growth factor of yeast: FAR1 is an inhibitor of a G1 cyclin, CLN2. Cell 63, 999–1011 (1990).

13. Wittenberg, C. & Reed, S. I. Plugging it in: signaling circuits and the yeast cell cycle. Curr. Opin. Cell Biol. 8, 223–230 (1996).

14. Peter, M. Joining the complex: Cyclin-dependent kinase inhibitory proteins and the cell cycle. Cell 79, 181–184 (1994).

15. Rupeš, I. Checking cell size in yeast. Trends Genet. 18, 479–485 (2002).

16. Mendenhall, M. D. & Hodge, A. E. Regulation of Cdc28 Cyclin-Dependent Protein Kinase Activity during the Cell Cycle of the Yeast *Saccharomyces cerevisiae*. Microbiol. Mol. Biol. Rev. 62, 1191–1243 (1998).

17. Bosl, W. J. & Li, R. Mitotic-exit control as an evolved complex system. Cell 121, 325–333 (2005).

18. Sullivan, M. & Morgan, D. O. Finishing mitosis, one step at a time. Nat. Rev. Mol. Cell Biol. 8, 894–903 (2007).

19. Chen, Y. et al. The N Terminus of the Centromere H3-Like Protein Cse4p Performs an Essential Function Distinct from That of the Histone Fold Domain. Mol. Cell. Biol. 20, 7037–7048 (2000).

20. Chen, K. C. et al. Integrative Analysis of Cell Cycle Control in Budding Yeast. Mol. Biol. Cell 15, 3841–3862 (2004).

21. Koch, C., Moll, T., Neuberg, M., Ahorn, H. & Nasmyth, K. A role for the transcription factors Mbp1 and Swi4 in progression from G1 to S phase. Science 261, 1551–1557 (1993).

22. Gallego, C. The Cln3 cyclin is down-regulated by translational repression and degradation during the G1 arrest caused by nitrogen deprivation in budding yeast. EMBO J. 16, 7196–7206 (1997).

23. Nasmyth, K. & Dirick, L. The role of SWI4 and SWI6 in the activity of G1 cyclins in yeast. Cell 66, 995–1013 (1991).

24. Irvali, D. et al. When yeast cells change their mind: cell cycle “Start” is reversible under starvation. EMBO J. 42, e110321 (2023).

25. Barik, D., Baumann, W. T., Paul, M. R., Novak, B. & Tyson, J. J. A model of yeast cell-cycle regulation based on multisite phosphorylation. Mol. Syst. Biol. 6, 405 (2010).

26. Bean, J. M., Siggia, E. D. & Cross, F. R. High Functional Overlap Between MluI Cell-Cycle Box Binding Factor and Swi4/6 Cell-Cycle Box Binding Factor in the G1/S Transcriptional Program in *Saccharomyces cerevisiae*. Genetics 171, 49–61 (2005).

27. Mendenhall, M. An inhibitor of p34CDC28 protein kinase activity from Saccharomyces cerevisiae. Science 259, 216–219 (1993).

28. Verma, R., Feldman, R. M. & Deshaies, R. J. SIC1 is ubiquitinated in vitro by a pathway that requires CDC4, CDC34, and cyclin/CDK activities. Mol. Biol. Cell 8, 1427–1437 (1997).

29. Amon, A. Regulation of B-type cyclin proteolysis by Cdc28-associated kinases in budding yeast. EMBO J. 16, 2693–2702 (1997).

30. Rieder, C. L. Mitosis in vertebrates: the G2/M and M/A transitions and their associated checkpoints. Chromosome Res. 19, 291–306 (2011).

31. Siegmund, R. F. & Nasmyth, K. A. The Saccharomyces cerevisiae Start-specific transcription factor Swi4 interacts through the ankyrin repeats with the mitotic Clb2/Cdc28 kinase and through its conserved carboxy terminus with Swi6. Mol. Cell. Biol. 16, 2647–2655 (1996).

32. Amon, A., Tyers, M., Futcher, B. & Nasmyth, K. Mechanisms that help the yeast cell cycle clock tick: G2 cyclins transcriptionally activate G2 cyclins and repress G1 cyclins. Cell 74, 993–1007 (1993).

33. de Bruin, R. A. M. et al. DNA replication checkpoint promotes G1-S transcription by inactivating the MBF repressor Nrm1. Proc. Natl. Acad. Sci. 105, 11230–11235 (2008).

34. Amon, A. The spindle checkpoint. Curr. Opin. Genet. Dev. 9, 69–75 (1999).

35. Jaspersen, S. L., Charles, J. F. & Morgan, D. O. Inhibitory phosphorylation of the APC regulator Hct1 is controlled by the kinase Cdc28 and the phosphatase Cdc14. Curr. Biol. 9, 227–236 (1999).

36. Visintin, R. et al. The Phosphatase Cdc14 Triggers Mitotic Exit by Reversal of Cdk-Dependent Phosphorylation. Mol. Cell 2, 709–718 (1998).

37. Cooper, K. F., Mallory, M. J., Egeland, D. B., Jarnik, M. & Strich, R. Ama1p is a meiosis-specific regulator of the anaphase promoting complex/cyclosome in yeast. Proc. Natl. Acad. Sci. 97, 14548–14553 (2000).

38. Oelschlaegel, T. et al. The Yeast APC/C Subunit Mnd2 Prevents Premature Sister Chromatid Separation Triggered by the Meiosis-Specific APC/C-Ama1. Cell 120, 773–788 (2005).

39. Penkner, A. M., Prinz, S., Ferscha, S. & Klein, F. Mnd2, an Essential Antagonist of the Anaphase-Promoting Complex during Meiotic Prophase. Cell 120, 789–801 (2005).

40. Alberts, B. et al. Molecular biology of the cell. (New York: Garland Science, 2002).

41. Hancioglu, B. & Tyson, J. J. A mathematical model of mitotic exit in budding yeast: the role of Polo kinase. PloS One 7, e30810 (2012).

42. Novak, B. & Tyson, J. J. Numerical analysis of a comprehensive model of M-phase control in Xenopus oocyte extracts and intact embryos. J. Cell Sci. 106, 1153–1168 (1993).

43. Azzam, R. et al. Phosphorylation by Cyclin B-Cdk Underlies Release of Mitotic Exit Activator Cdc14 from the Nucleolus. Science 305, 516–519 (2004).

44. Queralt, E., Lehane, C., Novak, B. & Uhlmann, F. Downregulation of PP2ACdc55 Phosphatase by Separase Initiates Mitotic Exit in Budding Yeast. Cell 125, 719–732 (2006).

45. Manzoni, R. et al. Oscillations in Cdc14 release and sequestration reveal a circuit underlying mitotic exit. J. Cell Biol. 190, 209–222 (2010).

46. Kraikivski, P., Chen, K. C., Laomettachit, T., Murali, T. M. & Tyson, J. J. From START to FINISH: computational analysis of cell cycle control in budding yeast. Npj Syst. Biol. Appl. 1, 15016 (2015).

47. Costanzo, M. et al. CDK Activity Antagonizes Whi5, an Inhibitor of G1/S Transcription in Yeast. Cell 117, 899–913 (2004).

48. de Bruin, R. A. M., McDonald, W. H., Kalashnikova, T. I., Yates, J. & Wittenberg, C. Cln3 Activates G1-Specific Transcription via Phosphorylation of the SBF Bound Repressor Whi5. Cell 117, 887–898 (2004).

49. Wijnen, H. & Futcher, B. Genetic Analysis of the Shared Role of CLN3 and BCK2 at the G1-S Transition in Saccharomyces cerevisiae. Genetics 153, 1131–1143 (1999).

50. Geymonat, M., Spanos, A., Wells, G. P., Smerdon, S. J. & Sedgwick, S. G. Clb6/Cdc28 and Cdc14 Regulate Phosphorylation Status and Cellular Localization of Swi6. Mol. Cell. Biol. 24, 2277–2285 (2004).

51. Johnson, A. & Skotheim, J. M. Start and the restriction point. Curr. Opin. Cell Biol. 25, 717–723 (2013).

52. Dryja, T. P., Friend, S. & Weinberg, R. A. Genetic sequences that predispose to retinoblastoma and osteosarcoma. Symp. Fundam. Cancer Res. 39, 115–119 (1986).

53. Friend, S. H. et al. A human DNA segment with properties of the gene that predisposes to retinoblastoma and osteosarcoma. Nature 323, 643–646 (1986).

54. Fung, Y. K. et al. Structural evidence for the authenticity of the human retinoblastoma gene. Science 236, 1657–1661 (1987).

55. Lee, W. H. et al. Human retinoblastoma susceptibility gene: cloning, identification, and sequence. Science 235, 1394–1399 (1987).

56. Mittnacht, S. & Weinberg, R. A. G1/S phosphorylation of the retinoblastoma protein is associated with an altered affinity for the nuclear compartment. Cell 65, 381–393 (1991).

57. Talia, S. D., Skotheim, J. M., Bean, J. M., Siggia, E. D. & Cross, F. R. The effects of molecular noise and size control on variability in the budding yeast cell cycle. Nature 448, 947–951 (2007).

58. Sidorova, J. M., Mikesell, G. E. & Breeden, L. L. Cell cycle-regulated phosphorylation of Swi6 controls its nuclear localization. Mol. Biol. Cell 6, 1641–1658 (1995).

59. Wagner, A., Grillitsch, K., Leitner, E. & Daum, G. Mobilization of steryl esters from lipid particles of the yeast Saccharomyces cerevisiae. Biochim. Biophys. Acta BBA - Mol. Cell Biol. Lipids 1791, 118–124 (2009).

60. Wijnen, H., Landman, A. & Futcher, B. The G _1_ Cyclin Cln3 Promotes Cell Cycle Entry via the Transcription Factor Swi6. Mol. Cell. Biol. 22, 4402–4418 (2002).

61. Brown, V. D., Phillips, R. A. & Gallie, B. L. Cumulative Effect of Phosphorylation of pRB on Regulation of E2F Activity. Mol. Cell. Biol. 19, 3246–3256 (1999).

62. Tyson, J. J., Chen, K. & Novak, B. Network dynamics and cell physiology. Nat. Rev. Mol. Cell Biol. 2, 908–916 (2001).

63. Tyson, J. & Sachsenmaier, W. Is nuclear division in Physarum controlled by a continuous limit cycle oscillator? J. Theor. Biol. 73, 723–738 (1978).

64. Goldbeter, A. & Koshland, D. E. An amplified sensitivity arising from covalent modification in biological systems. Proc. Natl. Acad. Sci. 78, 6840–6844 (1981).

65. Barberis, M., Klipp, E., Vanoni, M. & Alberghina, L. Cell Size at S Phase Initiation: An Emergent Property of the G1/S Network. PLoS Comput. Biol. 3, e64 (2007).

66. Barik, D., Ball, D. A., Peccoud, J. & Tyson, J. J. A Stochastic Model of the Yeast Cell Cycle Reveals Roles for Feedback Regulation in Limiting Cellular Variability. PLOS Comput. Biol. 12, e1005230 (2016).

67. Chandler-Brown, D., Schmoller, K. M., Winetraub, Y. & Skotheim, J. M. The Adder Phenomenon Emerges from Independent Control of Pre- and Post-Start Phases of the Budding Yeast Cell Cycle. Curr. Biol. 27, 2774–2783.e3 (2017).

68. Heldt, F. S., Lunstone, R., Tyson, J. J. & Novák, B. Dilution and titration of cell-cycle regulators may control cell size in budding yeast. PLOS Comput. Biol. 14, e1006548 (2018).

69. Dorsey, S. et al. G1/S Transcription Factor Copy Number Is a Growth-Dependent Determinant of Cell Cycle Commitment in Yeast. Cell Syst. 6, 539–554.e11 (2018).

70. Münzner, U., Klipp, E. & Krantz, M. A comprehensive, mechanistically detailed, and executable model of the cell division cycle in Saccharomyces cerevisiae. Nat. Commun. 10, 1308 (2019).

71. Laomettachit, T., Chen, K. C., Baumann, W. T. & Tyson, J. J. A Model of Yeast Cell-Cycle Regulation Based on a Standard Component Modeling Strategy for Protein Regulatory Networks. PLOS ONE 11, e0153738 (2016).

72. Laomettachit, T., Kraikivski, P. & Tyson, J. J. A continuous-time stochastic Boolean model provides a quantitative description of the budding yeast cell cycle. Sci. Rep. 12, 20302 (2022).

73. Zhang, T., Schmierer, B. & Novák, B. Cell cycle commitment in budding yeast emerges from the cooperation of multiple bistable switches. Open Biol. 1, 110009 (2011).

74. Li, W., Yi, M. & Zou, X. Mathematical modeling reveals the mechanisms of feedforward regulation in cell fate decisions in budding yeast. Quant. Biol. 3, 55–68 (2015).

75. Baetz, K. & Andrews, B. Regulation of Cell Cycle Transcription Factor Swi4 through Auto-Inhibition of DNA Binding. Mol. Cell. Biol. 19, 6729–6741 (1999).

76. Vergés, E., Colomina, N., Garí, E., Gallego, C. & Aldea, M. Cyclin Cln3 Is Retained at the ER and Released by the J Chaperone Ydj1 in Late G1 to Trigger Cell Cycle Entry. Mol. Cell 26, 649–662 (2007).

77. Ghaemmaghami, S. et al. Global analysis of protein expression in yeast. Nature 425, 737–741 (2003).

78. Queralt, E. & Igual, J. C. Cell Cycle Activation of the Swi6p Transcription Factor Is Linked to Nucleocytoplasmic Shuttling. Mol. Cell. Biol. 23, 3126–3140 (2003).

79. Dirick, L., Böhm, T. & Nasmyth, K. Roles and regulation of Cln-Cdc28 kinases at the start of the cell cycle of Saccharomyces cerevisiae. EMBO J. 14, 4803–4813 (1995).

80. de Bruin, R. A. M. et al. Constraining G1-Specific Transcription to Late G1 Phase: The MBF-Associated Corepressor Nrm1 Acts via Negative Feedback. Mol. Cell 23, 483–496 (2006).

81. Brewster, N. K., Val, D. L., Walker, M. E. & Wallace, J. C. Regulation of Pyruvate Carboxylase Isozyme (PYC1, PYC2) Gene Expression in Saccharomyces cerevisiae during Fermentative and Nonfermentative Growth. Arch. Biochem. Biophys. 311, 62–71 (1994).

82. Charvin, G., Cross, F. R. & Siggia, E. D. Forced periodic expression of G1 cyclins phase-locks the budding yeast cell cycle. Proc. Natl. Acad. Sci. U. S. A. 106, 6632–6637 (2009).

83. Charvin, G., Oikonomou, C., Siggia, E. D. & Cross, F. R. Origin of Irreversibility of Cell Cycle Start in Budding Yeast. PLoS Biol. 8, e1000284 (2010).

84. Lu, Y. & Cross, F. Mitotic exit in the absence of separase activity. Mol. Biol. Cell 20, 1576–1591 (2009).

85. Lara-Gonzalez, P., Westhorpe, F. G. & Taylor, S. S. The spindle assembly checkpoint. Curr. Biol. CB 22, R966–980 (2012).

86. Cross, F. R., Archambault, V., Miller, M. & Klovstad, M. Testing a Mathematical Model of the Yeast Cell Cycle. Mol. Biol. Cell 13, 52–70 (2002).

87. Miller, M. E. & Cross, F. R. Mechanisms Controlling Subcellular Localization of the G _1_ Cyclins Cln2p and Cln3p in Budding Yeast. Mol. Cell. Biol. 21, 6292–6311 (2001).

88. Jorgensen, P. et al. The Size of the Nucleus Increases as Yeast Cells Grow. Mol. Biol. Cell 18, 3523–3532 (2007).

89. Lord, P. G. & Wheals, A. E. Asymmetrical division of Saccharomyces cerevisiae. J. Bacteriol. 142, 808–818 (1980).

90. Taberner, F. J., Quilis, I. & Igual, J. C. Spatial regulation of the Start repressor Whi5. Cell Cycle 8, 3013–3022 (2009).

91. Adames, N. R. et al. Experimental testing of a new integrated model of the budding yeast S tart transition. Mol. Biol. Cell 26, 3966–3984 (2015).

92. Jaquenoud, M. Cell cycle-dependent nuclear export of Cdh1p may contribute to the inactivation of APC/CCdh1. EMBO J. 21, 6515–6526 (2002).

93. Ferrezuelo, F., Colomina, N., Futcher, B. & Aldea, M. The transcriptional network activated by Cln3 cyclin at the G1-to-S transition of the yeast cell cycle. Genome Biol. 11, R67 (2010).

94. Larson, D. R., Zenklusen, D., Wu, B., Chao, J. A. & Singer, R. H. Real-Time Observation of Transcription Initiation and Elongation on an Endogenous Yeast Gene. Science 332, 475–478 (2011).

95. Vass, M. et al. The JigCell Model Builder and Run Manager. Bioinformatics 20, 3680–3681 (2004).

96. Zwolak, J. W., Tyson, J. J. & Watson, L. T. Parameter Estimation for a Mathematical Model of the Cell Cycle in Frog Eggs. J. Comput. Biol. 12, 48–63 (2005).

97. Ermentrout, B. Simulating, Analyzing, and Animating Dynamical Systems: A Guide to XPPAUT for Researchers and Students. (Society for Industrial and Applied Mathematics, 2002). doi:10.1137/1.9780898718195.

