## Supplementary Material: Tables, Figures, Text for "Modeling the START transition in the budding yeast cell cycle"

**Data Availability and Reuse** **1**
**Supplementary Figures** **2**
Figure S1. Complex formation & promoter binding. 2
Figure S2. SBF regulation in wildtype cells. 3
Figure S3. SBF inactivation by Clbs. 4
Figure S4. Cartoon of Cln3 regulation by Ydj1/Ssa1. 5
Figure S5. Duration of daughter cycle times as function of mass doubling time. 6
Figure S6. Localization of different monomers. 7
Figure S7. Importance of the transport protein, Msn5. 8
Figure S8. Non-phosphorylable mutants. 9
Figure S9. Model contradictions. 13
Figure S10. Few model predictions and validations. 14
**Supplementary Tables** **19**
Table S1. Functions of different Cyclins in Budding Yeast cell cycle 19
Table S2. Description, abundance, regulation and localization of START components in the model; List of Key Assumptions 20
Table S3.Modifications in Parameter & Initial conditions corresponding to mutants. (Mutants exclusive to the current model are emphasized in bold). 24
Table S4. START mutants 26
**Supplementary Text** **30**
List of Abbreviations 30
Equations, Parameters and Initial Conditions 30
**License** **31**
**References** **32**

#### Data Availability and Reuse

All the simulation data and visualizations (for wildtype and 100s of mutants) are available in our interactive online simulator: sbmlsimulator.org/simulator/by-start. The web simulator is licensed under the MIT license. Our code, differential equation model, and parameters are available via GitHub: github.com/jravilab/start-bycc.

#### Supplementary Figures

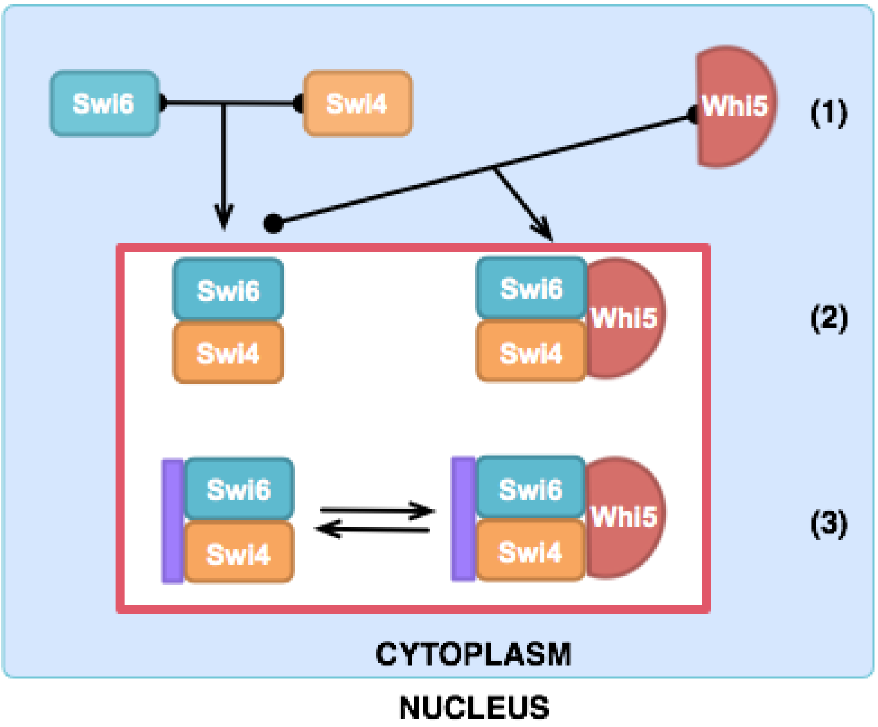


Figure S1. Complex formation & promoter binding.

Initial flow of events in G1 starting from (1) monomers. The top panel is the core model for SBF activation and inactivation as appeared in **Figure 2A**. The red box containing reactions involving in (2) the complex formation and (3) promoter binding is expanded and shown in the lower panel.

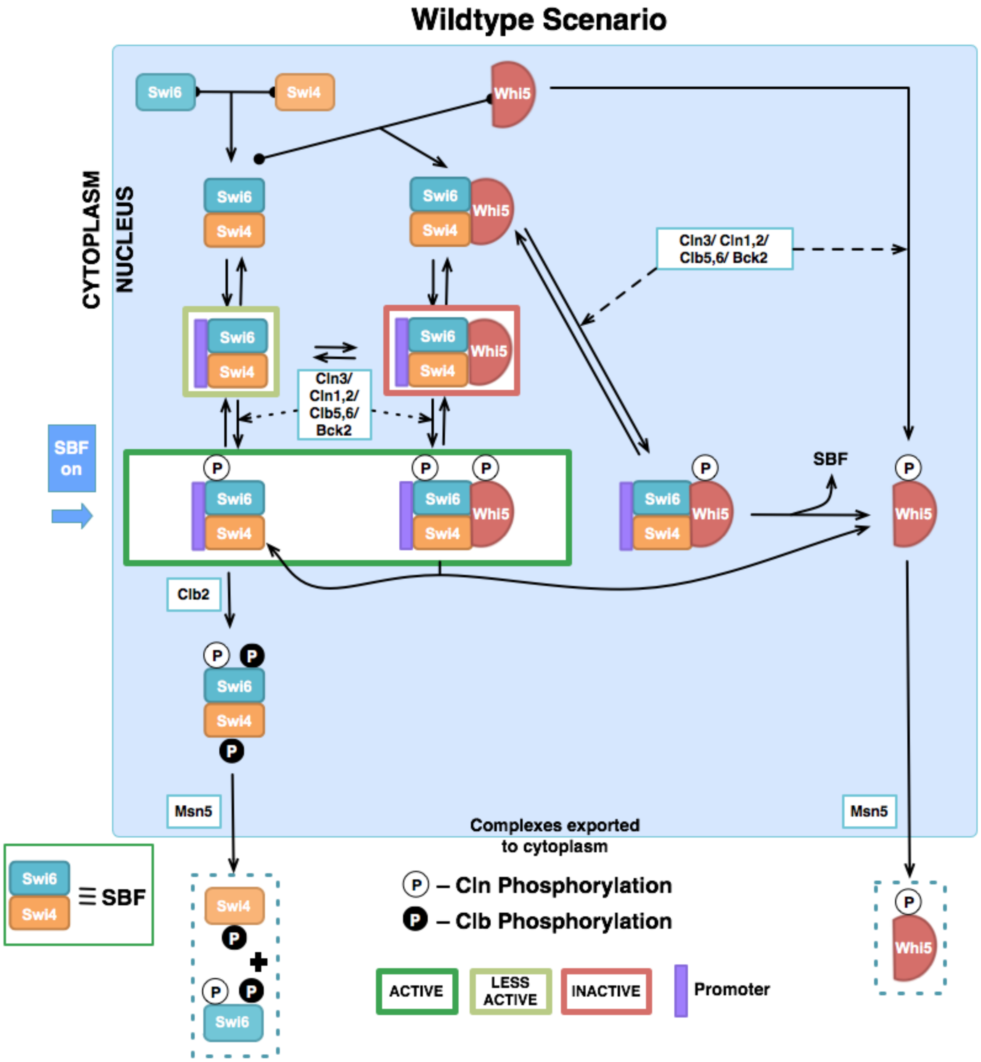


Figure S2. SBF regulation in wildtype cells.

(i) SBF activation and Whi5 export. In the figure, the cells start with monomers, which proceed to form complexes and bind to the promoter. In WT cells, both Swi6 (P-form) and Whi5 get phosphorylated by Clns (Cln3, Cln1,2 and Clb5,6) and activated by Bck2. We assume that this doubly phosphorylated form is unstable and dissociates to yield active SBF (with Swi6 phosphorylated) and phosphorylated Whi5 that is free to move to the cytoplasm with the help of export protein, Msn5. (ii) SBF inactivation and export. SBF gets inactivated by two sets of Clb phosphorylations (black-filled circles): Q-form on S160 site of Swi6 by Clb5,6 and Clb1,2, and on Swi4 by Clb1,2, leading to dissociation of SBF from promoter. Msn5 recognizes the Q-form of Swi6 phosphorylation (that is in complex with Swi4) for export to the cytoplasm. Complex dissociates in the cytoplasm soon after export. Phosphate groups are indicated as white-filled circles (activatory phosphorylations) or with black-filled circles (inhibitory phosphorylations).

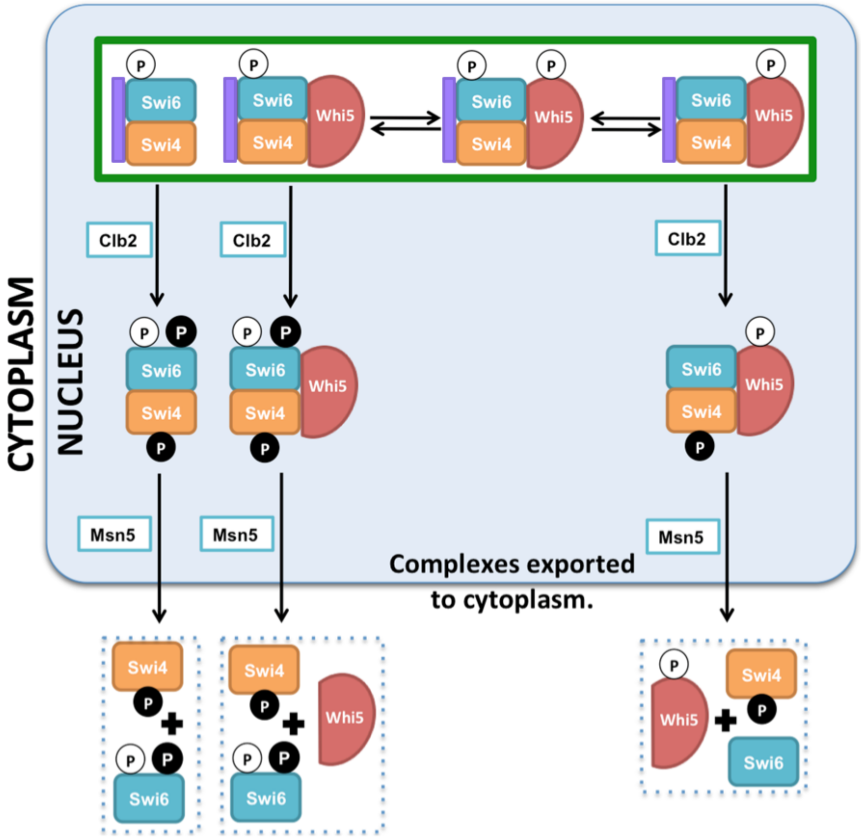


Figure S3. SBF inactivation by Clbs.

We assume that either Clb phosphorylation on Swi6 (S160) (Q-form), or Cln phosphorylation on Whi5 is necessary for recognition by transport protein, Msn5, and subsequent export to the cytoplasm. Additionally, we assume that in promoter-bound complexes, Clb phosphorylation of Swi4 is necessary and sufficient for the complexes to dissociate from the promoter necessary to turn off gene transcription and to facilitate SBF export. Similar to SBF inactivation in Figure S2, SBF complexes dissociate in the cytoplasm soon after export.

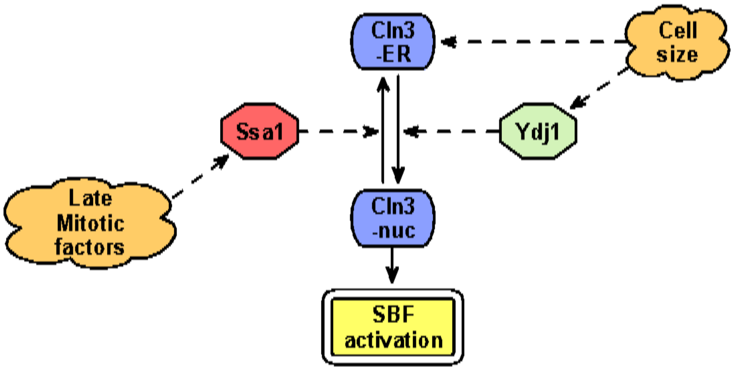


Figure S4. Cartoon of Cln3 regulation by Ydj1/Ssa1.

The nuclear form of Cln3 is the active form. Nuclear import is controlled by Ydj1 in response to cell size, whereas sequestration of Cln3 in the ER (inhibiting Cln3) is done by Ssa1, probably, in response to late mitotic factors. (See main text for details) We assume that Bck2 is regulated in a similar fashion.

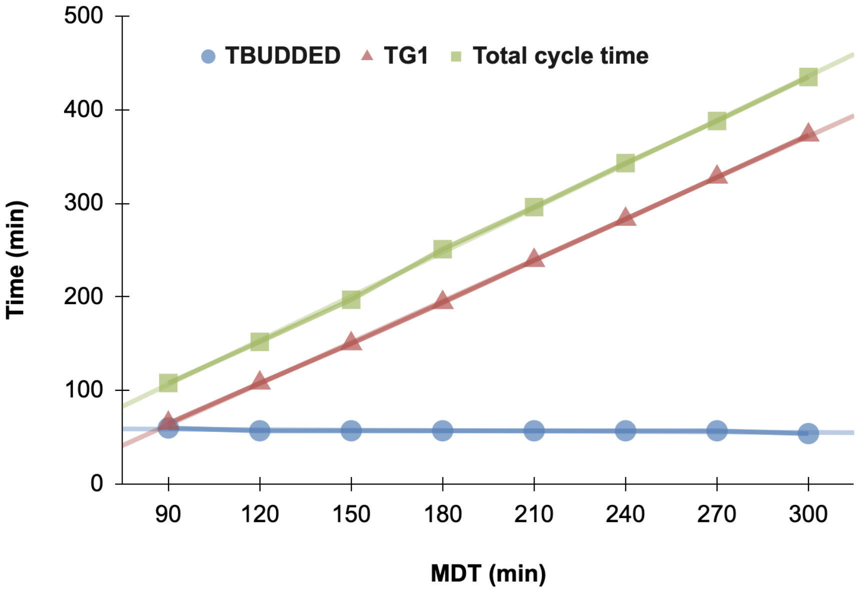


Figure S5. Duration of daughter cycle times as function of mass doubling time.

The duration plots of daughter cycle times (green filled squares) and G1 (red filled triangles) phase versus mass doubling time correspond to the experiments by Lord and Wheals (1980). The budded period (blue filled circles) was however, close to 50 and almost constant over the wide range, differing quantitatively from the corresponding curve in the experiments.

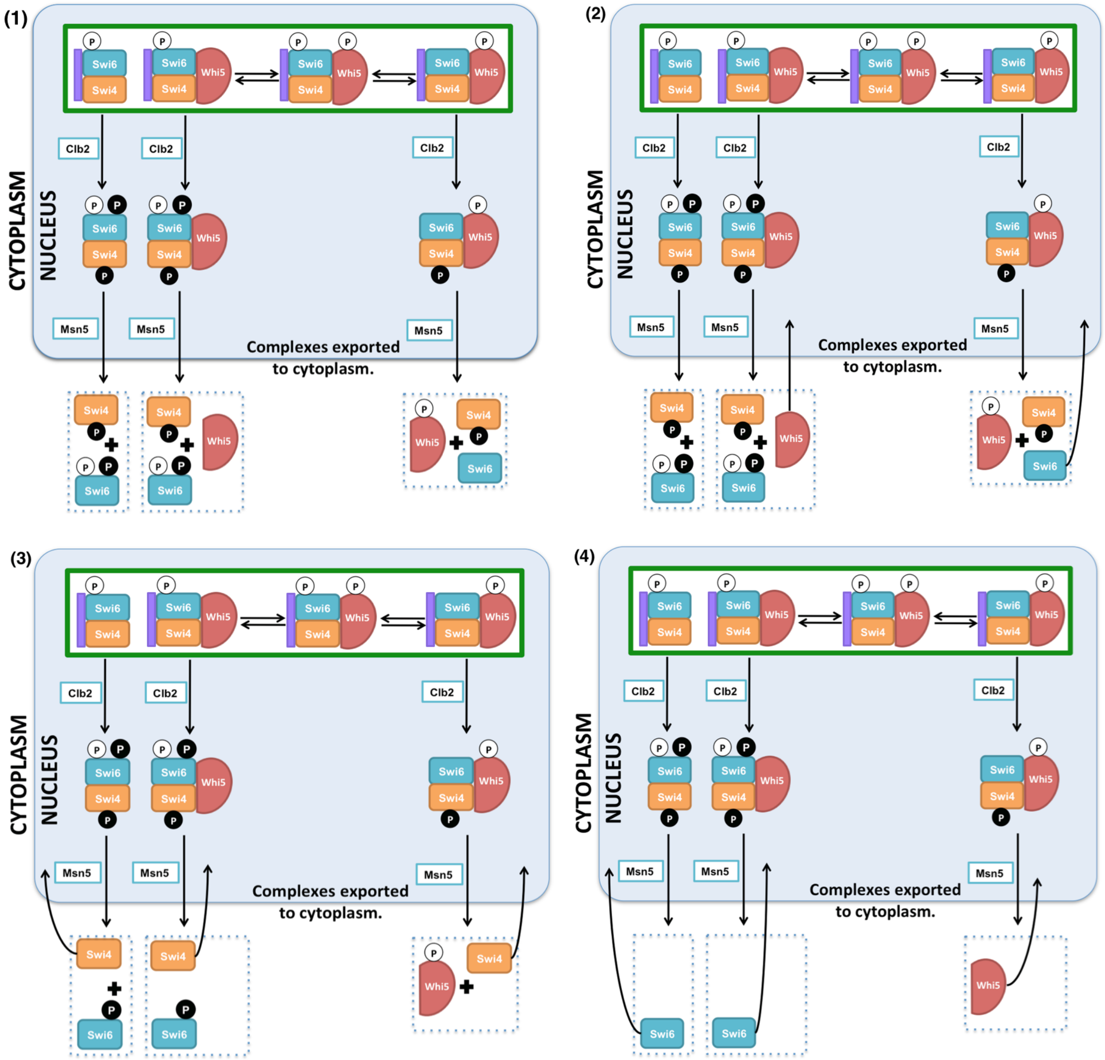


Figure S6. Localization of different monomers.

This figure describes the timing of re-import and hence overall temporal localization of the different monomers (Swi4, Swi6, Whi5). Export and reimport of Whi5P, although not shown explicitly, follows the same steps as phosphorylated Whi5 in the cartoon. Step (1): SBF complexes that have been phosphorylated on Whi5 or the S160 site of Swi6 are transported to the cytoplasm by Msn5 and dissociate immediately. Phosphorylated Whi5 monomers are also exported by Msn5. Step (2): Unphosphorylated monomers move back to the nucleus (regardless of the phase of the cell cycle). Step (3): Swi4 and the P-form of Swi6 (all phosphorylation sites except S160) get dephosphorylated by PP2A and move to the nucleus. Step (4): the phosphatase Cdc14 that accumulates at mitotic exit dephosphorylates Whi5 and Swi6 Q-form at residue S160, following which Whi5 and Swi6 get reimported to the nucleus resetting the localization state for the G1-phase of the next cell cycle.

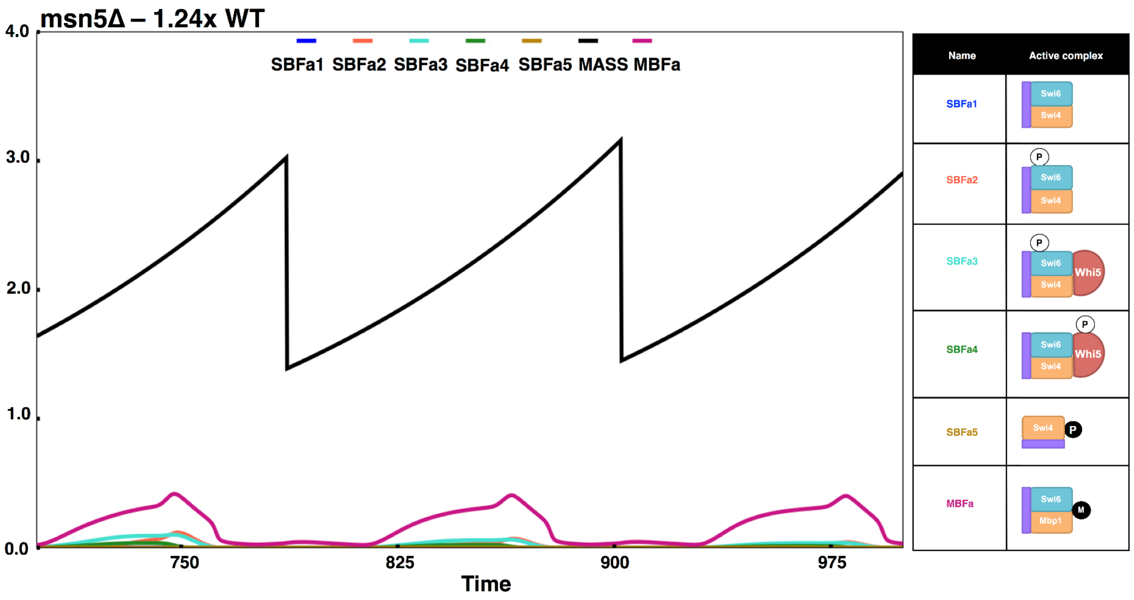


Figure S7. Importance of the transport protein, Msn5.

These simulations are plotted for msn5Δ cells (MSN5=0 in our model). The cells are slightly larger than WT cells due to the absence of active SBF, and lesser amount of active MBF (MBFa in plot above).

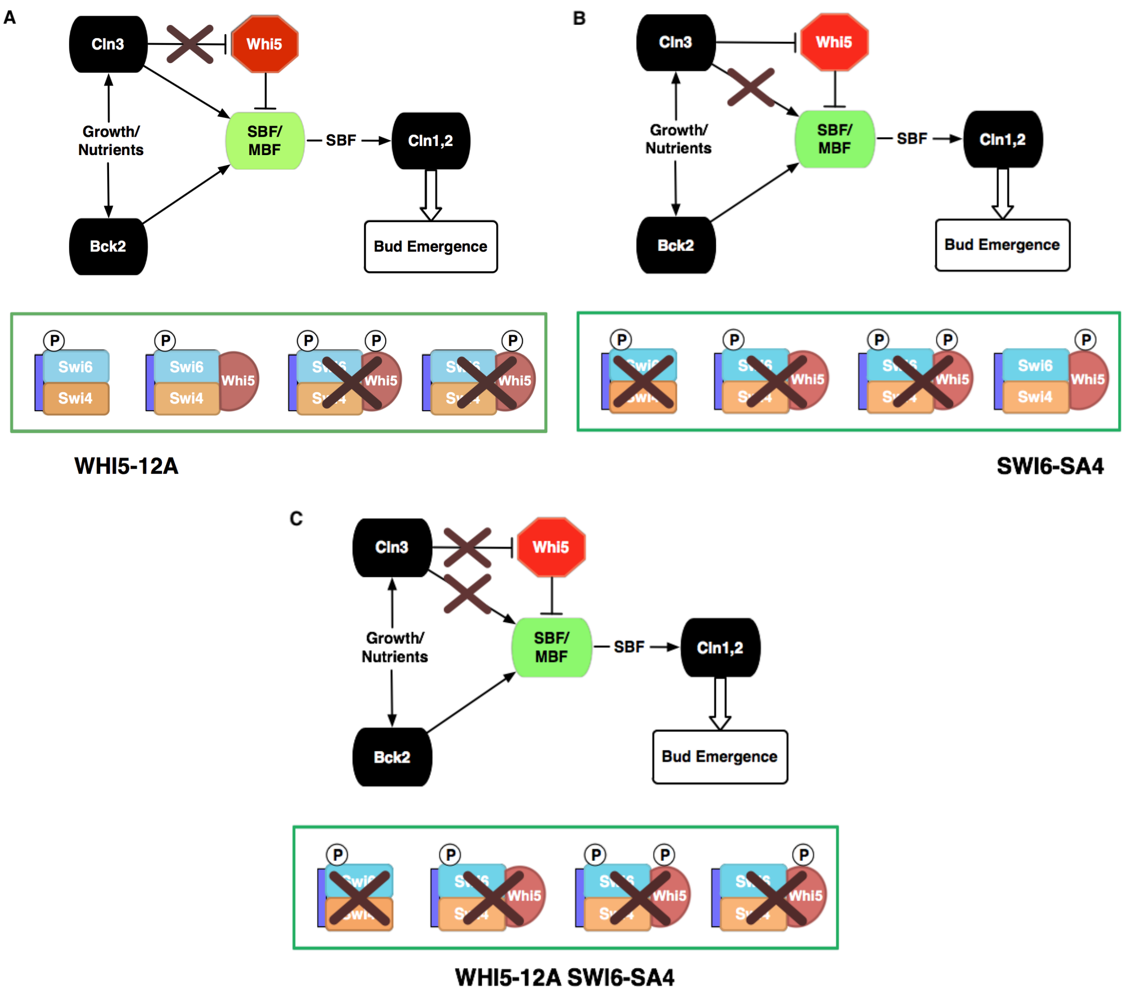


Figure S8. Non-phosphorylable mutants.

Active complexes in (A) *WHI5-12A* (This mutant will have size similar to that of WT due to the Swi6 P-forms; Figure 7A), (B) *SWI6-SA4* (WT size due to phosphorylation of Whi5; Figure 7B), (C) *WHI5-12A SWI6-SA4* (Viable, yet large, due to inactive SBF-Whi5 complex and support from Bck2 activation and MBF; Figure 7C).

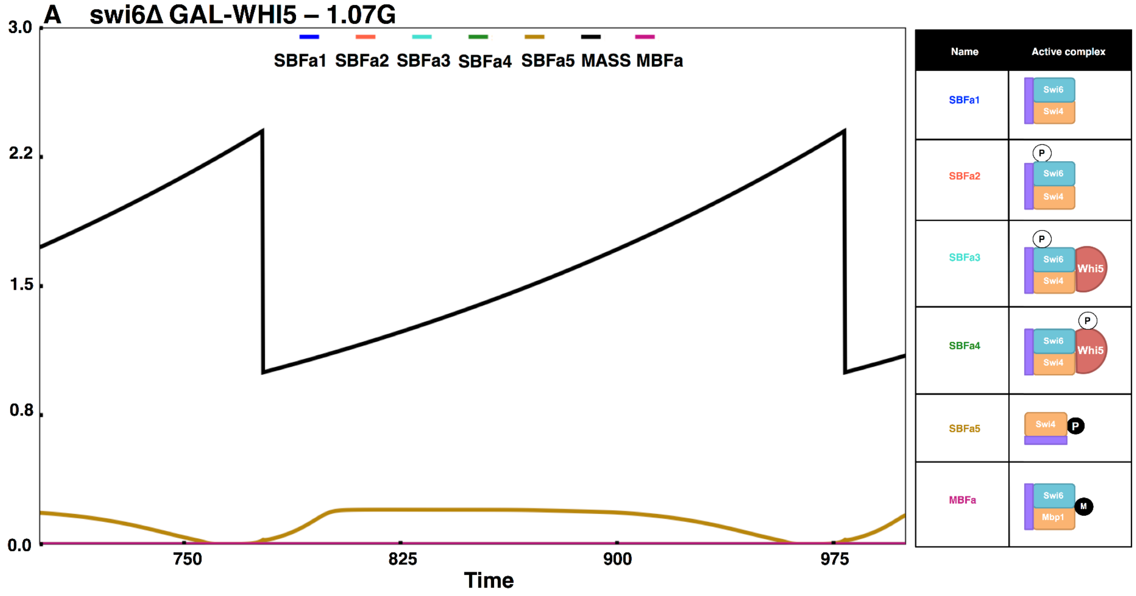
A swi6Δ GAL-WHI5 – 1.07G
SBFa1 SBFa2 SBFa3 SBFa4 SBFa5 MASS MBFa
3.0
2.2
1.5
0.8
0.0
750
825
900
975
Time
Name
Active complex
SBFa1
SBFa2
SBFa3
SBFa4
SBFa5
MBFa
Swi6
Swi4
Whi5
Mbp1
P
M

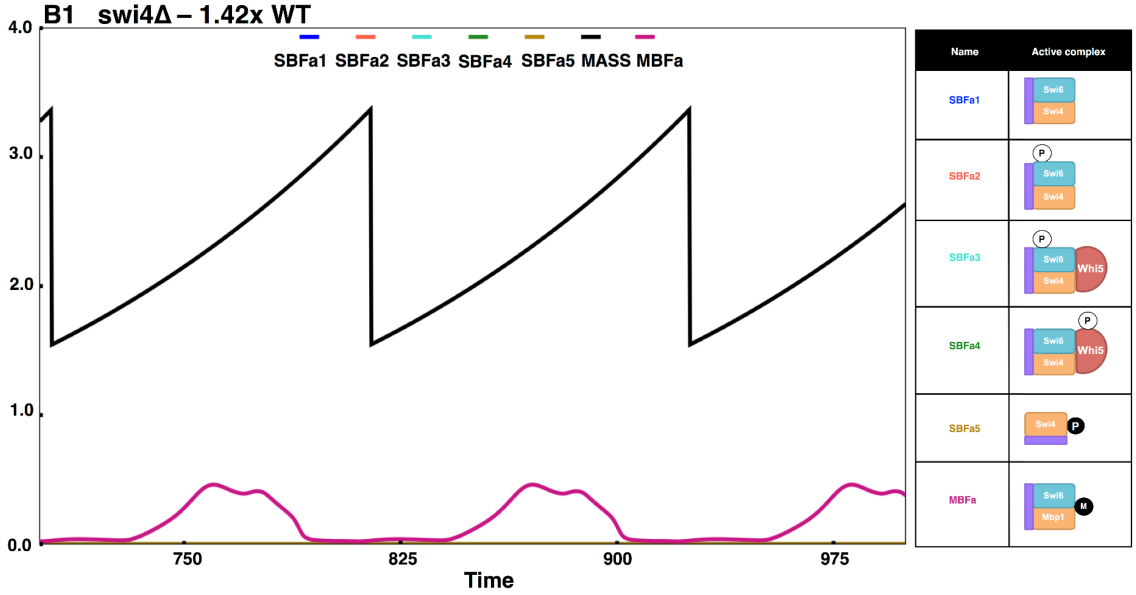
B1 swi4Δ – 1.42x WT
SBFa1 SBFa2 SBFa3 SBFa4 SBFa5 MASS MBFa
4.0
3.0
2.0
1.0
0.0
750
825
900
975
Time
Name
Active complex
SBFa1
SBFa2
SBFa3
SBFa4
SBFa5
MBFa
Swi6
Swi4
Whi5
Mbp1
P
M

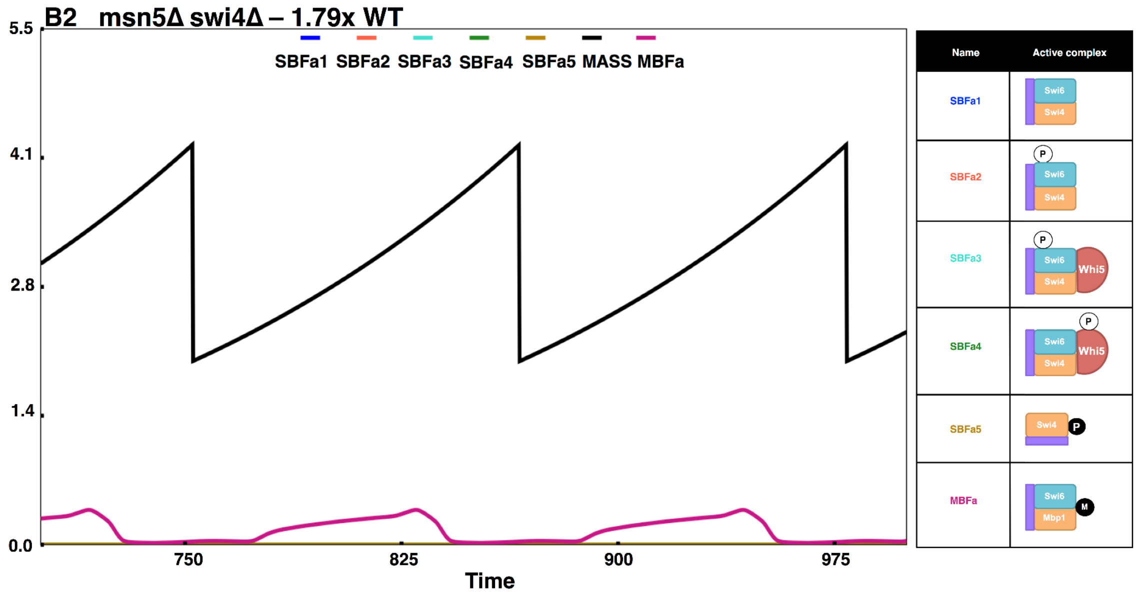
B2 msn5Δ swi4Δ – 1.79x WT
SBFa1 SBFa2 SBFa3 SBFa4 SBFa5 MASS MBFa
5.5
4.1
2.8
1.4
0.0
750
825
900
975
Time
Name
Active complex
SBFa1
SBFa2
SBFa3
SBFa4
SBFa5
MBFa
Swi6
Swi4
Whi5
Mbp1
P
M

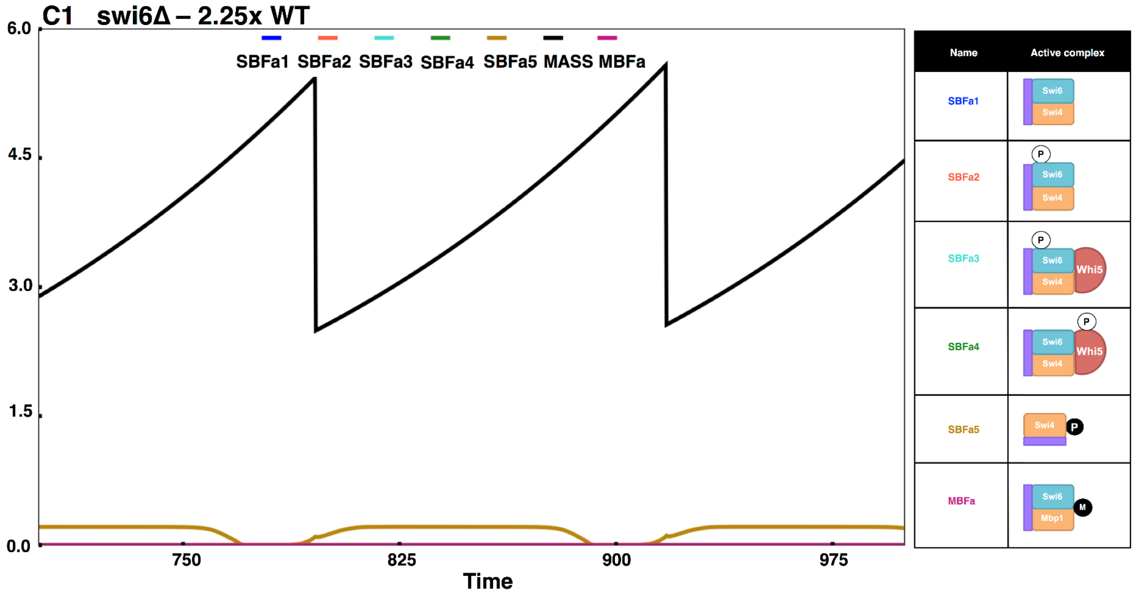
C1 swi6Δ – 2.25x WT
SBFa1 SBFa2 SBFa3 SBFa4 SBFa5 MASS MBFa
6.0
4.5
3.0
1.5
0.0
750
825
900
975
Time
Name
Active complex
SBFa1
SBFa2
SBFa3
SBFa4
SBFa5
MBFa
Swi6
Swi4
Whi5
Mbp1
P
M

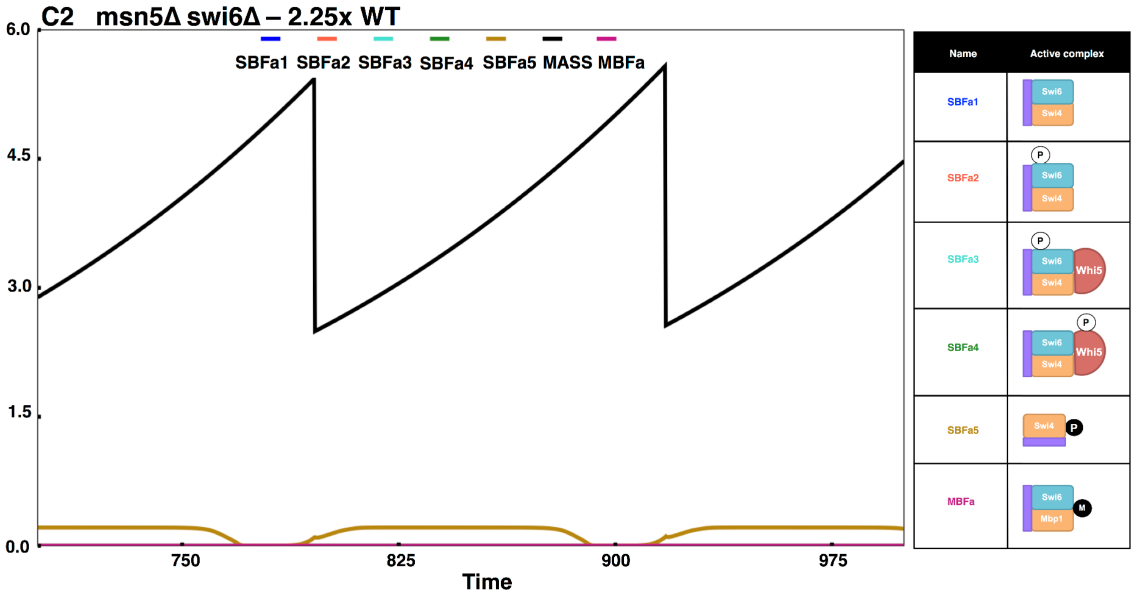
C2 msn5Δ swi6Δ – 2.25x WT
SBFa1 SBFa2 SBFa3 SBFa4 SBFa5 MASS MBFa
6.0
4.5
3.0
1.5
0.0
750
825
900
975
Time
Name
Active complex
SBFa1
SBFa2
SBFa3
SBFa4
SBFa5
MBFa
Swi6
Swi4
Whi5
Mbp1
P
M

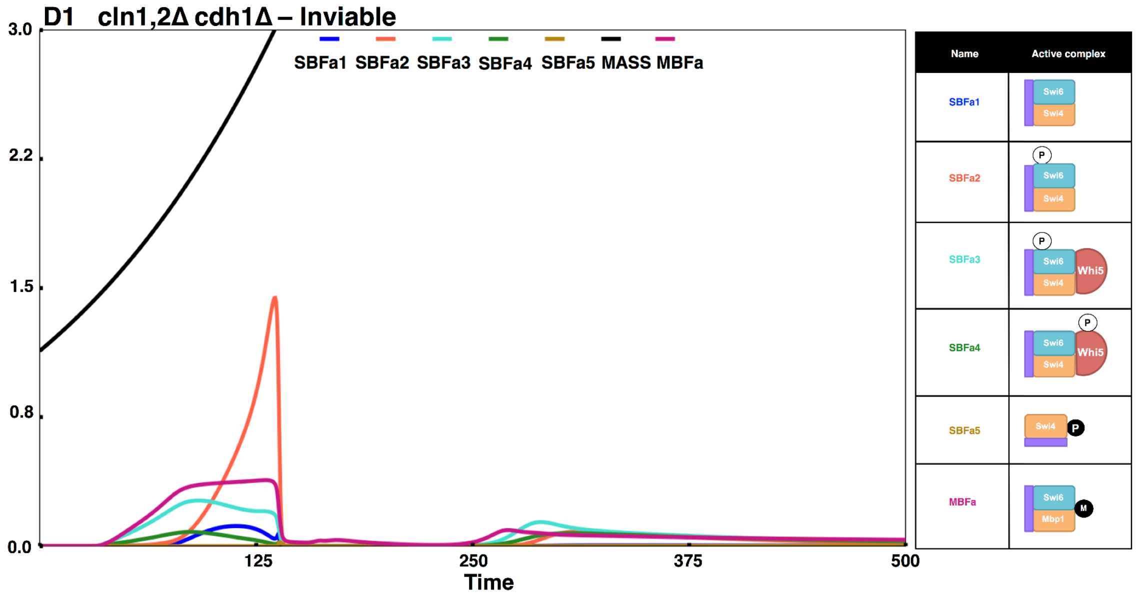
D1 cln1,2Δ cdh1Δ – Inviable
SBFa1 SBFa2 SBFa3 SBFa4 SBFa5 MASS MBFa
3.0
2.2
1.5
0.8
0.0
125
250
375
500
Time
Name
Active complex
SBFa1
SBFa2
SBFa3
SBFa4
SBFa5
MBFa
Swi6
Swi4
Whi5
Mbp1
P
M

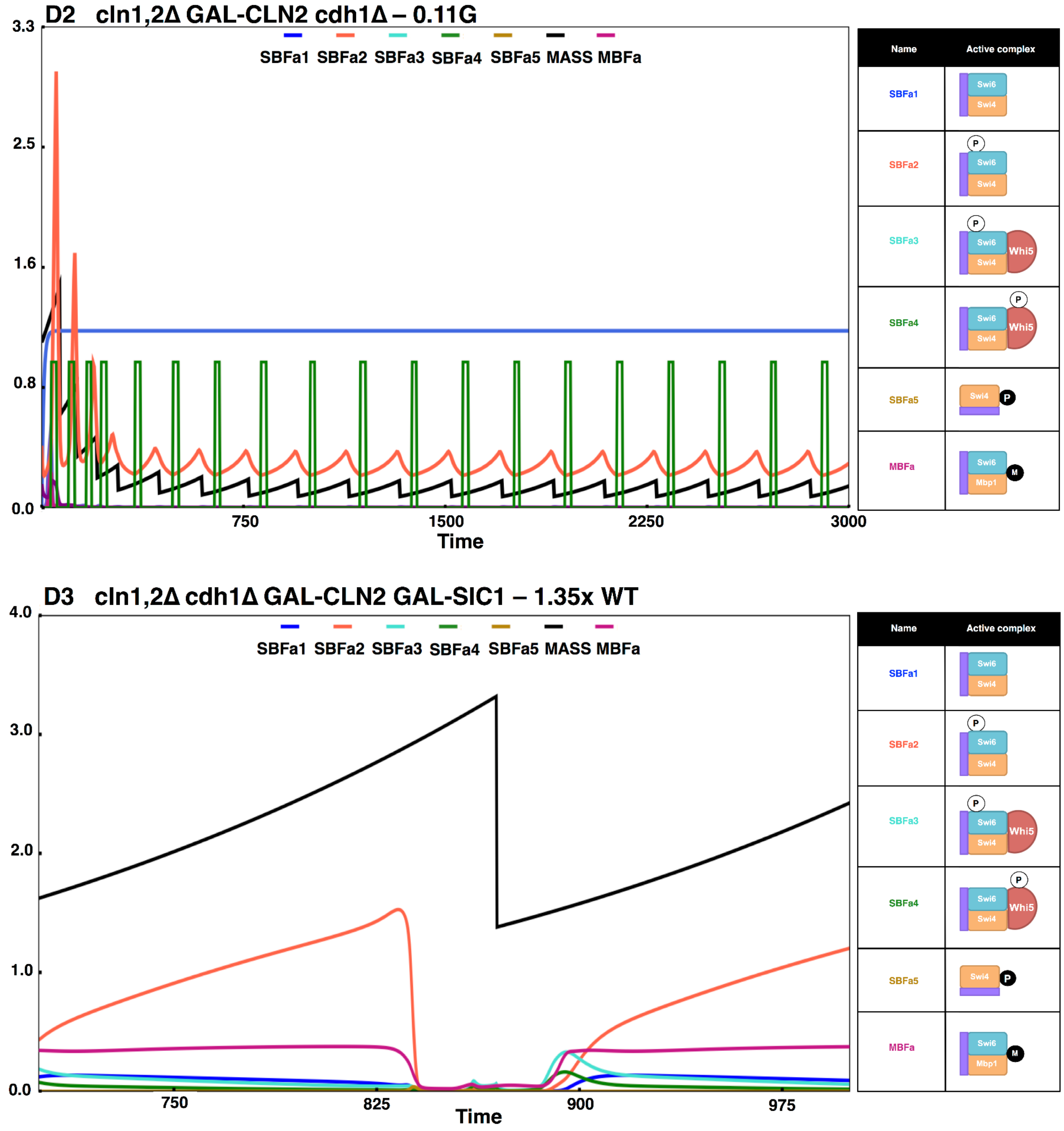


Figure S9. Model contradictions.

The figure shows simulations of the following mutants: (A) *swi6Δ GAL-WHI5* (only Swi4dimers present (SBFa5); cells are viable yet large), (B) B1: *swi4Δ* (only MBF present; cells are viable yet large), B2: *msn5Δ swi4Δ* (only MBF present; cells are viable and large), (C) C1: *swi6Δ* (only Swi4dimers (SBFa5) present; cells are viable yet large), C2: *msn5Δ swi6Δ* (only Swi4dimers (SBFa5) present; cells are viable yet large), (D) D1: *cln1Δ cln2Δ cdh1Δ* (cells are inviable), D2: *cln1Δ cln2Δ cdh1Δ GAL-CLN2* (cells are viable, very small). A, B2, C2, D1 and D2 are in contradiction with experimental findings, while D3: *cln1Δ cln2Δ cdh1Δ GAL-CLN2 GAL-SIC (cells are viable) is complementary with our hypothesis about the effect of* Cdh1 on Polo and Cdc14 on cell viability due to higher CKI levels*.*

A bck2Δ mbp1Δ – 1.43x WT

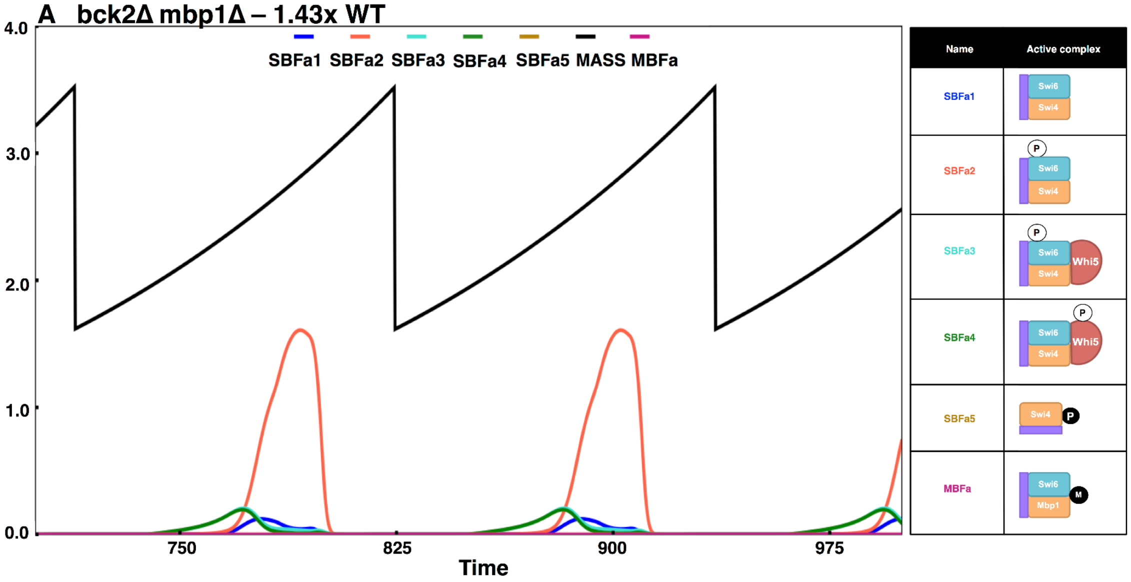


B bck2Δ mbp1Δ GAL-WHI5 – 1.55G

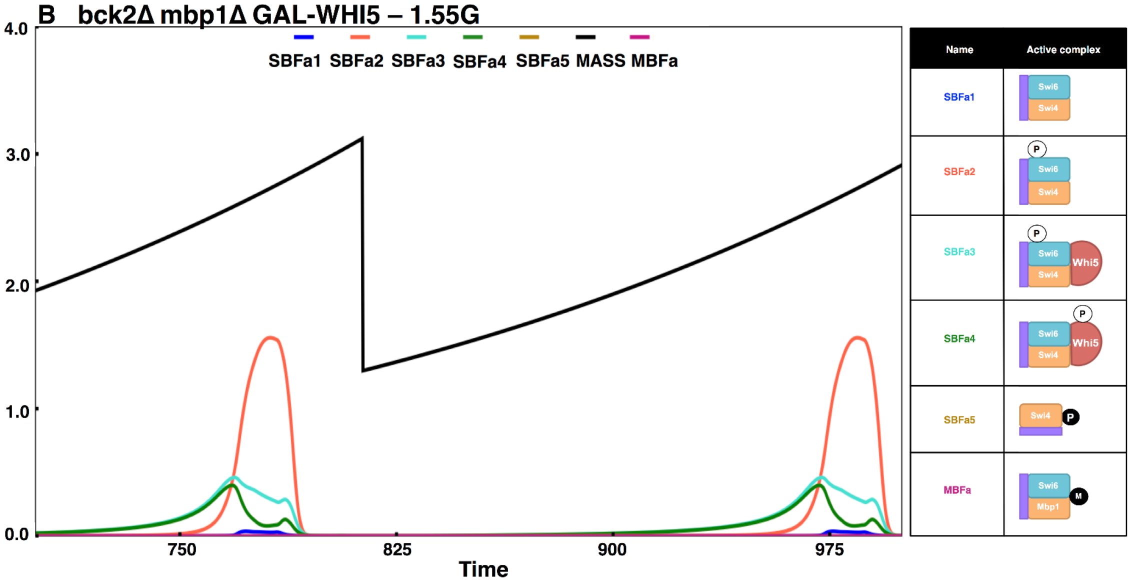

C cln3Δ mbp1Δ – 2.55x WT

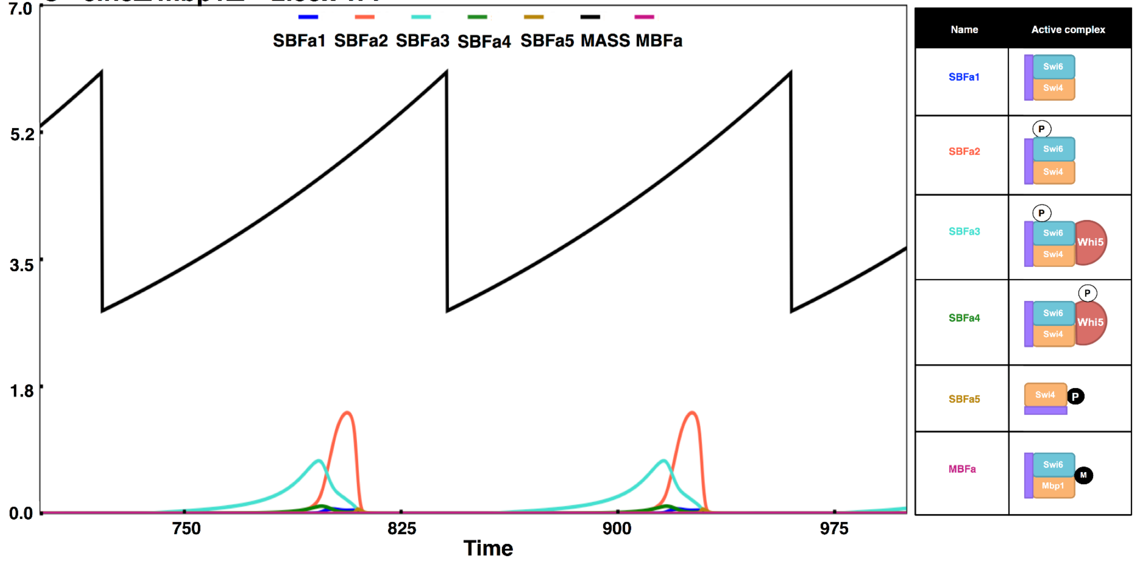


D cln3Δ mbp1Δ swi6Δ – 2.35x WT

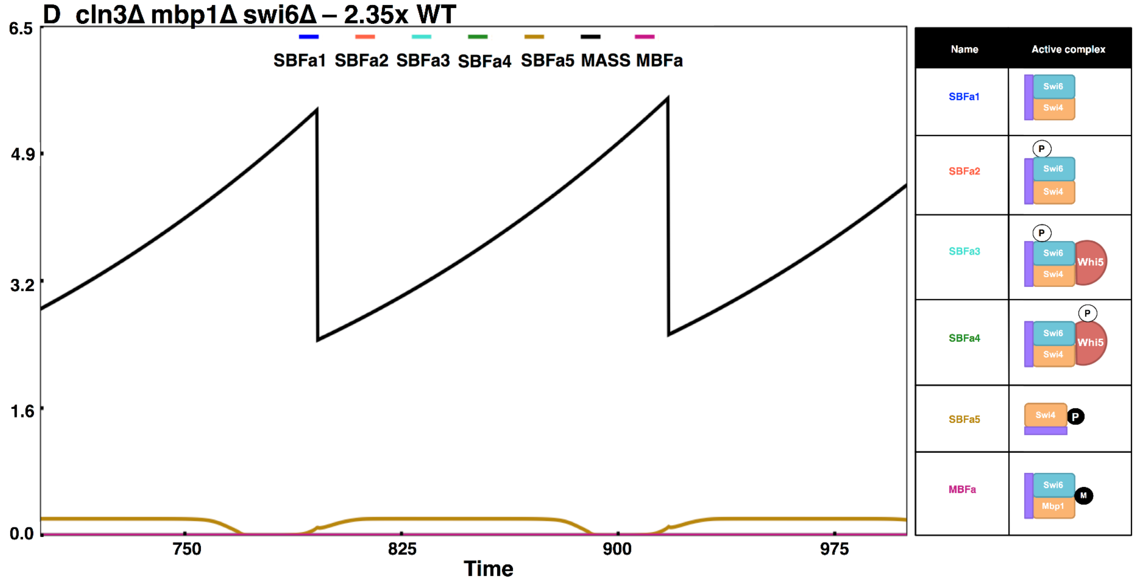

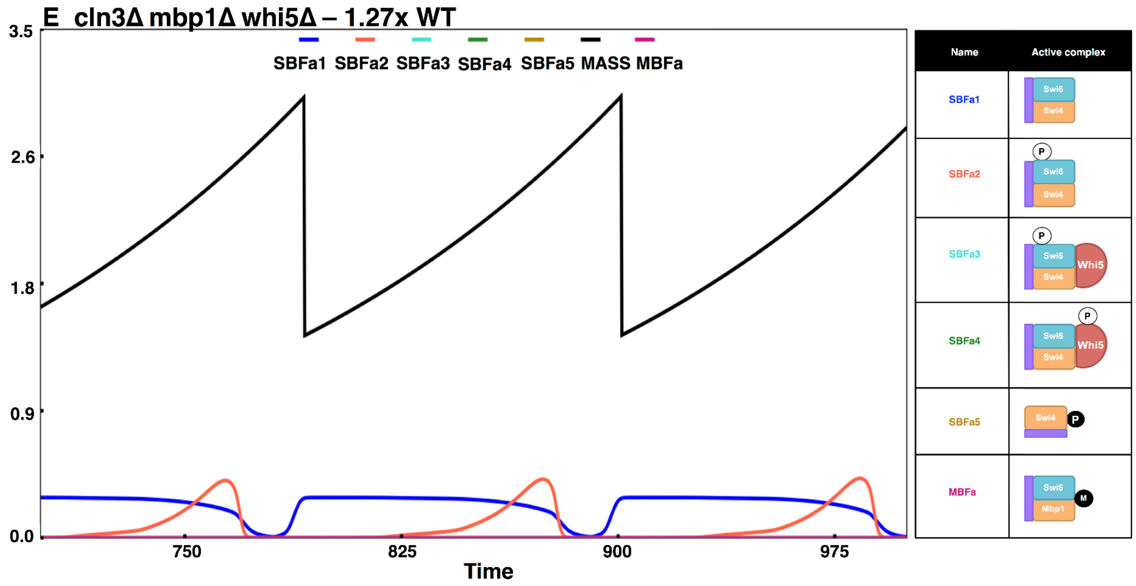
E cln3Δ mbp1Δ whi5Δ – 1.27x WT
SBFa1 SBFa2 SBFa3 SBFa4 SBFa5 MASS MBFa
3.5
2.6
1.8
0.9
0.0
750
825
900
975
Time
Name
Active complex
SBFa1
SBFa2
SBFa3
SBFa4
SBFa5
MBFa
Swi6
Swi4
Whi5
Mbp1
P
M

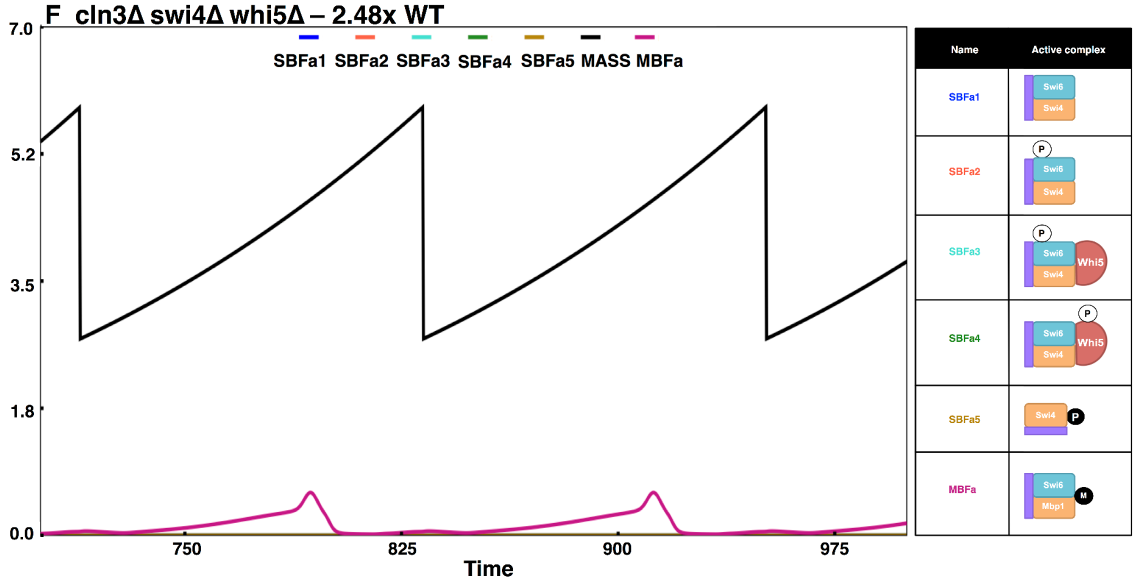
F cln3Δ swi4Δ whi5Δ – 2.48x WT
SBFa1 SBFa2 SBFa3 SBFa4 SBFa5 MASS MBFa
7.0
5.2
3.5
1.8
0.0
750
825
900
975
Time
Name
Active complex
SBFa1
SBFa2
SBFa3
SBFa4
SBFa5
MBFa
Swi6
Swi4
Whi5
Mbp1
P
M

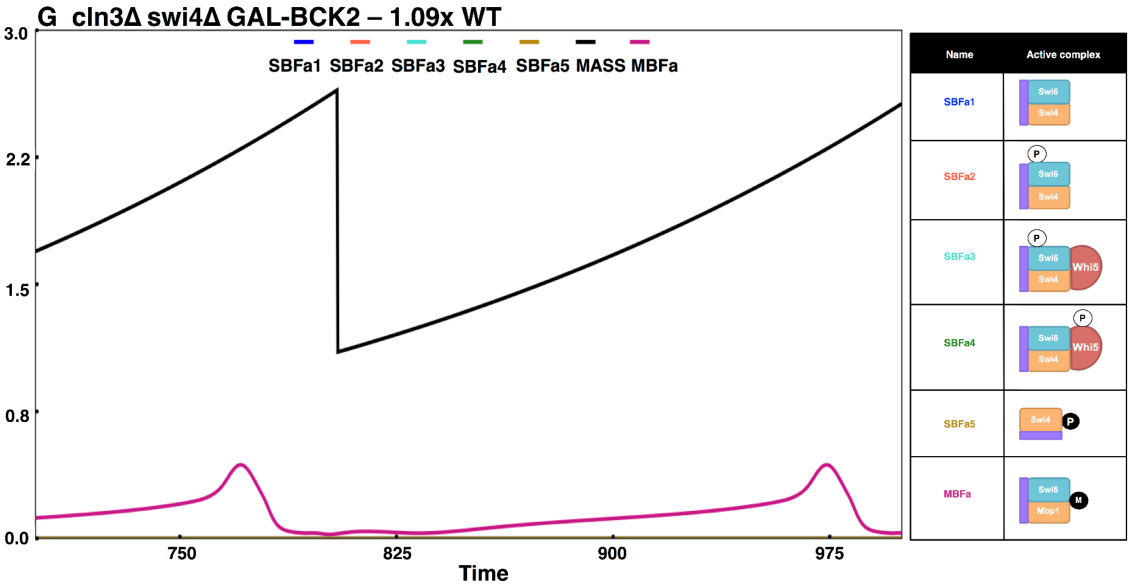
G cln3Δ swi4Δ GAL-BCK2 – 1.09x WT
SBFa1 SBFa2 SBFa3 SBFa4 SBFa5 MASS MBFa
3.0
2.2
1.5
0.8
0.0
750
825
900
975
Time
Name
Active complex
SBFa1
SBFa2
SBFa3
SBFa4
SBFa5
MBFa
Swi6
Swi4
Whi5
Mbp1
P
M

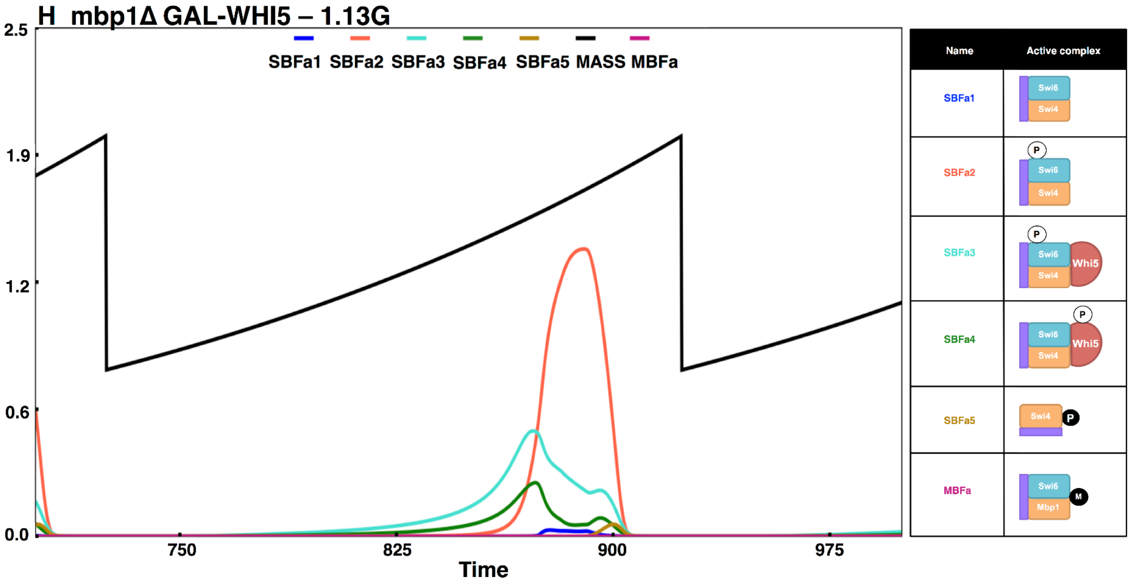
H mbp1Δ GAL-WHI5 – 1.13G
SBFa1 SBFa2 SBFa3 SBFa4 SBFa5 MASS MBFa
2.5
1.9
1.2
0.6
0.0
750
825
900
975
Time
Name
Active complex
SBFa1
SBFa2
SBFa3
SBFa4
SBFa5
MBFa
Swi6
Swi4
Whi5
Mbp1
P
M

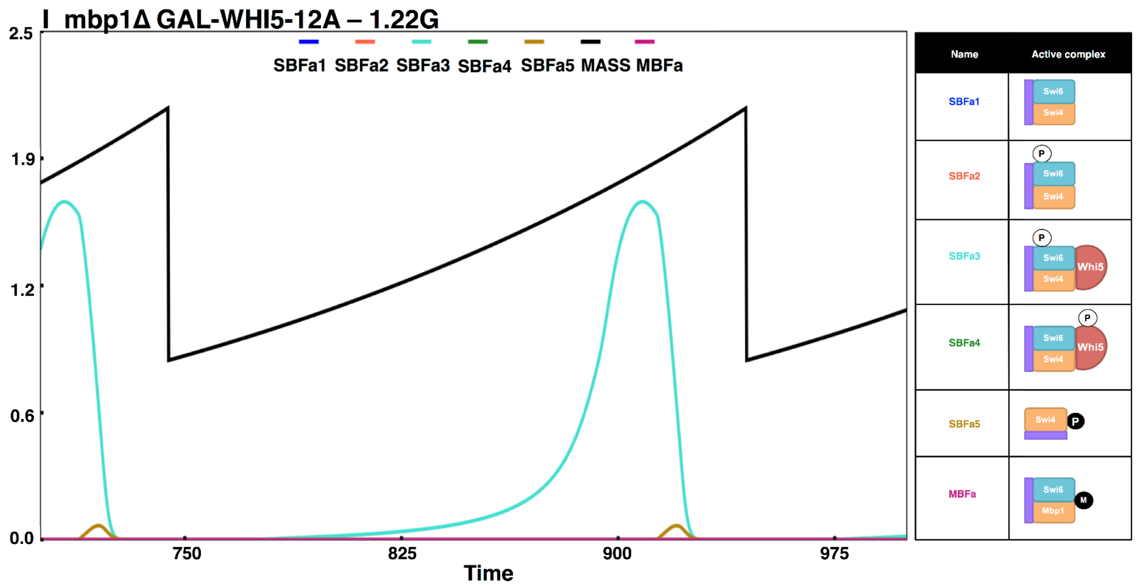


Figure S10. Few model predictions and validations.

(A) *bck2Δ mbp1Δ* (Cln-activated forms of SBF are present in varying fractions (SBFa2>SBFa3, SBFa4> SBFa1); cells are viable), (B) *bck2Δ mbp1Δ GAL-WHI5* (Cln-activated forms of SBF are present (SBFa2, SBFa3, SBFa4) even if they are lesser than in A; cells are still viable), (C) *cln3Δ mbp1Δ* (SBFa2 > SBFa3 >> SBFa4 > SBFa5 ;no MBF is present; cells are viable), (D) *cln3Δ mbp1Δ swi6Δ* (Swi4dimers (SBFa5) are present sufficiently enough to rescue cells; cells are viable), (E) *cln3Δ mbp1Δ whi5Δ* (Inhibition on SBF is relieved) ; cells are rescued), (F) *cln3Δ swi4Δ whi5Δ* (More Bck2-activated MBF is present (MBFa), there's no SBF; cells are viable), (G) *cln3Δ swi4Δ GAL-BCK2* (There's sufficient amount of Bck2-activated MBF (MBFa); cells are rescued), (H) *mbp1Δ GAL-WHI5* cells are viable since Whi5 is predicted to get activated and is present in the forms of SBFa2, SBFa3, and SBFa4 (I) *mbp1Δ GAL-WHI5-12A* cells are viable because Whi5 get converted to the activated form SBFa3.

### Supplementary Tables

Table S1. Functions of different Cyclins in Budding Yeast cell cycle

| | |
|---|---|
| Cln1,2 | - Bud emergence, START<br>- Inhibiting kinase inhibitors Sic1, Cdc6 and Cdh1 (which inhibit Clbs)<br>- In the model, Cln2 represents Cln1,2 |
| Cln3 | - Activates SBF (TF for Cln1,2), MBF (TF for Clb5,6) in response to cell size |
| Clb1,2 | - Late mitotic events including elongation of short mitotic spindle<br>- Promotes cell cycle transition into mitosis<br>- In the model, Clb2 represents Clb1,2 |
| Clb3,4 | - Early mitotic events including formation of short mitotic spindle<br>- Involved in DNA synthesis, spindle assembly as well as G2/M transition |
| Clb5,6 | - DNA Synthesis<br>- Inhibiting kinase inhibitors Sic1, Cdc6 and Cdh1<br>- In the model, Clb5 represents Clb5,6 |

Table S2. Description, abundance, regulation and localization of START components in the model; List of Key Assumptions

| Protein / Component | Description/Regulation |
| --- | --- |
| Whi5 | ***Description***<br>- Whi5 is a stoichiometric inhibitor of the transcription factor complexes, SBF[1,2] and MBF[1].<br>***Abundance***<br>- The abundance of Whi5 is roughly 1500 protein molecules per cell in an asynchronous culture[3]), although Whi5 is known to be transcriptionally regulated showing a 3-fold variation, peaking in late G1 phase[4]. Therefore, we assume available Whi5 (for inhibition) in G1 to be 1000 molecules.<br>- For simplicity, we assume total Whi5 to be conserved in the model.<br>***Regulation & Localization***<br>- Whi5 has 12 known CDK phosphorylation sites and 6 other non-CDK sites[5]. We assume Hill kinetics (with Hill's coefficient=5) for CDK phosphorylation of Whi5 to capture the non-linearity resulting from multi-site phosphorylation (compared and tested for a smaller version of model).<br>- We assume that Whi5 is phosphorylated by Cln3, Cln2 and Clb5 (with different efficiencies) and dephosphorylate by Cdc14 at mitotic exit[1,2,6]. We also assume that all forms of Whi5 (either free, or SBF-bound or SBF-promoter-bound) are subjected to phosphorylation and dephosphorylation.<br>- Whi5 is nuclear until late G1, and moves to cytoplasm prior to START, and stays cytoplasmic until mitotic exit[7]. This localization is dependent on phosphorylation of specific CDK phosphorylation sites on Whi5, and the transport protein Msn5. |
| Swi6 | ***Description***<br>- Swi6 is a **component** of the transcription factor, **SBF** and **MBF**[8,9].<br>***Abundance***<br>- It is 3 times more abundant than Whi5[3].<br>- We assume total Swi6 to be conserved in the model.<br>***Regulation & Localization***<br>- Swi6 has 5 consensus CDK phosphorylation sites[10].<br>- We assume that the Cln kinase phosphorylation on Whi5 and its phosphorylation on Swi6 have the same dynamics. That is, we |

| | |
|---|---|
| | use the same Hill function to describe the phosphorylation of Cln kinase on Whi5 as well as on Swi6.<br>- In the model, the sites bearing the P-form – the activatory phosphorylations are done by Cln3, Cln2 and Clb5 with different efficiencies, and the dephosphorylation is by unspecified phosphatase.<br>- Phosphorylation at the S160 site, is known to be responsible for the cytoplasmic localization of Swi6 from mid-S phase until mitotic exit[11]. This is considered to be the Q-form (inactivated form) in the model and is required for cytoplasmic localization. The Q-form phosphorylation is carried out by Clbs (Clb5 and Clb2 in our model), and dephosphorylated by Cdc14 at mitotic exit[12]. Transport of Q-forms to the cytoplasm requires Msn5 and Swi4[13].<br>- We assume phosphorylation (P- and Q-forms) occurs for all forms of Swi6 (either free or promoter bound forms). |
| Swi4 | ***Description***<br>- Swi4 is a component of the transcription factor, SBF[8,9].<br>***Abundance***<br>- Based on observed protein abundances, we consider the relative abundance of Swi4 and Mbp1 to be ~0.55x as abundant as Whi5[3]. Therefore, Swi4 is the limiting component of the SBF complex.<br>- We assume total Swi4 to be conserved in the model.<br>***Regulation and Localization***<br>- We assume that phosphorylation of Swi4 is necessary for SBF inactivation[14]. Phosphorylation of Swi4 subunit of SBF causes SBF to dissociate from the promoter.<br>- Swi4 is nuclear at all times[15]. In the model, phosphorylated Swi4 gets dephosphorylated by a constitutively active phosphatase, PP2A. The unphosphorylated form remains nuclear at all times.<br>- Since *swi6Δ* mutant is viable, but *swi6Δ bck2Δ* is inviable, we assume that Swi4 has some residual activity in the absence of SBF[8,16], but it requires Bck2 for its activity. The active Swi4 complex is also assumed to be inactivated by Clb2. |
| Mbp1 | ***Description***<br>- Mbp1 is a component of the transcription factor, MBF[8,9].<br>***Abundance***<br>- As mentioned above, we assume Mbp1 to have a relative abundance of 0.55x w.r.t Whi5. Mbp1 is, therefore, a limiting component of MBF[3]. |

| | |
|---|---|
| | - We assume total Mbp1 to be conserved in the model.<br>***Regulation and Localization***<br>- MBF regulation is driven by Cln3 and Bck2 for activation[8], and Nrm1[17] and Clb2 (to a much lesser extent) for inactivation.<br>- Since little information is available about the modification or localization of Mbp1, they are not considered in the model. |
| Msn5 | ***Description***<br>- Msn5 is a transport protein that exports phosphoproteins (Swi6 and Whi5 in our model) from the nucleus to the cytoplasm[6,13].<br>***Regulation***<br>- We do not consider the regulation of Msn5. It is assumed to be constant in our model. |
| Cln3 & Bck2 (Modification from Chen2004) | ***Regulation & Localization***<br>- **Activation** of Cln3, which is by nuclear import, depends on chaperone protein, Ydj1, which is proposed to be the sensor of cell size[18]. In the model, Ydj1 depends on mass. We also assume that the abundance of Cln3 depends on mass. This assumption is necessary for mutant cells with longer G1 to have shorter G2/M so that the total cycle time (G1 + S/G2/M) is the same as wild type cells. Thus, mutant cells can maintain their size homeostasis generation after generation.<br>- **Inactivation** of Cln3 is done by Ssa1[18]. To explain the change of Cln3 localization during the cell cycle, (being nuclear in late G1, just prior to START and become cytoplasmic from late S phase on), we assume that Ssa1 is activated by Clb2 and Swi5, both of which accumulate during M-phase.<br>- Currently, there is no evidence for Bck2 being regulated the same way as Cln3. But since Bck2 is known to be a cell size regulator too[19], we assume that its activation and inactivation depend on similar mechanisms that are contingent on mass and Clb2, Swi5 (instead of Ydj1 and Ssa1). |
| Promoters | - We assume the promoter concentration to be 0.2 relative to Whi5 abundance, corresponding to 200 of SBF regulated genes.<br>- We consider two species, Prom2 and Prom5 corresponding to genes regulated by SBF and MBF, respectively. |

| List of major assumptions in the model |
| --- |
| SBF and MBF overlap functionally. |
| START proteins Swi4, Swi6, Mbp1 and Whi5 are expressed constitutively. |
| Whi5 inhibits SBF strongly and MBF weakly. |
| Relative abundances of Swi4, Mbp1, Swi6, Whi5, SCB, MCB are 5.5, 5.5, 30, 10, 2, 2. |
| Cln3-kinase activates SBF in two ways: by activating Swi6 and by inactivating Whi5. |
| Phosphorylations of SBF by G1/S cyclins are described by a Hill function with nH = 5. |
| Clb1,2-kinases inactivate SBF by phosphorylating Swi4 and Swi6. |
| Whi5 is exported from the nucleus by Msn5 when it is phosphorylated. |
| Phosphorylated Whi5 dissociates from the promoter if the Swi6 moiety in the Swi4/Swi6/Whi5 complex is also phosphorylated. |
| Swi4/Swi6 complex is exported from the nucleus by Msn5 when Swi6 is phosphorylated. |
| Swi4 is dephosphorylated constitutively by an active phosphatase and, hence, it is nuclear throughout the cell cycle. |
| Bck2 and Cln3 activate SBF and MBF in a similar manner. |
| A Swi4-only form of SBF (that requires Swi4 and Bck2 but not Swi6) is responsible for the viability of swi6Δ cells. |
| MBF, like SBF, is activated by Cln3 and Bck2; it is inactivated by Clb2 and Nrm1. |
| Nuclear entry of Cln3 and Bck2 are controlled by Ydj1 and Ssa1 in a cell-size dependent manner. |
| Ssa1 is responsible for nutrient modulation of the critical size threshold for START. |
| Rates of Clb1,2 and Clb5,6 synthesis increase with cell mass. |
| The various promoter-bound forms SBF and MBF have different activities. |
| A simulated cell is considered viable if and only if it satisfies certain rules for “viability”. |

Table S3.Modifications in Parameter & Initial conditions corresponding to mutants. (Mutants exclusive to the current model are emphasized in bold).

| Mutants | Parameters changed in model |
|---|---|
| **Loss of function mutants** | |
| **G1, S** | |
| *cln3Δ* | CLN3T=0 |
| *bck2Δ* | BCK2T=0 |
| ***whi5Δ*** | init WHI5=0 |
| ***WHI5-12A*** | ef5p=0, mdt=150, init WHI5 = whi5op*WHI5 |
| ***swi4Δ*** | init SWI4=0 |
| ***swi6Δ*** | init SWI6=0 |
| ***SWI6-SA4*** | ef6p=ef6q=0 |
| ***mbp1Δ*** | init MBP1=0 |
| ***msn5Δ*** | MSN5=0 |
| *cln2Δ* | ksn2'=ksn2"=ksn2'"=0; init CLN2=0 |
| *clb5Δ* | ksb5'=ksb5"=ksb5'"=0; init CLB5=0 |
| *CLB5-dbΔ* | kdb5"=0 |
| **Cyclin Antagonists** | |
| *sic1Δ* | ksc1'=ksc1"=0; init SIC1=SIC1P=C2=C2P=C5=C5P=0 |
| *cdc6Δ (cdc6 2-49Δ)* | ksc6'=ksc6"=ksc6'"=0; init CDC61=CDC6P=F2=F2P=F5=F5P=0 |
| *ckiΔ* | ksc1'=ksc1"=ksc6'=ksc6"=ksc6'"=0;<br>init SIC1=SIC1P=C2=C2P=C5=C5P=CDC61=CDC6P=F2=F2P=F5=F5P |
| *cdh1Δ* | kscdh=0; CDH1=CDH1i=0 |
| **M-phase** | |
| *swi5Δ* | ksc1"=ksf6"=0 |
| *clb2Δ* | ksb2'=ksb2"=0 |
| *CLB2-dbΔ* | kdb2"=0.25*kdb2", kdb2'"=0 |
| *CLB1 clb2Δ* | ksb2'=0.33*ksb2', ksb2"=0.33*ksb2" |
| *cdc20Δ* | ks20'=ks20"=0 |
| *cdc20-ts* | |
| *apc-ts* | ks20'=ks20"=kscdh=0; init CDH1=CDH1i=0 |
| *APC-A* | ka20"=0 |
| *pds1Δ* | kspds'=0; PDS1=PE=ESP1=0 |
| *PDS1-dbΔ* | kdpds"=kdpds'"=0 |
| *esp1-ts* | kasesp=0.002*kasesp, kdirent=0.002*kdirent, ki=0.04*ki |
| *ppxΔ* | PP2AT=0 |
| *tem1-ts* | ka15"=0.003*ka15 |
| *net1-ts* | kasrent=0.04*kasrent, kasrentp=0.04*kasrentp |
| *cdc15Δ* | kpnet"=0 |
| *TAB6-1* | kasrent=0.02*kasrent, kasrentp=0.02*kasrentp |
| *cdc14-ts* | ks14=0; init CDC14=0 |
| *mad2Δ* | mad2h=0.01; init MAD2=0.01 |
| *bub2Δ* | bub2h=bub2l=0; init BUB2=0 |
| *Cells in nocodazole* | ksspn=0 |

| Over-expression (gain of function) mutants | |
|---|---|
| *Cells in galactose* | mdt=150 (applicable for all *GAL* mutants below) |
| *GAL-CLN3* | CLN3T=kgalcln3*CLN3T |
| ***GAL-BCK2*** | BCK2=3*BCK2T |
| ***GAL-WHI5*** | init WHI5=10*WHI5 |
| ***GAL-WHI5-12A*** | ef5p=0; init WHI5=10*WHI5 |
| *GAL-CLN2* | ksn2'=0.165 |
| *GAL-CLB5* | ksb5'=0.016 |
| *GAL-SIC1* | ksc1'=0.132 |
| *GAL-CDC6* | ksf6'=0.4 |
| *GAL-CLB2* | ksb2'=0.38 |
| *GAL-CDC20* | ks20'=6 |
| *GALL-CDC20* | ks20'=0.6 |
| *GAL-ESP1* | init ESP1=4*ESP1, PE=4*PE |
| *GAL-PDS1* | kspds'=0.2 |
| *GAL-PPX* | PP2AT=6 |
| *GAL-CDC15* | init CDC15i=20*CDC15i, CDC15=20*CDC15 (20 copies) |
| *GAL-NET1* | ksnet=4*ksnet |
| *GAL-CDC14* | ks14=4*ks14 |
| *GAL-TEM1* | init TEM1GDP=20*TEM1GDP, TEM1GTB=20*TEM1GTB (20 copies) |
| *CLN3-1* | CLN3T=kmccln3*CLN3T |
| *CDH1 constitutively active* | mdt=150, kicdh"=0, kscdh=3*kscdh |
| *GAL-SIC-dbΔ* | mdt=150, ksc1'=kgalsic1 , kd3c1=0.132 (same as GAL-SIC1) |
| *GAL-CLB5-dbΔ* | mdt=150, ksb5'=kgalblb5 , kdb5"=0.016 (same as GAL-CLB5) |
| *GAL-PDS1-dbΔ* | kdpds"=kdps'''=0 |
| **Multi-copy (mc) mutants** | |
| mc BCK2 | BCK2T=5*BCK2T |
| mc CLN2 | ksn2"=4*ksn2", ksn2'''=4*ksn2''' |
| mc CLB5 | ksb5'=4*ksb5', ksb5"=4*ksb5", ksb5'''=4*ksb5''' |
| mc GAL-CLB2 | mdt=150, ksab2'=0.72 |
| mc SIC1 | ksc1'=4*ksc1', ksc1"=4*ksc1" |
| mc CDC6 | ksf6'=0.4*ksf6', ksf6"=0.4*ksf6", ksf6'''=0.4*ksf6''' |
| mc CDC20 | ks20'=5*ks20', ks20"=5*ks20" |
| mc CDC14 | ks14=3*ks14 |
| mc TEM1 | init TEM1GDP=20*TEM1GDP, TEM1GTB=20*TEM1GTB (20 copies) |
| mc CDC15 | init CDC15i=20*CDC15i, CDC15=20*CDC15 (20 copies) |

Table S4. START mutants

In blue: Predictions; In red: Contradictions; *GAL* mutants are scaled w.r.t *GAL*-WT.
Size of WT in Glucose: 1x; Size of WT in Galactose: 1G

| # | Genotype | Experimental phenotypes | Simulation results | References |
|---|---|---|---|---|
| ***Mutants pertaining to SBF, MBF; G1, G1/S, S cyclins; Bck2 and cyclin antagonists.*** | | | | |
| 1 | *WHI5-12A* | 1x | 1.07x | Wagner 09 |
| 2 | *SWI6-SA4* | 1x | 1.09x | Wagner 09 |
| 3 | *SWI6-SA4 WHI5-12A* | 1.4x | 1.73x | Wagner 09 |
| 4 | *GAL-WHI5-12A* | >1x | 1.2G | Wagner 09 |
| 5 | *GAL-WHI5-12A SWI6-SA4* | Inviable | G1 arrest | Wagner 09 |
| 6 | *bck2Δ* | 1.3x | 1.32x | Wijnen 99 |
| 7 | *Multi-copy BCK2* | 0.8x | 0.77x | Di Como 95 |
| 8 | *GAL-BCK2* | Viable | 0.75G | Costanzo 04 |
| 9 | *cln2Δ* | 3.2x | 1.92x | Dirick 95 |
| 10 | *GAL-CLN2* | 0.5x | 0.52G | Dirick 95 |
| 11 | *Multi-copy CLN2* | Viable, <1x | 0.81x | |
| 12 | *cln3Δ* | 1.8-2.7x | 2.2x | Dirick 95, Costanzo 04 |
| 13 | *CLN3-1* | 0.7x | 0.49x | Costanzo 04 |
| 14 | *GAL-CLN3* | 0.5x | 0.40G | Tyers 92 |
| 15 | *whi5Δ* | 0.6-0.7x | 0.82x | de Bruin 04, Costanzo 04 |
| 16 | *GAL-WHI5* | >1x | 1.09G | de Bruin 04, Costanzo 04 |
| 17 | *clb56Δ* | >1x | 1.11x | Schwob 93 |
| 18 | *Multi-copy CLB5* | CEN, Viable | 0.98x, 4 copies | |
| 19 | *GAL-CLB5* | Viable | 0.95G | Schwob 93 |
| 20 | *CLB5-dbΔ* | Viable | 1.04x | Wasch 02 |
| 21 | *GAL-CLB5dbΔ* | Inviable, DNA synthesis not advanced | T arrest, ORI advanced by 7.8’ | Schwob 94 |
| 22 | *triple-cln* | Inviable | G1 arrest ORI 248’ | Richardson 89 |
| 23 | *mbp1Δ* | 1.3x | 1.15x | Ferrezuelo 09, Koch 93 |
| 24 | *swi4Δ* | 1.3-1.5x | 1.42x | Wijnen 99, Wijnen 02 |
| 25 | *swi6Δ* | 2.4x | 2.25x | Wijnen 02 |
| 26 | *msn5Δ* | 1.4x | 1.34x | Queralt 03 |
| 27 | *sic1Δ* | <1x | 0.90x | Schneider 96 |
| 28 | *GAL-SIC1* | >1x | 1.09G | Nugro 94, Verma 97 |
| 29 | *GAL-SIC1dbΔ* | G1 arrest | G1 arrest | Verma 97 |
| 30 | *Multi-copy SIC1* | Viable | 1.10x | |
| 31 | *cdc6Δ* | <1x | 1.02x | Calzada 01 |
| 32 | *GAL-CDC6* | Viable | 1.06G | Archambault 03 |
| 33 | *Multi-copy CDC6* | Viable | 1.02x | |
| 34 | *ckiΔ* | Viable, <1x | 0.88x | Wasch 02 |
| 35 | *swi5Δ* | Viable | 0.98x | Toyn 97, Giaever 02 |
| 36 | *cdh1Δ* | Viable, <1x | 0.94x | Schwab 97, Wasch 02 |

| | | | | |
|---|---|---|---|---|
| 51 | *bck2Δ swi6Δ GAL-CLN3* | Prediction: Not rescued | Inviable | – |
| 52 | *bck2Δ swi6Δ GAL-CLN2* | Prediction: Rescued | 1.41G | – |
| 53 | *bck2Δ swi6Δ whi5Δ* | Inviable | G1 arrest | de Bruin 04 |
| 54 | *bck2Δ whi5Δ* | 0.85x | 0.91x | de Bruin 04, Costanzo 04 |
| 55 | *bck2Δ GAL-WHI5* | Viable, >*GAL-WHI5* | 1.49G | Costanzo 04 |
| 56 | *bck2Δ GAL-WHI5-12A* | Prediction: Viable, large | 1.72G | – |
| 57 | *GAL-BCK2 whi5Δ* | 0.5x | 0.57G | Costanzo 04 |
| 58 | *cln1Δ cln2Δ clb5Δ clb6Δ* | Inviable | G1 arrest | Schwob 93 |
| 59 | *cln1Δ cln2Δ cdh1Δ* | Viable | T arrest | Cross 02 |
| 60 | *cln1Δ cln2Δ GAL-CLN2 cdh1Δ* | Viable | 0.11G | Cross 02 |
| 61 | *cln1Δ cln2Δ sic1Δ* | Viable | 1.08x | Dirick 95 |
| 62 | *cln1Δ cln2Δ GAL-SIC1* | G1 arrest | G1 arrest | Cross 02 |
| 63 | *GAL-CLN2 GAL-SIC1* | Viable | 2.85G | Cross 02 |
| 64 | *GAL-CLN2 cdh1Δ GAL-SIC1* | Viable | 0.52G | Cross 02 |
| 65 | *cln2Δ cdh1Δ GAL-SIC1* | Inviable | G1 arrest | Cross 02 |
| 66 | *cln3Δ mbp1Δ* | Prediction: Inviable | 2.25x | Adames 15 |
| 67 | *cln3Δ mbp1Δ swi6Δ* | Prediction: Rescued, ~*swi6Δ* | 1.59x | Adames 15 |
| 68 | *cln3Δ mbp1Δ whi5Δ* | Prediction: Rescued | 1.23x | Adames 15 |
| 69 | *cln3Δ mbp1Δ mc-BCK2* | Prediction: Viable | 0.98x | Adames 15 |
| 70 | *cln3Δ mbp1Δwhi5Δ bck2Δ* | Prediction: Viable | 1.43x | Adames 15 |
| 71 | *cln3Δ swi4Δ* | Inviable | Inviable | Ferrezuelo 09 |
| 72 | *cln3Δ swi4Δ whi5Δ* | Prediction: Rescued | 2.36x | – |
| 73 | *cln3Δ swi4Δ GAL-BCK2* | Prediction: Rescued | 1.50G | – |
| 74 | *cln3Δ swi4Δ whi5Δ sic1Δ* | Prediction: Rescued | 1.73x | – |
| 75 | *cln3Δ swi4Δ whi5Δ GAL-BCK2* | Prediction: Rescued | 2.41G | – |
| 76 | *cln3Δ swi6Δ* | 2.4x, ~*swi6Δ* | 2.34x | Wijnen 02 |
| 77 | *CLN3-1 swi6Δ* | 2.4x, ~*swi6Δ* | 2.29x | Wijnen 02 |
| 78 | *cln3Δ whi5Δ* | 0.7-0.9x | 1.07x | de Bruin 04, Costanzo 04 |
| 79 | *CLN3-1 whi5Δ* | Viable, small | 0.36G | Costanzo 04 |
| 80 | *cln3Δ GAL-WHI5* | Inviable | 2.76G | Costanzo 04, Wagner 09 |
| 81 | *cln3Δ GAL-WHI5-12A* | Inviable | 2.79G | Costanzo 04 |
| 82 | *triple-cln GAL-CLN2* | Viable | 0.93G | Cross 91 |
| 83 | *triple-cln GAL-CLN3* | Viable | 1.36G | Cross 90 |
| 84 | *triple-cln sic1Δ* | >>1x | 1.84x | Tyers 96 |
| 85 | *triple-cln cdh1Δ* | T arrest | T arrest | Schwab 97 |
| 86 | *triple-cln mc-CLB5* | CEN, Viable | 2.58x | Epstein 92 |
| 87 | *triple-cln GAL-CLB5* | Viable | 1.81G | Schwob 93 |
| 88 | *triple-cln mc-BCK2* | Viable | 1.14x | Epstein 94 |
| 89 | *triple-cln GAL-CLB2* | G1 arrest | T arrest, ORI 148' | Amon 94 |
| 90 | *mbp1Δ swi4Δ* | Inviable | G1 arrest | Koch 93 |

| | | | | |
|---|---|---|---|---|
| 91 | *mbp1Δ swi4Δ* | Inviable | G1 arrest | Koch 93 |
| 92 | *mbp1Δ whi5Δ* | <1x | 0.92x | de Bruin 04 |
| 93 | *mbp1Δ GAL-WHI5* | Prediction: Viable | 1.13G | – |
| 94 | *mbp1Δ GAL-WHI5-12A* | Prediction: Viable | 1.22G | – |
| 95 | *mbp1Δ swi6Δ* | 2.4x, ~*swi6Δ* | 1.59x | Ferrezulo 09 |
| 96 | *swi4Δ swi6Δ* | Inviable | G1 arrest | Dirick 91 |
| 97 | *swi4Δ swi6Δ SWI6-SA4* | Rescued | 1.38x | Wijnen 02 |
| 98 | *swi4Δ swi6Δ GAL-CLB5* | Prediction: Viable, >1x | 1.87G | – |
| 99 | *swi4Δ swi6Δ GAL-CLN3* | Inviable | Inviable | – |
| 100 | *swi4Δ swi6Δ GAL-CLN2* | Rescued | 1.30G | – |
| 101 | *swi4Δ whi5Δ* | 1.3-1.5, ~*swi4Δ* | 1.19x | Jorgensen 02 |
| 102 | *swi4Δ GAL-WHI5* | Viable, ~*swi4Δ* | 1.82G | de Bruin 04 |
| 103 | *swi4Δ swi6Δ whi5Δ* | Prediction: Inviable | Inviable | – |
| 104 | *swi6Δ whi5Δ* | 2.4x, ~*swi6Δ* | 1.59x | Costanzo 04 |
| 105 | *swi6Δ GAL-WHI5* | Inviable | 1.07G | Costanzo 04 |
| 106 | *swi6Δ mc-WHI5* | Prediction: Viable | 1.60x | – |
| 107 | *msn5Δ swi4Δ* | Inviable | 1.79x | Queralt 06 |
| 108 | *msn5Δ swi6Δ* | Inviable | 1.59x | Queralt 06 |
| 109 | *cdh1Δ sic1Δ* | Inviable | T arrest | Schwab 97, Archambault 03 |
| 110 | *cdh1Δ cdc6Δ* | Viable, >1x | 0.96x | Calzada 01 |
| 111 | *cdh1Δ sic1Δ cdc6Δ* | Inviable | T arrest | Archambault 03 |
| 112 | *cdh1Δ swi5Δ* | Inviable | T arrest | Archambault 03 |
| 113 | *cdh1Δ swi5Δ GAL-SIC1* | Rescued | 1.06G | Archambault 03 |
| 114 | *GAL-CLB5 cdh1Δ* | Inviable*; Viable in model | 0.83G | Chen 04 |
| 115 | *GAL-CLB5 sic1Δ* | Inviable | Inviable, ORI not relicensed | Jacobson 00 |
| 116 | *CLB5-dbΔ sic1Δ* | Inviable, ORI not relicensed | Inviable, ORI not relicensed | Jacobson 00, Wasch 02 |
| ***Some essential exit mutants*** | | | | |
| 117 | *triple-cln apc-ts* | M arrest | M arrest | Iringer 97 |
| 118 | *cdh1Δ sic1Δ GALL-CDC20* | Viable, >1x | 0.97G | Cross 03 |
| 119 | *cdh1Δ sic1Δ cdc6Δ GALL-CDC20* | Rescued | 1.01G | Cross 03 |
| 120 | *CLB5-dbΔ pds1 Δ* | Viable | 1.15x | Wasch 02 |
| 121 | *CLB5-dbΔ pds1Δ cdc20Δ* | T arrest | T arrest | Wasch 02 |
| 122 | *cdc20Δ* | Inviable | M arrest | Lim 98 |
| 123 | *pds1Δ* | Viable | 1.08x | Yamamoto 96 |
| 124 | *tem1-ts* | Inviable | T arrest | Schirayama 94 |
| 125 | *cdc15Δ* | Inviable | T arrest | Jasperson 96 |
| 126 | *net1-ts* | Viable, >1x | 1.08x | Visintin 99 |
| 127 | *cdc14Δ* | Inviable | T arrest | Visintin 98 |
| 128 | *ppxΔ* | Viable | 1.27x | Wang 97 |
| 129 | *cdc20Δ clb5Δ* | M arrest | M arrest | Shirayama 98 |
| 130 | *cdc20Δ pds1Δ* | T arrest | T arrest | Shirayama 98 |
| 131 | *cdc20Δ clb5Δ pds1Δ* | Viable, >1x | 1.39x | Shirayama 98 |
| 132 | *cdc20Δ GAL-ESP1* | T arrest, Cdc14 out | T arrest, Cdc14 out | Uhlmann 99 |
| 133 | *APC-A* | Viable | 1.20x | Rudner 00, Cross 03 |
| 134 | *APC-A cdh1Δ* | T arrest | T arrest | Cross 03 |
| 135 | *APC-A cdh1Δ in GAL* | T arrest | T arrest | Cross 03 |
| 136 | *APC-A cdh1Δ mc-SIC1* | Rescued | 1.41x | Cross 03 |
| 137 | *APC-A cdh1Δ GAL-SIC1* | Rescued | 1.14G | Cross 03 |
| 138 | *APC-A cdh1Δ mc-CDC6* | Rescued | 1.16x | Cross 03 |
| 139 | *APC-A cdh1Δ GAL-CDC6* | Rescued | 1.11G | Cross 03 |
| 140 | *APC-A cdh1Δ mc-CDC20* | Rescued | 0.84x | Cross 03 |
| 141 | *APC-A sic1Δ* | Rescued | 1.06x | Cross 03 |
| 142 | *APC-A GAL-CLB2* | T arrest | T arrest | Cross 03 |

* Fred Cross: Although *GAL-CLB5 cdh1Δ* is ultimately inviable or extremely slow-growing on galactose, these mutants are not associated with any obvious problems in a short-term experiment. These cells go through several doublings, probably without much difficulty, on galactose medium, and they remain reasonably viable when returned to glucose, like *GAL-CLB5* cells. So, it is not reasonable to expect the mathematical model to predict inviability of *GAL-CLB5 cdh1Δ* mutants. They should probably look viable.

### Supplementary Text

#### List of Abbreviations

DNA – Deoxy-Ribo Nucleic Acid
S, M, G1, G2 – Phases of cell cycle: DNA Synthesis, Mitosis, Gap phases 1, 2
CDK – Cyclin Dependent Kinase
CKI – Cyclin dependent Kinase Inhibitor
SBF, MBF – Scb- and Mcb-element Binding Factor
R point – Restriction Point
BYCC – Chen et al, 2004, Budding Yeast Cell Cycle model
START-BYCC – New START model incorporated with BYCC
ORI – species in our model, which is a marker for Origin of replication
WT – Wild Type
BUD – species in our model, which is a marker for Budding
SBFB – SBF Bound to promoter
WSB – Whi5-SBF Bound to promoter
Swi4B – Swi4 dimer Bound to promoter
SBFa1-a5 – explained in legend of Figure 4E.

#### Equations, Parameters and Initial Conditions

(will be reformatted and inserted from the latest PET file)
https://github.com/jravilab/start-bycc

### License

License Agreement from ASM J Bacteriology (CCC marketplace) to reuse and adapt Fig. 6 from Lord and Wheals (1980) – requested for Fig. 5 in main manuscript.

CCC | Marketplace

This is a License Agreement between Janani Ravi ("User") and Copyright Clearance Center, Inc. ("CCC") on behalf of the Rightsholder identified in the order details below. The license consists of the order details, the Marketplace Permissions General Terms and Conditions below, and any Rightsholder Terms and Conditions which are included below.
All payments must be made in full to CCC in accordance with the Marketplace Permissions General Terms and Conditions below.

| | | | |
|---|---|---|---|
| **Order Date** | 01-Nov-2023 | **Type of Use** | Republish in other published product |
| **Order License ID** | 1412393-1 | **Publisher** | AMERICAN SOCIETY FOR MICROBIOLOGY |
| **ISSN** | 1098-5530 | **Portion** | Chart/graph/table/figure |

#### LICENSED CONTENT

| | | | |
|---|---|---|---|
| **Publication Title** | Journal of bacteriology : JB | **Publication Type** | e-Journal |
| **Article Title** | Asymmetrical division of Saccharomyces cerevisiae. | **Start Page** | 808 |
| **Author/Editor** | American Society for Microbiology. | **End Page** | 818 |
| **Date** | 01/01/1916 | **Issue** | 3 |
| **Language** | English | **Volume** | 142 |
| **Country** | United States of America | **URL** | https://journals.asm.org/journal/jb |
| **Rightsholder** | American Society for Microbiology - Journals | | |

#### REQUEST DETAILS

| | | | |
|---|---|---|---|
| **Portion Type** | Chart/graph/table/figure | **Distribution** | Worldwide |
| **Number of Charts / Graphs / Tables / Figures Requested** | 1 | **Translation** | Original language of publication |
| **Format (select all that apply)** | Electronic | **Copies for the Disabled?** | Yes |
| **Who Will Republish the Content?** | Publisher, not-for-profit | **Minor Editing Privileges?** | Yes |
| **Duration of Use** | Life of current edition | **Incidental Promotional Use?** | No |
| **Lifetime Unit Quantity** | Up to 499 | **Currency** | USD |
| **Rights Requested** | Main product | | |

#### NEW WORK DETAILS

| | | | |
|---|---|---|---|
| **Title** | Modeling the START transition in the budding yeast cell cycle | **Produced by** | unknown. first submitting as preprint to bioRxiv. |
| **Author** | Janani Ravi, Kewalin Samart, Jason Zwolak | **Expected Publication Date** | 2023-11-01 |

#### ADDITIONAL DETAILS

| | | | |
|---|---|---|---|
| **Order Reference Number** | N/A | **The Requesting Person / Organization to Appear on the License** | Janani Ravi |

#### REQUESTED CONTENT DETAILS

| | | | |
|---|---|---|---|
| **Title, Description or Numeric Reference of the Portion(s)** | Fig 6 | **Title of the Article / Chapter the Portion Is From** | Asymmetrical division of Saccharomyces cerevisiae. |
| **Editor of Portion(s)** | Lord, P G; Wheals, A E | **Author of Portion(s)** | Lord, P G; Wheals, A E |
| **Volume / Edition** | 142 | **Issue, if Republishing an Article From a Serial** | 3 |
| **Page or Page Range of Portion** | 808-818 | **Publication Date of Portion** | 1980-05-31 |

#### References

1. Costanzo M, Nishikawa JL, Tang X, et al. CDK Activity Antagonizes Whi5, an Inhibitor of G1/S Transcription in Yeast. *Cell*. 2004;117(7):899-913. doi:10.1016/j.cell.2004.05.024
2. de Bruin RAM, McDonald WH, Kalashnikova TI, Yates J, Wittenberg C. Cln3 Activates G1-Specific Transcription via Phosphorylation of the SBF Bound Repressor Whi5. *Cell*. 2004;117(7):887-898. doi:10.1016/j.cell.2004.05.025
3. Ghaemmaghami S, Huh WK, Bower K, et al. Global analysis of protein expression in yeast. *Nature*. 2003;425(6959):737-741. doi:10.1038/nature02046
4. Pramila T. The Forkhead transcription factor Hcm1 regulates chromosome segregation genes and fills the S-phase gap in the transcriptional circuitry of the cell cycle. *Genes Dev*. 2006;20(16):2266-2278. doi:10.1101/gad.1450606
5. Wagner A, Grillitsch K, Leitner E, Daum G. Mobilization of steryl esters from lipid particles of the yeast Saccharomyces cerevisiae. *Biochim Biophys Acta BBA - Mol Cell Biol Lipids*. 2009;1791(2):118-124. doi:10.1016/j.bbalip.2008.11.004
6. Taberner FJ, Quilis I, Igual JC. Spatial regulation of the Start repressor Whi5. *Cell Cycle*. 2009;8(18):3013-3022. doi:10.4161/cc.8.18.9621
7. Talia SD, Skotheim JM, Bean JM, Siggia ED, Cross FR. The effects of molecular noise and size control on variability in the budding yeast cell cycle. *Nature*. 2007;448(7156):947-951. doi:10.1038/nature06072
8. Koch C, Moll T, Neuberg M, Ahorn H, Nasmyth K. A role for the transcription factors Mbp1 and Swi4 in progression from G1 to S phase. *Science*. 1993;261(5128):1551-1557. doi:10.1126/science.8372350
9. Moll T, Tebb G, Surana U, Robitsch H, Nasmyth K. The role of phosphorylation and the CDC28 protein kinase in cell cycle-regulated nuclear import of the S. cerevisiae transcription factor SWI5. *Cell*. 1991;66(4):743-758. doi:10.1016/0092-8674(91)90118-i
10. Wijnen H, Landman A, Futcher B. The G $_1$ Cyclin Cln3 Promotes Cell Cycle Entry via the Transcription Factor Swi6. *Mol Cell Biol*. 2002;22(12):4402-4418. doi:10.1128/MCB.22.12.4402-4418.2002
11. Sidorova JM, Mikesell GE, Breeden LL. Cell cycle-regulated phosphorylation of Swi6 controls its nuclear localization. *Mol Biol Cell*. 1995;6(12):1641-1658. doi:10.1091/mbc.6.12.1641
12. Geymonat M, Spanos A, Wells GP, Smerdon SJ, Sedgwick SG. Clb6/Cdc28 and Cdc14 Regulate Phosphorylation Status and Cellular Localization of Swi6. *Mol Cell Biol*. 2004;24(6):2277-2285. doi:10.1128/MCB.24.6.2277-2285.2004
13. Queralt E, Igual JC. Cell Cycle Activation of the Swi6p Transcription Factor Is Linked to Nucleocytoplasmic Shuttling. *Mol Cell Biol*. 2003;23(9):3126-3140. doi:10.1128/MCB.23.9.3126-3140.2003
14. Siegmund RF, Nasmyth KA. The Saccharomyces cerevisiae Start-specific transcription factor Swi4 interacts through the ankyrin repeats with the mitotic Clb2/Cdc28 kinase and through its conserved carboxy terminus with Swi6. *Mol Cell Biol*. 1996;16(6):2647-2655. doi:10.1128/MCB.16.6.2647
15. Baetz K, Andrews B. Regulation of Cell Cycle Transcription Factor Swi4 through Auto-Inhibition of DNA Binding. *Mol Cell Biol*. 1999;19(10):6729-6741. doi:10.1128/MCB.19.10.6729
16. Dirick L, Nasmyth K. Positive feedback in the activation of G1 cyclins in yeast. *Nature*. 1991;351(6329):754-757. doi:10.1038/351754a0

17. de Bruin RAM, Kalashnikova TI, Aslanian A, et al. DNA replication checkpoint promotes G1-S transcription by inactivating the MBF repressor Nrm1. *Proc Natl Acad Sci*. 2008;105(32):11230-11235. doi:10.1073/pnas.0801106105
18. Vergés E, Colomina N, Garí E, Gallego C, Aldea M. Cyclin Cln3 Is Retained at the ER and Released by the J Chaperone Ydj1 in Late G1 to Trigger Cell Cycle Entry. *Mol Cell*. 2007;26(5):649-662. doi:10.1016/j.molcel.2007.04.023
19. Wijnen H, Futcher B. Genetic Analysis of the Shared Role of CLN3 and BCK2 at the G1-S Transition in Saccharomyces cerevisiae. *Genetics*. 1999;153(3):1131-1143. doi:10.1093/genetics/153.3.1131
